# Astrocyte cell-surface proteomics identified CD44 as an OPN/SPP1 receptor regulating lipid metabolism and glial crosstalk in Alzheimer’s disease

**DOI:** 10.64898/2025.12.05.692670

**Authors:** Omar Peña-Ramos, Manasee Gedam, Xue Zhang, Shuo Wang, Chuangye Qi, Tzu-Hsuan Huang, Chetan Singh Rajpurohit, Mikayla Dunlap, Yu Deng, Niccole Auld, Sung Yun Jung, Hongjie Li, Jiefu Li, Liqun Luo, Junmin Peng, Chonghui Cheng, Hui Zheng

## Abstract

The development of β-amyloid (Aβ) pathology in Alzheimer’s disease (AD) is accompanied by profound changes in astrocytes and microglia. How these responses are orchestrated by cell surface proteins, key mediators of cell-cell communication, remain unclear. Using in situ astrocyte cell-surface proteome profiling in 5xFAD mice, we identified a set of dysregulated surface proteins induced by Aβ pathology, including CD44. CD44 was selectively upregulated in plaque-adjacent astrocytes and interacted with osteopontin (OPN), encoded by the disease-associated microglia gene *Spp1*, to promote lipid accumulation, and this effect is γ-secretase dependent. Astrocytic CD44 in turn regulated *Spp1* expression and microglial activity. Conditional deletion of *Cd44* in adult astrocytes of 5xFAD mice attenuated glial reactivity, reduced Aβ pathology, and improved cognition. These findings define a plaque-proximal OPN–CD44 axis that controls astrocyte lipid metabolism and glial activity, positioning CD44 as a surface-accessible therapeutic target in AD.

## Introduction

Astrocytes are essential for brain homeostasis, providing trophic and metabolic support to neurons, regulating synaptic activity, maintaining the blood-brain barrier, and modulating immune and inflammatory pathways^1,2^. These diverse functions are mediated in part by cell-surface proteins (CSPs), which sense extracellular cues, transduce signals into intracellular responses, and orchestrate cell-cell communication. During the development of Aβ pathology in AD, astrocytes transition to a reactive state characterized by morphological and transcriptional remodeling^3,4^. However, it is still unclear how these changes are reflected at the protein level especially within the CSP repertoire, because RNA and protein levels correlate only modestly^5,6^. Defining astrocytic CSPs dysregulated in AD is therefore critical to understanding their contribution to disease progression and may represent therapeutically tractable targets given their surface accessibility.

To address this gap, we applied in situ cell-surface proteome extraction by extracellular labeling (iPEEL)^7–9^, which uses a mouse line expressing membrane-anchored horseradish peroxidase (HRP) in a Cre-dependent manner. This system enables proximity labeling and mass spectrometry (MS)-based identification of CSPs and their extracellular interactors directly in intact brain tissue. Using the GFAP-Cre, we profiled astrocytic CSPs in the 5xFAD model of amyloidosis and identified a set of surface receptors and soluble proteins that were up or downregulated in response to Aβ pathology, including the type I transmembrane protein CD44.

CD44 is a cell adhesion molecule that acts as a receptor or co-receptor in diverse biological processes, including cell proliferation, migration, metabolism, and phagocytosis^10–12^. Although its canonical ligand is hyaluronic acid (HA), a major extracellular matrix component, CD44 also binds other ligands, most notably osteopontin (OPN)^13^, which is encoded by the disease-associated microglia (DAM) gene *Spp1*^14^. CD44 signaling activates downstream pathways such as STAT3, a central mediator of astrocyte reactivity^15–18^. Of interest, CD44 undergoes ectodomain shedding followed by γ-secretase processing similar to APP and Notch, and the released CD44 intracellular domain (CD44ICD) has been shown to translocate to the nucleus to regulate gene expression^15,19–26^.

Within the CNS, CD44 expression is largely restricted to astrocytes and is upregulated in AD brains as well as in pan-reactive astrocytes induced by microglia-derived TNFα, IL-1α and C1q (TIC)^27–29^. In a large-scale AD proteomic and coexpression network analysis, CD44 emerged as a hub protein within an astrocyte/microglia module highly enriched in AD brains, with expression positively correlating with AD neuropathology and cognitive decline^30^. Notably, CD44 levels are elevated in the cerebrospinal fluid of asymptomatic individuals with AD, underscoring its potential as an early biomarker^30,31^.

Here, we demonstrate that CD44 defines a subpopulation of astrocytes that promotes lipid accumulation in an OPN- and γ-secretase-dependent but HA- and STAT3-independent manner. Genetic deletion of *Cd44* in adult astrocytes attenuated both astrocyte and microglial reactivity, reduced plaque burden, and improved cognitive performance in 5xFAD mice.

## Results

### In situ cell-surface proteome profiling of astrocytes in 5xFAD mice

We used iPEEL to identify Aβ-induced CSPs changes in astrocytes **(Figure 1A)**. Specifically, we crossed HRP^loxP-STOP-loxP^ (Ctrl) mice with GFAP^Cre^ mice to express HA-tagged, membrane-tethered HRP in astrocytes (herein referred to as HRP), and further introduced these alleles into the 5xFAD background (HRP;5xFAD). Only female GFAP^Cre^ mice were used for breeding to a prevent germline recombination caused by Cre activity in male germ cells^32,33^. We validated astrocyte-specific expression by HA and GFAP co-staining **(Figure 1B)**. Immunoblotting confirmed HRP expression in hippocampus and cortex of HRP mice, but not in Cre-negative controls **(Figure 1C)**. Incubation of fresh brain sections with hydrogen peroxide (H₂O₂) and the membrane-impermeable substrate Biotin-xx-Phenol (BxxP) enabled HRP-dependent covalent biotinylation of proximal membrane and extracellular proteins in live tissue. Neutravidin staining showed robust biotin labeling throughout treated brain sections **(Figure 1D)**. Silver staining and streptavidin pulldown further confirmed efficient enrichment of biotinylated proteins from HRP samples relative to controls **(Figure 1E)**. These results demonstrate low background and robust astrocyte CSP labeling.

**Figure 1.**
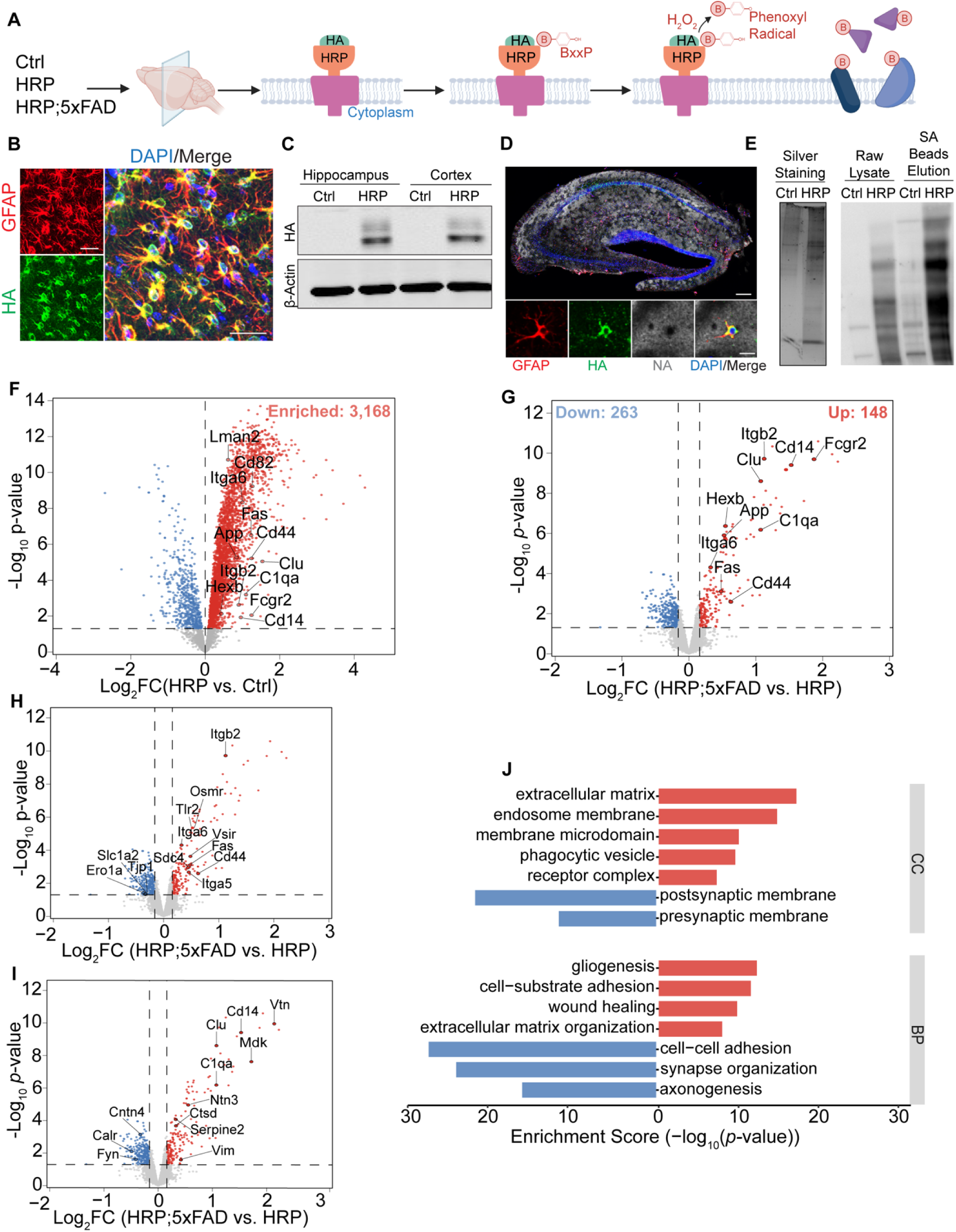
Astrocyte cell surface proteome analysis of mouse brains using iPEEL. **(A)** Schematic iPEEL performed on 300 μm sagittal brain sections from HRP^loxP-STOP-loxP^ (Ctrl), HRP HRP^loxP-STOP-loxP^;GFAP^Cre^ (HRP), and HRP;5xFAD mice at 4.5 months of age. In the presence of H2O2 and the membrane impermeable dye Biotin-xx-Phenol (BxxP), the extracellular HA-tagged HRP on astrocyte membrane generates phenoxyl radicals that covalently biotinylate nearby membrane and extracellular proteins. **(B)** Representative hippocampal images of a 2-month-old HRP mouse showing astrocyte expression of HRP (HA, green) colocalizing with GFAP (red). DAPI (blue) marks nuclei in the merged image. Scale bar: 20 μm. **(C)** Immunoblot of hippocampal and cortical lysates from Ctrl and HRP mice using anti-HA and loading control anti-β-actin antibodies. **(D)** Representative iPEEL labeling across a hippocampal section from 2-month-old HRP mice stained with NeutrAvidin (NA, grey), GFAP (red), HA (green), and DAPI (blue). Inset, single astrocyte showing surface labelling. Scale bars: 200 μm (overview) and 20 μm (inset). **(E)** Silver staining and streptavidin blotting of streptavidin (SA)-beads-enriched proteins and corresponding lysates. **(F)** Volcano plot of biotinylated CSPs in HRP versus Ctrl samples. Enriched (red) defined by log2FC > 0 and *p* < 0.05. Dashed lines indicate threshold. Selected proteins are labeled. **(G-I)** Volcano plots of differentially expressed CSPs (G), which are further stratified by receptors (H) or ligands (I) in HRP;5xFAD versus HRP with selected upregulated (red) and downregulated (blue) CSPs highlighted. Cut off: |log2 fold-change| > 1 SD (SD = 0.16) and *p* < 0.05. Y-axis shows -log10 *p*-value. Enrichment tested by hypergeometric test (*p* < 0.05). **(J)** GO term enrichment analysis of upregulated (red) and downregulated (blue) CSPs comparing HRP;5xFAD vs HRP for Cellular Compartment (CC) and Biological Processes (BP) terms.

We next performed multiplexed tandem mass tag (TMT)-MS on streptavidin-enriched hippocampal samples from Ctrl, HRP and 5xFAD;HRP mice at 4.5-4.8 months of age (3M +3F per group)^34^. Following quality control by silver staining, data normalization, and principal component analysis (PCA) (**Figure S1),** comparative analysis of HRP versus non-HRP samples identified >3000 enriched biotinylated proteins **(Figure 1F)**. These proteins served as the reference set for differential expression analysis between 5xFAD;HRP and HRP mice, revealing 263 downregulated and 148 upregulated proteins **(Figure 1G and Table S1)**. This dataset included known Aβ-associated proteins like App, Apoe, Clu, and C1qa, validating the iPEEL approach. Cross-referencing the TMT datasets with the curated ligand-receptor interactions from the Omnipath database^35^ generated a curated list of differentially expressed receptors **(Figure 1H)** and ligands **(Figure 1I)**. Gene ontology (GO) analysis showed enrichment in extracellular matrix, membrane microdomains, receptor complex, phagocytic vesicles, and synaptic compartments, with biological processes including glycogenesis, wound healing, cell adhesion and synapse organization **(Figure 1J)**. Together, these findings establish iPEEL as a robust platform for identifying astrocyte CSPs and their Aβ-induced remodeling.

### CD44 defines a subset of astrocytes associated with Aβ pathology in mouse and human brains

CD44 emerged as a significantly upregulated cell surface receptor in our iPEEL profiling (**Figure 1H**). To validate this finding, we performed immunostaining in 6-month-old 5xFAD and 9-month-old APP^NL-G-F^ knock-in mouse brains, which revealed robust CD44 enrichment in S100β^+^ astrocytes located near methoxy-X04 (X04) labeled plaques, whereas distal astrocytes and wild-type controls showed minimal CD44 staining (**Figure 2A**). This plaque-associated pattern is consistent with observations in human AD brain tissue^29^. Western blot analysis further confirmed increased CD44 abundance in hippocampal lysates of 5xFAD mice compared to wild-type controls (**Figure S2**).

**Figure 2.**
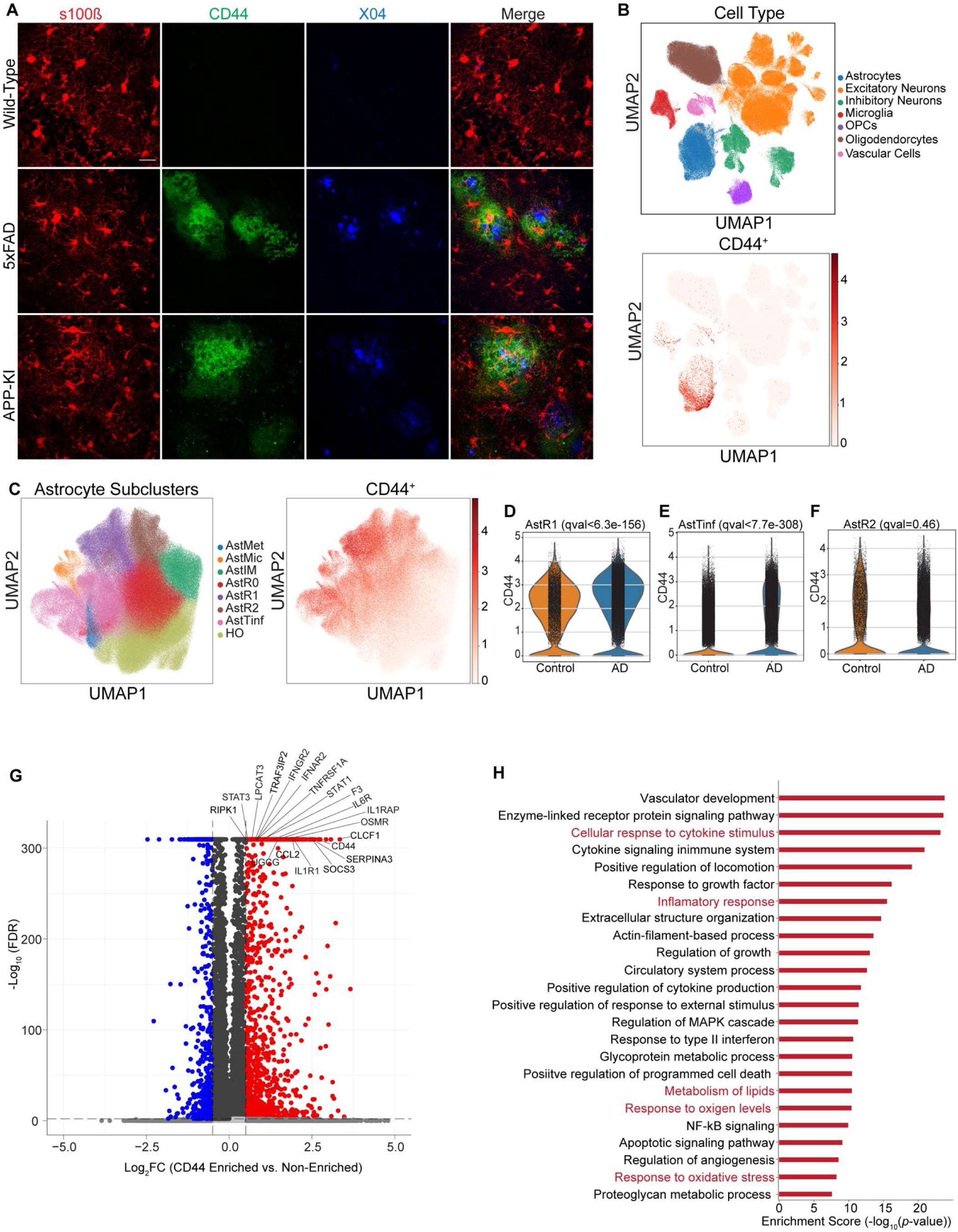
CD44 is upregulated in a subset of astrocytes close to Aβ plaques exhibiting dysregulated immune and metabolic profiles. **(A**) Representative images of the hippocampus from 6-month-old wild-type (WT) and 5xFAD, and 9-month-old APP^NL-G-F^ knock-in (APP-KI) mice showing selective CD44 (green) staining in S100β^+^ astrocytes (red) near methoxy-X04^+^ (X04, blue) plaques. Scale bar: 20 µm. **(B)** UMAP embedding of snRNA-seq from human prefrontal cortex (n=238,706 nuclei from 24 donors) annotated by major cell types showing enriched *CD44* expression in astrocytes. **(C)** UMAP plots of astrocyte nuclei (n=545,288) showing *CD44* expression across astrocyte subclusters. **(D-F)** Violin plots showing significantly elevated *CD44* expression in AD compared to control within AstR1 **(D)**, AstTinf **(E)**, but not AstrR2 **(F)** subclusters. **(G)** Volcano plot showing differentially expressed genes in *CD44* enriched vs. non-enriched astrocytes, with upregulated genes in red and downregulated in blue (FDR < 0.05 and log2(fold change) > 0.25). **(H)** GO term enrichment analysis of upregulated pathways in CD44 enriched astrocytes, highlighting pathways related to inflammatory response, immune activation, reactive oxygen species (ROS) production, and lipid metabolism.

To establish cellular specificity and relevance to human AD, we analyzed a single-nucleus RNA-sequencing (snRNA-seq) dataset from the human prefrontal cortex of 24 donors^27^. *CD44* transcripts were highly enriched in astrocytes relative to other major cell types and exhibited marked heterogeneity across astrocyte populations (**Figure 2B**). To investigate this heterogeneity in greater depth, we analyzed a second snRNA-seq dataset enriched for astrocyte nuclei across multiple brain regions^29^. After cell type annotation and normalization across sex, diagnosis, brain region, pathology stage and donor age (**Figure S3**), we examined *CD44* expression across the eight astrocyte subclusters defined by Serrano-Pozo et al. (**Figure 2C**). *CD44* expression was restricted to three astrocyte subclusters, AstR1, AstR2, and AstTinf, and was significantly upregulated in AstR1 and AstTinf, but not in AstR2, in AD relative to controls (**Figures 2D-2F**).

To define the molecular signatures of *CD44^+^* astrocytes, we compared differentially expressed genes (DEGs) in *CD44^+^* versus *CD44^-^* astrocytes (**Figures 2G and 2H**). We identified 131 DEGs, including 89 significantly upregulated genes largely corresponding to immune and inflammatory pathway markers (**Figure 2G**). GO term analysis revealed enrichment of pathways related to inflammation, lipid metabolism, and oxidative stress in *CD44^+^* astrocytes (**Figure 2H**).

Collectively, these findings demonstrate that CD44 marks a subset of astrocytes selectively enriched around Aβ plaques^29,36^, and define *CD44^+^* astrocytes as a molecularly distinct population characterized by upregulated immune and lipid metabolic programs.

### Positive correlation between CD44 and BODIPY^+^ lipids in sorted astrocytes

Given the dysregulated lipid and immune pathways in CD44^+^ astrocytes, we next examined whether astrocytic CD44 is associated with increased lipid accumulation in vivo. To this end, we developed a flow cytometry strategy to quantify lipid content in freshly isolated astrocytes using BODIPY fluorescence (**Figures 3A and S4**). Single cell suspensions were prepared from 12-month-old WT and 5xFAD mouse brains, and astrocytes were identified via ACSA2^+^ labelling followed by sorting into CD44^+^ and CD44^-^populations. Using BODIPY to measure neutral lipid content, we found that nearly all CD44^+^ cells were BODIPY^+^ whereas only 25% of the CD44^-^ cells were BODIPY^+^ (**Figure 3B**). This pattern was observed in both WT and 5xFAD mice; however, further gating based on BODIPY intensity revealed a significant increase in the BODIPY-high population in 5xFAD astrocytes compared with WT controls, the latter showed a corresponding shift toward the BODIPY-low population (**Figures 3C and 3D**). These findings establish a strong positive correlation between CD44 expressions and lipid accumulation in astrocytes, both of which were elevated in 5xFAD mice.

**Figure 3.**
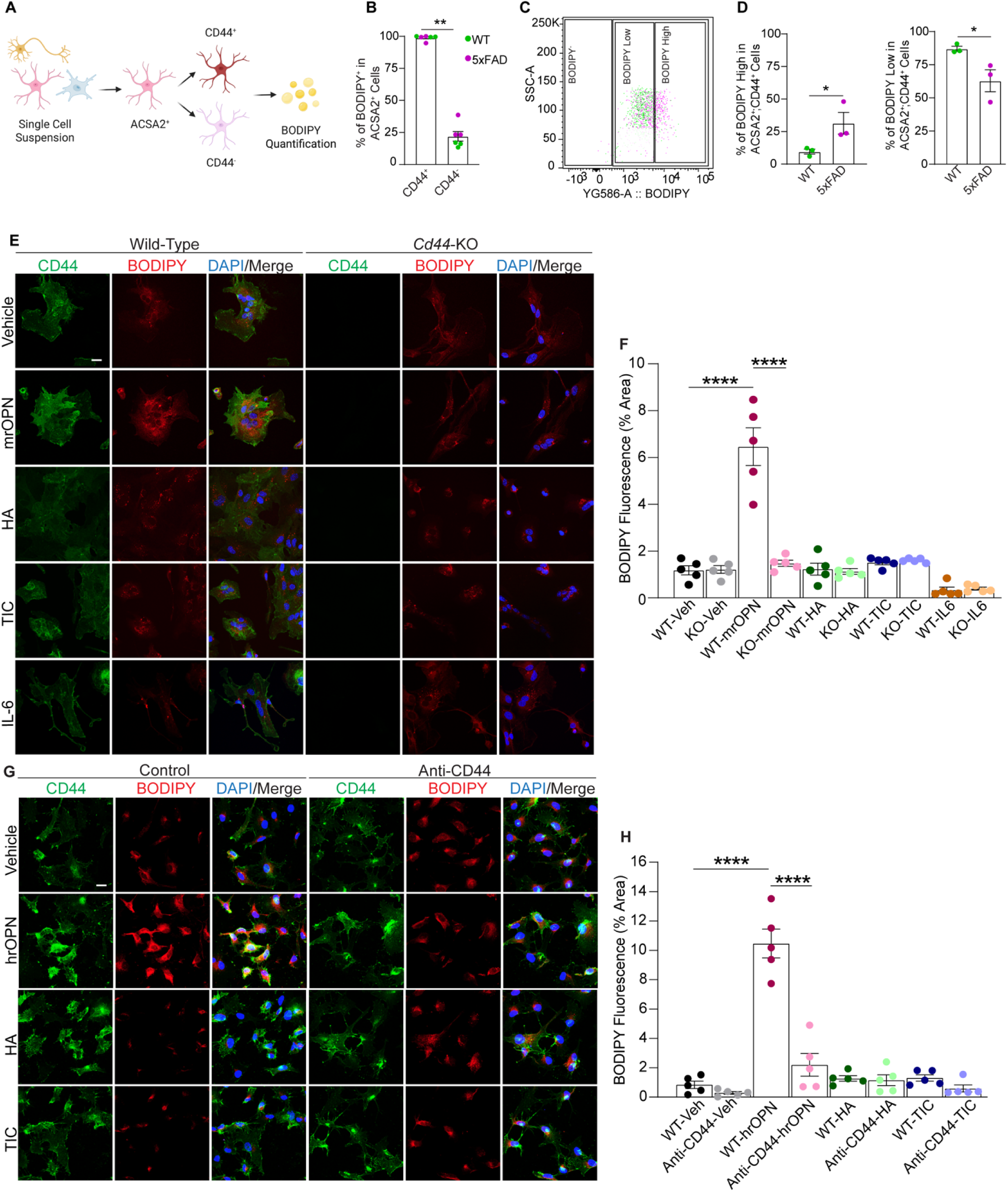
OPN-CD44 interaction induces lipid accumulation. **(A)** Schematic outlining the flow cytometry strategy to isolate ASCA2^+^ astrocytes from mouse brain single cell suspension of 12-month-old WT and 5xFAD mice followed by sorting into CD44^+^ and CD44^-^populations for BODIPY-fluorescence based lipid quantification. **(B)** Quantification of BODIPY+ astrocytes within CD44^+^ and CD44^-^ populations. **(C)** Flow cytometry dot plot showing the distribution of BODIPY-, BODIPY low, and BODIPY high ACSA2^+^ astrocytes populations in WT and 5xFAD mice. **(D)** Quantification of BODIPY high (left) and BODIPY low (right) fractions within ACSA2^+^;CD44^+^ astrocytes in WT and 5xFAD mice. Each dot represents one animal (n=3 mice per genotype/experiment). **(E)** Representative images of primary mouse astrocytes from wild-type and *Cd44-*KO mice treated with vehicle, mrOPN, HA, TIC, or IL-6 and stained for CD44, BODIPY, and DAPI. **(F)** Quantification of percentage of BODIPY^+^ area. **(G)** Representative images of human-iPSC-derived astrocytes treated with vehicle, hrOPN, HA or TIC, with or without the anti-CD44 blocking antibody and stained for CD44, BODIPY, and DAPI. **(H)** Quantification of percentage of BODIPY^+^ area. Each dot represents the average of six images collected from three independent coverslips. Data is presented as mean ± SEM. Statistical analyses were performed using 2-sided Student’s *t* test (B and D) or One-way ANOVA with Tukey’s multiple comparison (F and H). \*\**p* < 0.01; \*\*\*\**p* < 0.0001. Scale bars: 20 µm.

CD44 interacts with multiple extracellular ligands, particularly hyaluronic acid (HA), synthesized by the hyaluronan synthases HAS1–3, and OPN, encoded by the DAM gene *SPP1*. Correlation analysis using a human parahippocampal gyrus RNA-seq dataset^37^ revealed a strong positive correlation between *CD44* and *SPP1*, whereas no positive correlation was observed between *CD44* and *HAS1–3* (**Figures S5A-S5D**). Immunostaining of HA using an HA-binding protein antibody further showed no appreciable differences between wild-type and 5xFAD brains (**Figure S5E**). Together, these data support a potential interaction between OPN and astrocytic CD44 in AD.

### An OPN-CD44-γ-secretase axis driving astrocyte lipid accumulation

The OPN-CD44 signaling has been implicated in regulating inflammation and lipid metabolism in peripheral tissues^25,38,39^. We therefore tested whether CD44-associated lipid phenotype is OPN dependent. We used primary mouse astrocyte cultures and treated them with mouse recombinant OPN (mrOPN). To evaluate the OPN specificity, we also included treatments with HA, the cytokine cocktail TNFα, IL-1α and C1q (TIC) known to induce neurotoxic astrocytes^28^, and IL-6, an activator of the JAK/STAT3 pathway implicated in astrocyte reactivity and a potential downstream target of CD44^40^. mrOPN robustly increased BODIPY fluorescence in WT astrocytes, whereas HA, TIC, and IL-6 failed to induce this phenotype (**Figures 3E and 3F**). Notably, the OPN-induced lipid accumulation was blunted in *Cd44-*KO astrocytes, demonstrating that this effect is CD44 dependent **(Figures 3E and 3F**). Further supporting a STAT3 independent mechanism, IL-6 induced pSTAT3 nuclear localization in both WT and *Cd44*-KO cells, while OPN had no effect (**Figure S6**).

To determine whether OPN-CD44-mediated lipid accumulation is conserved in humans, we generated human iPSC-derived astrocytes. As in mouse cultures, human recombinant OPN (hrOPN), but not HA or TIC, significantly induced BODIPY^+^ lipid levels, and this effect was blocked by a CD44 neutralizing antibody, confirming CD44 dependency (**Figures 3G and 3H**). These findings demonstrate that OPN-CD44 signaling drives astrocyte lipid accumulation across species and in an STAT3 independent manner.

CD44 is a γ-secretase substrate. The OPN-CD44 signaling has been reported to promote glioma growth through γ-secretase processing and CD44 intracellular domain (CD44ICD) production^24^. We thus tested the requirement of CD44 processing and signaling in lipid accumulation dynamics using γ-secretase inhibitors DAPT and Compound E.

Using APP as a bona fide γ-secretase substrate, we verified that both DAPT and Compound E treatments led to accumulation of APP C-terminal fragment (CTF) in human iPSC-derived neurons^41^ and astrocytes (**Figure S7A**), indicating effective γ-secretase inhibition. We then treated primary mouse astrocytes (**Figures 4A and 4B**) and human iPSC-derived astrocytes (**Figures 4C and 4D**) with OPN and γ-secretase inhibitor alone or combined. As expected, OPN significantly increased BODIPY^+^ lipid accumulation in both cultures, and this effect was abolished by co-treatment with DAPT **(Figures 4A-4D)** and Compound E (**Figures S7B and 7C**), indicating that CD44ICD is required for OPN-induced lipid accumulation. DAPT alone did not alter lipid levels, suggesting that γ-secretase inhibition specifically blocks OPN-CD44 signaling rather than basal lipid homeostasis.

**Figure 4.**
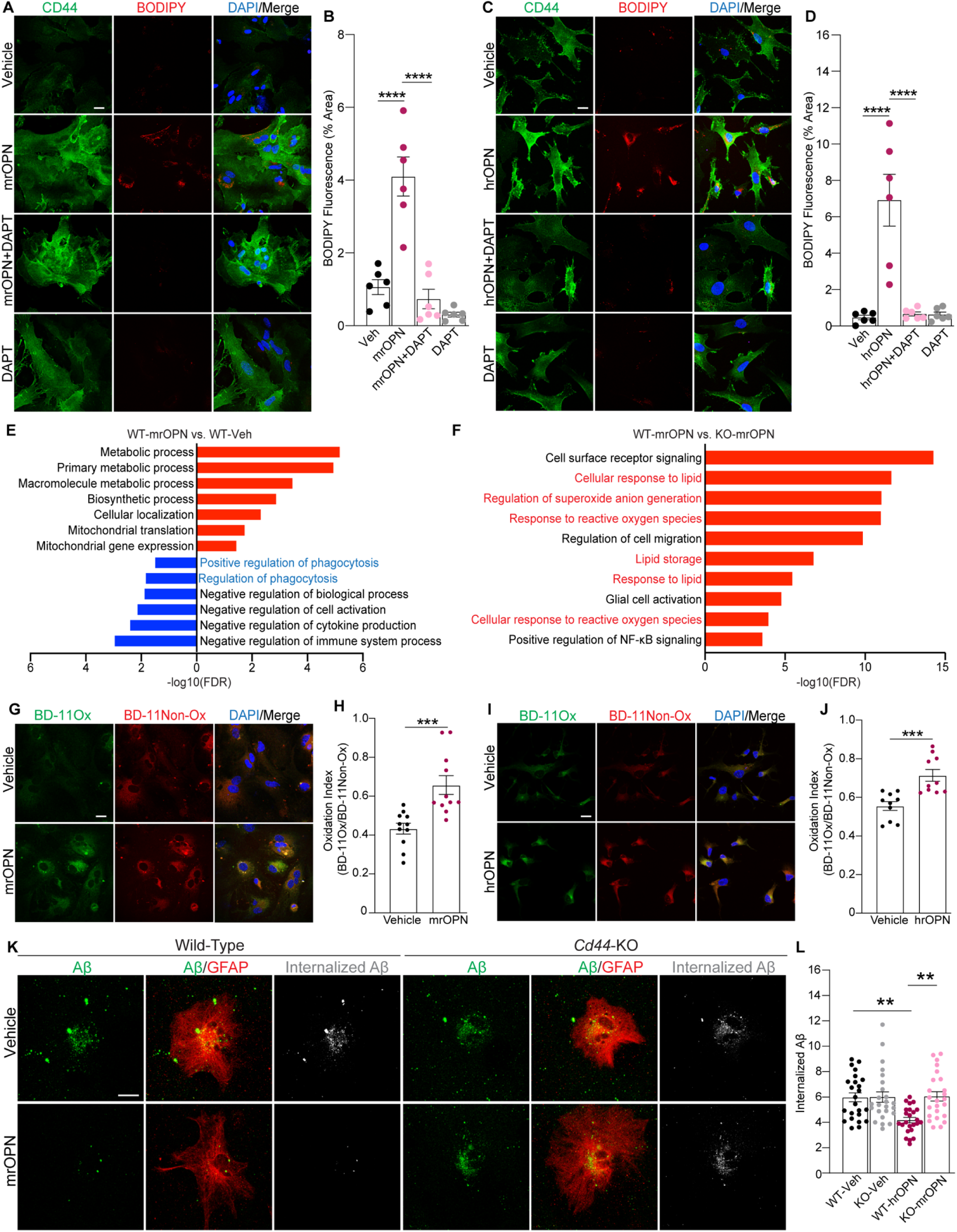
Mechanistic and functional characterization of the OPN-CD44 pathway. **(A)** Representative confocal images of primary mouse astrocytes treated with vehicle, mrOPN, mrOPN + DAPT, or DAPT alone, and stained for CD44 and BODIPY. **(B)** Quantification of BODIPY-positive area (% area). Each dot represents the average of six images collected from three independent coverslips. **(C)** Representative confocal images of human iPSC-derived astrocytes treated as in (A). **(D)** Quantification of BODIPY-positive area in human iPSC-derived astrocytes. Each dot represents one confocal image. **(E)** GO term enrichment analysis of differentially expressed proteins in WT astrocytes treated with mrOPN vs untreated controls. Red bars indicate upregulated pathways; blue bars downregulated bars, with phagocytosis-related pathways highlighted. **(F)** GO term enrichment analysis comparing mrOPN-treated WT and *Cd44*-KO astrocytes highlighting upregulation of pathways associated with lipid metabolism and ROS generation (red). **(G,I)** Representative images of primary mouse astrocytes and human iPSC-derive astrocytes treated with vehicle or recombinant mouse/human OPN stained with BODIPY C11 (green oxidative, red non-oxidative) and DAPI. **(H,J)** Quantification of lipid peroxidation index (BD-11Ox/BD-11Non-Ox). Each dot represents one field of view. n= ∼100 cells per condition. **(K)** Representative images showing internalization of fluorescently labeled Aβ (green) by WT and *Cd44*-KO primary astrocytes (GFAP, red) with or without OPN treatment. Internalized Aβ appears in white. **(L)** Quantification of internalized Aβ calculated as % of Aβ signal overlapping with GFAP area. Each dot represents one image n=∼60 cells per condition. Data is presented as mean ± SEM. Statistical analyses were performed using One-way ANOVA with Tukey’s multiple comparison test (B, D, and L) or 2-sided Student’s *t* test (H and J). \*\**p* < 0.01, \*\*\**p* < 0.001, \*\*\*\**p* < 0.0001 Scale bars: 20 μm.

### OPN-CD44 signaling modulates lipid peroxidation and phagocytosis in astrocytes

To gain molecular insight into OPN-CD44 signaling, we performed quantitative proteomic profiling of WT and *Cd44*-KO primary astrocytes with or without OPN treatment for 24 hours (**Table S2**). Consistent with reported CD44 functions^10,25,42^, GO term analysis revealed downregulation of stress response, hyaluronan metabolism, and cell migration pathway proteins in *Cd44*-KO astrocytes compared to WT controls (**Figure S8**). In OPN-treated WT astrocytes, we observed upregulation of pathways related to mitochondrial activity and general metabolic processes, accompanied by downregulation of phagocytosis-associated proteins. (**Figure 4E**). To directly assess the OPN-CD44 signaling, we compared OPN-treated WT versus OPN-treated *Cd44*-KO astrocytes, which revealed enrichment of reactive oxygen species (ROS) signaling, lipid metabolism, and inflammatory response pathways in WT astrocytes (**Figure 4F**), suggesting that CD44 serves as a key modulator of these OPN-driven metabolic and inflammatory programs.

Previous studies have shown that OPN promotes ROS generation in cancer^43^ and stroke models^44^. Consistent with this, our proteomic analysis revealed enriched oxidative signatures in OPN-treated astrocytes. Because elevated ROS can drive lipid peroxidation, we next asked whether the lipid accumulation observed in OPN-treated astrocytes included peroxidized species. Using BODIPY C11 (581/591), a fluorescent lipid peroxidation sensor that shifts emission from red to green upon oxidation, we found that OPN-treated astrocytes exhibited a pronounced red-to-green shift, indicative of increased lipid peroxidation (**Figures 4G and 4H**). Similar increases were observed in human iPSC-derived astrocytes (**Figures 4I and 4J**), underscoring the conserved nature of the OPN-CD44 response across species. This effect was comparable to that induced by Rotenone/Antimycin A (Rot/AA) (**Figure S9**), a combination of inhibitors that triggers mitochondrial dysfunction and ROS production^45^.

Given the transcriptional suppression in phagocytosis-related pathways in response to OPN, we next tested whether OPN directly impairs Aβ uptake through CD44 signaling. Under baseline conditions, both WT and *Cd44*-KO astrocytes effectively internalized Aβ. However, OPN treatment significantly reduced Aβ uptake in WT astrocytes, while this effect was abolished in *Cd44*-KO cells (**Figures 4K and 4L**). These findings suggest that CD44 is essential for mediating OPN’s suppression effect on astrocyte Aβ phagocytosis.

### Astrocyte-specific *Cd44* deletion reduces microglia reactivity and OPN levels

To investigate the functional role of astrocytic CD44 in AD pathogenesis, we generated a *Cd44* floxed (*Cd44*^fl/fl^) mouse line by inserting two LoxP sequences across exon 3 of the *Cd44* locus using CRISPR/Cas9 (**Figure S10**) and produced astrocyte-specific *Cd44* knockout mice by crossing this allele with two glial Cre drivers: the inducible Aldh1l1^CreER^ (cKO) and the constitutive GFAP^Cre^ (GcKO) **(Figure 5A)**, which were further expressed on the 5xFAD background. For inducible deletion, we administered tamoxifen (75 mg/kg body weight) intraperitonially once daily for 5 days at 5–7 weeks of age^46^. Immunocytochemistry analysis in 6-month-old mice confirmed efficient CD44 depletion in astrocytes in both models (**Figures 5B and 5D**).

**Figure 5.**
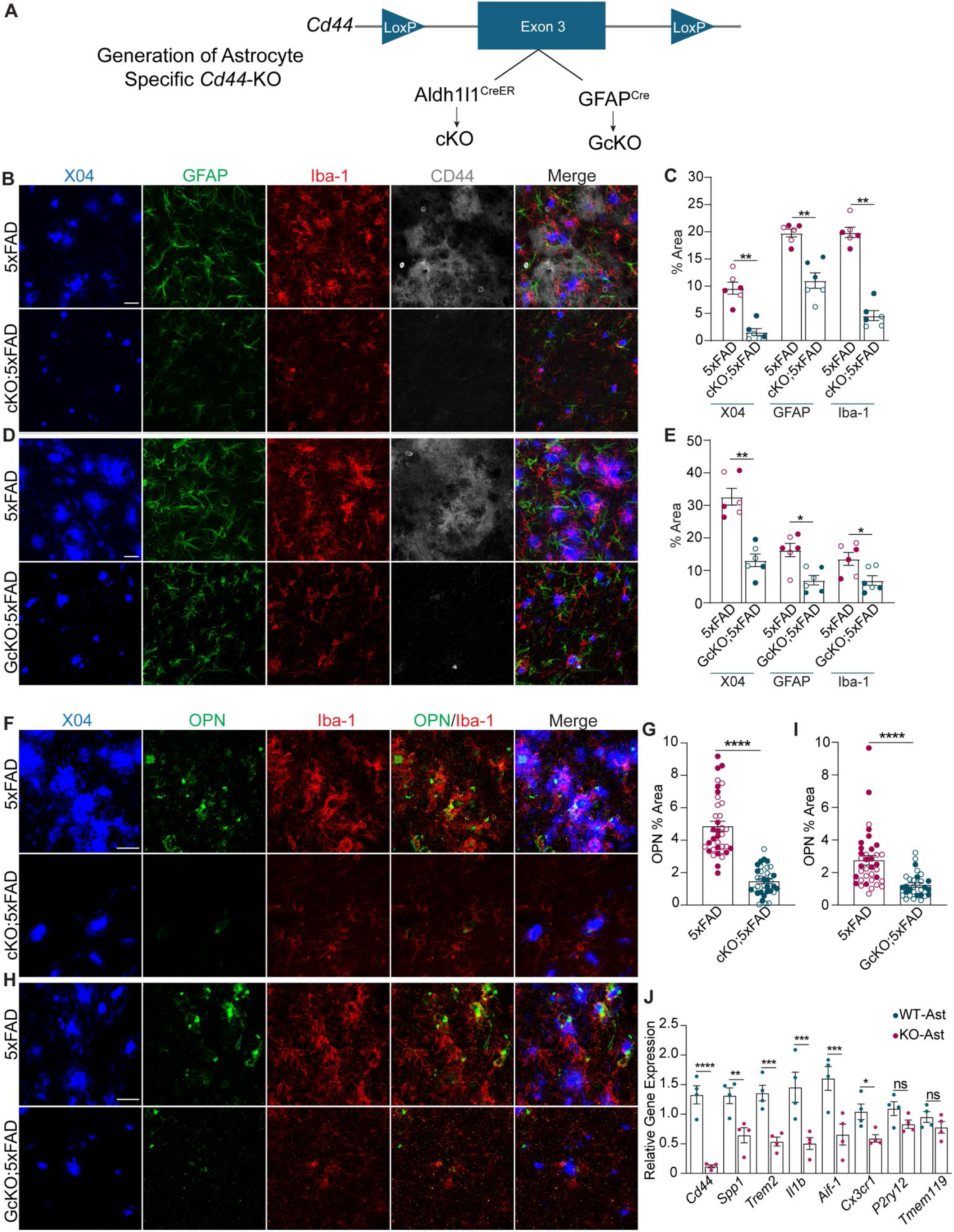
Astrocyte-specific deletion of *Cd44* attenuates microglial reactivity and Aβ pathology in 5xFAD mice. **(A)** Schematic diagram of the Cd44 floxed allele and generation of cKO and GcKO. **(B and D)** Representative images of methoxy-X04 (X04), GFAP, Iba-1, and CD44 in the subiculum of 6-month-old 5xFAD and cKO;5xFAD **(B)** or 5xFAD andGcKO;5xFAD mice **(D)**. **(C and E)** Quantification of percent area for X04, GFAP, and Iba-1 corresponding to the images in (A) and (C) respectively. Each dot represents an average of six images collected from two separate brain sections per mouse. n=6 mice per group. **(F and H)** Representative images of X04, OPN and Iba-1 in the subiculum of 6-month-old 5xFAD and cKO;5xFAD **(F)** or 5xFAD and GcKO;5xFAD mice **(H)**. **(G and I)** Quantification of OPN positive percentage area corresponding to (E) and (G) respectively. n=6 mice per group. For each mouse, six images were analyzed, derived from two brain sections (three images per section). Each dot represents one image. **(J)** qPCR analysis of co-culture of WT microglia with either WT astrocytes (WT-Ast) or *Cd44-*KO astrocytes (KO-Ast) for 48 hours. Data shown is normalized to housekeeping gene GAPDH. Graphs display mean ± SEM. Filled circles denote male, open circles female. Pairwise comparison by two-sided Student’s *t* test. \**p* < 0.05, \*\**p* < 0.01; \*\*\**p* < 0.001; \*\*\*\**p* < 0.0001; ns, not significant. Scale bars: 20 µm.

Quantification of methoxy-X04 labeled Aβ plaques revealed significantly reduced plaque burden in both cKO;5xFAD and GcKO;5xFAD mice compared to 5xFAD controls (**Figures 5B-5E**). This reduction was accompanied by attenuated astrocyte and microglia reactivity, as assessed by GFAP and Iba-1 fluorescence intensity (**Figures 5B-5E**). Immunostaining with an anti-OPN antibody further showed a marked reduction of OPN in microglia surrounding Aβ plaques in both cKO and GcKO animals (**Figures 5H and 5I**). These findings demonstrate a potent effect of astrocytic CD44 in regulating plaque pathology and microglial responses, potentially through the OPN/*Spp1* axis.

To directly test whether astrocytic CD44 modulates microglial phenotype, we performed astrocyte-microglia co-culture experiments using WT microglia paired with either WT or *Cd44*-KO astrocytes. qPCR analysis after 48 hours revealed significantly decreased expression of *Spp1* and key microglial activation markers (*Trem2*, *Il1b,* and *Aif-1*) in *Cd44* deficient astrocytes compared to WT controls. Among homeostatic genes, *Cx3cr1* was modestly but significantly reduced, while other homeostatic markers *(P2ry12*, and *Tmem119)* were not significantly changed (**Figure 5J**).

Together, these findings establish astrocytic CD44 as a key modulator of OPN/*Spp1* expression and microglia activity during Aβ pathology.

### Astrocytic *Cd44* deletion enhances plaque compaction and rescues synaptic and cognitive deficits

To further characterize the Aβ and microglial phenotypes, we performed 3D reconstruction of methoxy-X04^+^ plaques and associated Iba-1^+^ microglia in 6-months-old 5xFAD and cKO;5xFAD mice. Although the total plaque number was unchanged, plaque volume was significantly reduced in cKO;5xFAD animals (**Figures 6A-6C**). This reduction was associated with decreased Iba-1^+^ microglial volume (**Figure 6D**), and reduced dystrophic neurites marked by LAMP1 immunoreactivity (**Figures 6E and 6F**). Similar results were also observed in GcKO;5xFAD mice (**Figure S11**).

**Figure 6.**
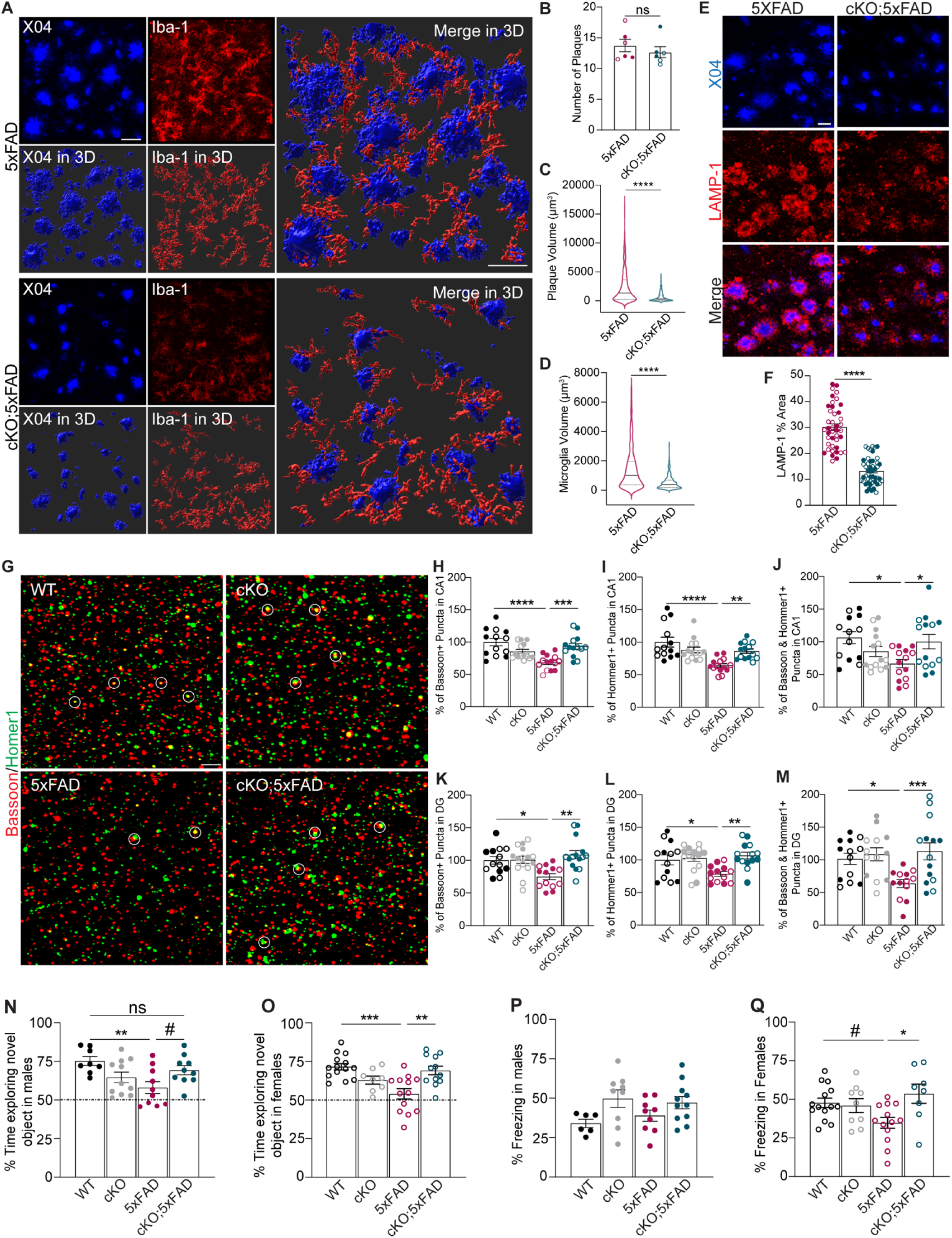
Astrocytic *Cd44* deletion ameliorates amyloid burden, rescues synaptic integrity, and cognitive impairments. **(A)** Representative X04 and Iba-1 images with 3D reconstruction from the subiculum of 6-month-old 5xFAD and cKO;5xFAD mice. **(B)** Quantification of plaque numbers in 5xFAD and cKO;5xFAD animals. Each dot represents the average plaque count from six images collected from two independent brain sections per mouse. n=6 mice per group. **(C, D)** Violin plots showing the volume of X04^+^ plaques **(C)** and microglial volume per plaque **(D)** in 5xFAD and cKO;5xFAD. **(E)** Representative images of X04 and LAMP-1 in 6-month-old 5xFAD and cKO;5xFAD animals. **(F)** Quantification of LAMP-1 percent area. n=6 mice per group. For each mouse, six images were analyzed, derived from two intendent brain sections (three images per section). Each dot represents one image. **(G)** Representative images of hippocampal CA1 region stained for presynaptic (Bassoon) and postsynaptic (Homer1) markers in WT, cKO, 5xFAD and cKO;5xFAD mice. The circles mark selected co-localized puncta. **(H–J)** Quantification of percentage of Bassoon^+^ (H), Homer1^+^ (I), and Bassoon and Homer1colocalized synaptic puncta (J). **(K–M)** Same quantification in DG area. **(N-Q)** Behavioral analysis of WT, cKO, 5xFAD and cKO;5xFAD mice at 6-8 months of age. **(N, O)** Quantification of percentage of time exploring the novel object in male **(N)** and female **(O)** mice. Dashed line represents a 50% change of random object exploration. WT: n= 8♂and 14♀; cKO: n=11♂and 8♀; 5xFAD: n=10♂ and 13♀; and cKO;5xFAD: n=10 ♂ and 12♀. **(P, Q)** Quantification of percentage of freezing time in contextual fear conditioning in males **(P)** and female **(Q)**. WT: n=6♂ and 14♀; cKO: n=9♂ and 9♀; 5xFAD: n=11♂ and 8♀; and cKO;5xFAD: n=10 ♂and 11♀. Graphs are displayed as mean ± SEM. Filled circles denote male, open circles female. B-D and F were performed by pairwise comparison by 2-sided Student’s *t* test; I-L were performed by One-way ANOVA with Tukey’s multiple comparison test. \**p* < 0.05; \*\**p* < 0.01, \*\*\**p* < 0.001, \*\*\*\**p* < 0.0001; ^#^*p* = 0.075; ns, not significant. Scale bars in A and E, 20 µm; in G: 3 µm.

We next assessed synaptic integrity by staining for the presynaptic marker Bassoon and the postsynaptic marker Homer1 in the hippocampal CA1 and dentate gyrus (DG) of 6-month-old WT, cKO, 5xFAD and cKO;5xFAD mice. As expected, 5xFAD mice showed marked reductions of Bassoon^+^, Homer1^+^, and co-localized synaptic puncta compared to WT and cKO mice (**Figures 6G-6M**). Notably, cKO;5xFAD mice displayed a robust restoration of synaptic puncta density and colocalization, reaching levels comparable to WT and cKO controls (**Figures 6G-6M**).

To determine whether the synaptic marker improvements translated into better cognitive performance, we conducted behavioral analyses in 6-8-month-old WT, cKO, 5xFAD and cKO;5xFAD mice using open field, novel object recognition (NOR) and fear conditioning tests. The open field test revealed no group differences in locomotor activities or movement behavior (**Figures S12A-S12F**). During the NOR training phase, all groups spent similar time exploring the identical objects (**Figure S12G**), and freezing behavior during the conditioning phase of the fear conditioning test was also comparable across groups (**Figures S12H-S12J**), indicating intact locomotion and baseline neurological function among all groups.

In the NOR test, 5xFAD mice displayed reduced novel object exploration, whereas cKO;5xFAD mice showed improved performance, with trends toward significance in males (**Figure 6N**) and statistically significant improvement in females (**Figure 6O**). In the contextual fear conditioning test, male 5xFAD mice did not exhibit clear deficits (**Figure 6P**), limiting the ability to assess rescue; however, female cKO;5xFAD mice performed significantly better than 5xFAD females (**Figure 6Q**).

Overall, these findings demonstrate that astrocyte-specific deletion of *Cd44* not only attenuates plaque-associated neuritic and synaptic damage but also improved cognitive function, particularly in female 5xFAD mice.

## Discussion

CSPs are critical mediators of cell–cell communication and receptor–ligand signaling in the brain, and their accessibility on the cell surface makes them particularly tractable therapeutic targets. While the role of microglial CSPs in AD is well established^47^, the contribution of astrocytic CSPs remains less understood. In this study, we applied iPEEL to profile astroglial CSP changes in response to Aβ pathology in 5xFAD mice. This genetic system enables consistent and robust cell-surface labeling, avoiding the mouse-to-mouse variability inherent to AAV-mediated delivery previously used to identify astrocytic and neuronal CSPs^48^ and allowed us to generate the first systematic profile of astroglial CSP remodeling in response to Aβ pathology in 5xFAD mice. Our findings establish a framework for expanding iPEEL to additional cell types, brain regions, and disease models for CSP discovery and mechanistic and therapeutic exploration.

Overwhelming evidence implicates dysregulated immune and lipid metabolic pathways in late-onset AD^47,49–53^. Here, we identify astroglial CD44, discovered through iPEEL profiling, as a critical mediator of both processes. We uncover an OPN-CD44 axis that modulates the microglia-astrocyte crosstalk: while OPN binds astrocytic CD44 to induce γ-secretase dependent lipid accumulation, astrocytic CD44 promotes *Spp1*/OPN expression and microglial reactivity. This axis is upregulated in AD mouse models and human brains, particularly in peri-plaque regions, and is reversed by astrocytic *Cd44* ablation.

CD44 has been linked to lipid metabolism in peripheral tissues, where its expression is elevated in white adipose tissue under obesity conditions, and its inactivation confers resistance to high-fat diet–induced lipid dysregulation and metabolic impairment^54–56^. Here we observed increased BODIPY-positive lipid accumulation in CD44^+^ astrocytes, levels of which were further elevated in 5xFAD mice. Using primary mouse astrocyte cultures and human iPSC-derived astrocytes, we demonstrate a causal role of CD44 in lipid induction. This process requires OPN and γ-secretase but is independent of CD44’s canonical ligand HA or factors that induce astrocyte reactivity, particularly TIC or IL6. Of interest, OPN-CD44 signaling promotes glioma growth via γ-secretase cleavage of CD44 and subsequent binding of the CD44 intracellular domain (CD44ICD) to CBP/p300, which drives transcriptional activation of hypoxia-inducible factor 2α (HIF-2α)^24^. It is therefore plausible that a similar mechanism contributes to the lipid induction observed here. However, given the broad effects of γ-secretase inhibition, the specific target mediating the lipid phenotype downstream of OPN–CD44–γ-secretase signaling and whether it requires CD44ICD-dependent transcriptional regulation remains to be determined.

The accumulated lipids are enriched in peroxidized species linked to pro-inflammatory response, metabolic reprogramming and astrocyte dysfunction including impaired phagocytosis^57–59^, and is consistent with our proteomic data, which revealed dysregulated immune, metabolic, and phagocytic pathways and significant upregulation of lipid and reactive oxygen species pathways in OPN-treated wild-type cultures compared to vehicle-treated wild-type or OPN-treated *Cd44* KO cultures, respectively. Functionally, OPN treated wild-type astrocytes exhibited reduced Aβ phagocytosis, and these phenotypes were rescued by *Cd44* loss. An intimate relationship between lipid, immune, and phagocytic pathways is also documented in microglia in which lipid-accumulating microglia are proinflammatory and phagocytosis defective^53,60^.

We show that CD44 was elevated in a subset of astrocytes in close proximity to Aβ plaques, and that astrocytic deletion of *Cd44* in 5xFAD mice reduces GFAP and Iba1 immunoreactivities and results in more compact methoxy-X04 positive plaques. These changes were accompanied by decreased dystrophic neurites, restored synaptic integrity, and improved cognitive performance, particularly in female 5xFAD mice. The sex difference is likely due to the more pronounced Aβ pathology in female 5xFAD mice^61,62^, providing a more sensitive baseline for detecting behavioral rescue. Indeed, we were unable to detect impaired contextual fear conditioning in male 5xFAD mice at 6-7 months, making it difficult to assess the impact of CD44 loss. Nevertheless, the data combined clearly demonstrate a potent role for astrocytic CD44 in regulating microglia and Aβ dynamics in both male and female 5xFAD mice.

The observed reduction in gliosis and Aβ pathology by *Cd44* deficiency could reflect a primary astrocytic CD44-Aβ interaction, with reduced microglia reactivity as a secondary consequence of diminished Aβ burden. Alternatively, CD44 could engage a direct astrocyte-microglia interaction that jointly influence Aβ pathology. Our co-culture data showing reduced expression of *Spp1* and other microglial genes when co-cultured with *Cd44* deficient astrocytes supports a direct astrocyte-microglia interaction. A reciprocal regulation of microglial Spp1/OPN by astrocytic CD44 is further supported by a recent iPSC-derived neuron-astrocyte-microglia co-culture study showing *SPP1* induction in microglia co-cultured with CD44-positive astrocytes independent of neurons^63^

Interestingly, the CD44 mediated astrocyte-microglia-Aβ dynamics we observed parallel those of Plexin-B1, a guidance receptor and astrocyte hub gene enriched in the AD gene network^64^. Genetic deletion of *Plexin-B1* in APP/PS1 mice similarly reduced astroglial and microglial reactivities, plaque size and dystrophic neurites^64^. These effects were attributed to Plexin-B1’s role in shaping the peri-plaque glial net. CD44 is known to interact with extracellular matrix proteins, including OPN, to regulate cell adhesion and motility^11^. Its intracellular domain binds actin cytoskeleton via Ezrin/Radixin/Moesin (ERM) proteins, enabling cytoskeletal remodeling and cell migration^65^. Notably, both CD44 and Moesin (MSN) have been identified as top astrocytic hub proteins enriched in AD^30,66,67^. These observations raise the possibility that microglial OPN acts as a chemoattractant to recruit CD44^+^ astrocytes to the peri-plaque region via an MSN-dependent mechanism, where ensuing astrocyte–microglia interactions jointly modulate the peri-plaque glial net and Aβ pathology.

*Spp1* is a well-established DAM gene strongly upregulated in brains of AD mouse models and human patients^14,68,69^. Qiu et al. showed that OPN is selectively produced in CD11c^+^ microglia, and its expression divides DAM into two functionally distinct subsets with OPN^-^ microglia being protective whereas OPN^+^ microglia being pathogenic^68^. Thus, whether or not astrocytic CD44 directly regulates *Spp1*/OPN expression, the drastic reduction of OPN in microglia of 5xFAD mice lacking astrocytic *Cd44* is expected to reduce pathogenic microglia and improve its function. Supporting the functional relevance of the CD44-OPN crosstalk, both the astrocytic *Cd44* knockout and *Spp1* deletion in 5xFAD mice show dampened microglia and immune activity, reduced Aβ pathology, and improved cognitive function^68^. However, because *Spp1* is also expressed in other immune cells in the brain^70,71^, defining the specific role of microglial OPN-astrocytic CD44 interactions in vivo requires further investigation.

Overall, our findings support a bidirectional OPN-CD44 mediated astrocyte-microglia crosstalk in AD, in which OPN engages astrocytic CD44 to drive lipid accumulation, while astrocytic CD44 promotes microglial *Spp1*/OPN expression and associated immune dysfunction, forming a vicious cycle. Combined with the fact that both OPN and CD44 are elevated early in AD and correlate with the neuropathology severity and cognitive decline^30,68^, these findings provide a compelling rationale for targeting astrocytic CD44 to disrupt this cycle and restore immune and lipid homeostasis.

Building on this rationale, CD44 has emerged as a promising therapeutic candidate in AD. Its detectability in CSF during the asymptomatic phase^30,31^ and its functional contribution to the disease pathology, as demonstrated in our study, underscores its clinical relevance. Although no CD44-directed therapies have been approved for CNS indications, several candidates have been advanced for oncology indications, including the humanized monoclonal antibody RG7356^72,73^ and a small molecule inhibitor verbascoside^74^, both of which suppress tumor growth. These existing therapeutic agents provide a translational entry point for repurposing CD44-targeted strategies in AD and related neurodegenerative disease. By defining a mechanistic link between CD44 and glial dysfunction, our study lays the foundation for therapeutic interventions that could ultimately reshape the care for Alzheimer’s disease.

## Limitations of the study

To ensure robust HRP expression in astrocytes, we used the GFAP-Cre line for CSP profiling. Because this line exhibits known leaky expression in neurons, some differentially expressed proteins may originate from neurons rather than astrocytes. Astrocyte-specific Cre drivers such as Aldh1l1-CreER may represent better alternatives; however, given the intimate association between astrocytes and neurons, neuronal proteins are likely to be captured even with an astrocyte-restricted driver. This was demonstrated in a recent CSP profiling study using AAV-mediated astrocytic or neuronal HRP expression, which showed extensive overlap in the striatal cell-surface proteome, reflecting cross-labeling at the astrocyte-neuron interfaces^48^. Thus, detection of CSPs from adjacent cell types may be inherent to this proximity-labeling approach. Additional stratification using single-cell transcriptomic datasets will help resolve cell-type-specific contributions.

For CD44, we identified a potent and specific role for the OPN–CD44 interaction in driving lipid accumulation in astrocyte cultures. However, the functional consequences of the lipid phenotype in vivo remain to be defined. Given the presence of multiple OPN receptors and CD44 ligands^11^, the contribution of the specific OPN–CD44 axis in vivo and in AD requires further clarification. Moreover, the mechanisms by which astrocytic CD44 regulate *Spp1* expression and microglial reactivity remain unresolved and warrant deeper investigation.

## ACKNOWLEDGEMENTS

We thank Brittany Reeves and Haiying Liu for expert technical assistance and members of the Zheng lab for valuable comments and suggestions. We acknowledge the support from the Cytometry and Cell Sorting Core at Baylor College of Medicine, with funding from the NIH (P30 AI036211, P30 CA125123, and S10 RR024574). This work was supported by grants from the NIH (R01 NS093652, R01 AG020670, P01 AG066606, and RF1 AG088197 to HZ), Cure Alzheimer Fund (to HZ and JP) and BrightFocus Foundation (to OPR).

## AUTHOR CONTRIBUTIONS

OPR, MG and HZ designed the overall study; OPR and MG performed major biological experiments with support from CSR on iPSC cultures and flow cytometry and SW on behavioral analysis and synaptic marker staining; XZ and JP performed TMT-MS analysis; CC created *Cd44* floxed mice; CQ assisted with the re-analysis of human snRNA-seq data and SYJ carried out proteomic experiments; LL provided the iPEEL mice and HL, JL and TH offered technical support. OPR, MG and HZ wrote the paper, with contributions from XZ and JP on TMT-MS and CC on *Cd44* floxed mice. All authors provided input, read, and approved the manuscript.

## DECLARATION OF INTERESTS

The authors declare no competing financial interests.

## KEY RESOURCE TABLE

| REAGENT or RESOURCE | SOURCE | IDENTIFIER |
| --- | --- | --- |
| **Antibodies** |  |  |
| Rat Anti-CD44 | Invitrogen | Cat# 14044185 |
| Rabbit Anti-Iba1 | Wako | Cat# 01919741 |
| Rabbit Anti-S100β | Sigma-Aldrich | Cat# S2644 |
| Goat Anti-GFAP | Sigma-Aldrich | Cat# SAB2500462 |
| Rat Anti-LAMP1 | BD Pharmingen | Cat# 553792 |
| Goat Anti-SPP1 | R&D Systems | Cat# AF808 |
| Rabbit Anti-HA | Cell Signaling | Cat# 3724 |
| Mouse Anti-HA | Cell Signaling | Cat# 2367 |
| Hyaluronic Acid Binding Protein | Millipore | Cat# 385911 |
| Rabbit Anti-Bassoon | SYSY Antibodies | Cat# 141003 |
| Chicken Anti-Hommer1 | SYSY Antibodies | Cat# 16006 |
| Rabbit Anti-β-Actin | ProteinTech | Cat# 205361AP |
| Rabbit Anti-PSTAT3 | Cell Signaling | Cat# 9145S |
| Rat anti-mouse ACSA-2 Antibody, APC | Miltenyi Biotec | Cat# 1130117535 |
| Rat Anti-Mouse CD16/CD32 (Mouse BD Fc Block™) | BD Biosciences | Cat# 553141 |
| Anti-Amyloid Precursor Protein antibody [Y188] | Abcam | Cat# ab32136 |
| Alexa Fluor 488 Donkey Anti-Rat | Invitrogen | Cat# A21208 |
| Alexa Fluor 488 Donkey Anti-Chicken | Invitrogen | Cat# A78948 |
| Alexa Fluor 488 Donkey Anti-Goat | Invitrogen | Cat# A11055 |
| Alexa Fluor 647 Donkey Anti-Rat | Invitrogen | Cat# A78947 |
| Alexa Fluor 647 Donkey Anti-Goat | Invitrogen | Cat# A21447 |
| Alexa Fluor 555 Donkey Anti-Rabbit | Invitrogen | Cat# A31572 |
| IRDye® 800CW Anti-Rabbit | LI-COR | Cat# 926-32211 |
| IRDye® 680RD Anti-Rat | LI-COR | Cat# 926-68076 |
| HRP-Conjugated Streptavidin | Thermo Fisher | Cat# N100 |

| **Chemicals, peptides, and recombinant proteins** |  |  |
| --- | --- | --- |
| Papain | Worthington Biochemical | Cat # LK003172 |
| DNase | Worthington Biochemical | Cat# LK003178 |
| Percoll Plus | MilliporeSigma | Cat# E0414 |
| Recombinant IL-1α | Cell Signaling | Cat# 62643S |
| Recombinant Tnf-α | Cell Signaling | Cat# 16769S |
| Recombinant C1q | BioSource | Cat# MBS147305 |
| Recombinant Human OPN | R&D | Cat# 1433-OP/CF |
| Recombinant Mouse OPN | R&D | Cat# 441-OP/CF |
| Hyaluronic Acid Powder | Sigma-Aldrich | Cat# H5388 |
| DAPT | Tocris | Cat# 2634 |
| Compound E | Medchem Express | Cat# HY-14176 |
| Beta-Amyloid (1-42), FAM-labeled | Anaspec | Cat# AS23526-01 |
| Methoxy-X04 | Tocris Bioscience | Cat# 863918789 |
| BODIPY 558/568 | Thermo Fisher Scientific | Cat# D3835 |
| BODIPY 581/591 Lipid Peroxidation Sensor | Thermo Fisher Scientific | Cat# D3861 |
| Fluoromount-G | EMS | Cat# 1798425 |
| BSA Protease | Akron | Cat# AK1391100 |
| Recombinant Human OPN | R&D | Cat# 1433-OP/CF |
| Recombinant Mouse OPN | R&D | Cat# 441-OP/CF |
| Trypsin Inhibitor | Sigma-Aldrich | Cat# 93620 |
| Trypsin | Gibco | Cat# 27250018 |
| DNASE1 | Sigma-Aldrich | Cat# SLCR1247 |
| DNQX | Cayman | Cat# 14583 |
| AP5 | Cayman | Cat# 14539 |
| TTX | Hello Bio | Cat# HB1035 |
| BxxP | ApexBio | Cat# A8012 |
| DTT | Sigma-Aldrich | Cat# D063 |
| 4X Laemmli sample buffer | Bio-Rad | Ca# 1610747 |
| Triton X-100 | Sigma-Aldrich | Cat# X100 |
| 1M Tris-HCl, pH 8.0 | Invitrogen | Cat# 15568-025 |
| Protease Inhibitor Cocktail | Sigma-Aldrich | Cat# P884 |
| Phenylmethylsulfonylfluoride, PMSF | Sigma-Aldrich | Cat# P-7626 |
| Streptavidin beads | Thermo Fisher | Cat# 88816 |
| NeutrAvidin Protein, DyLight™ 650 | Invitrogen | Cat# 84607 |
| DAPI | Abcam | Cat# ab228549 |
| Normal Donkey Serum | Jackson Immuno Research Labs | Cat# 017-000-121 |
| Tamoxifen | Sigma-Aldrich | Cat# T5648 |
| Lysyl Endopeptidase | Sigma-Aldrich | Cat# 121-05063 |
| C18 ultra micro spin columns | Harvard Apparatus | Cat# 74-7206 |
| Matrigel | Millipore | Cat# CLS354234 |
| mTeSR Plus medium | Stem Cell Technologies | Cat# 100-0276 |
| Accutase | Stem Cell Technologies | Cat# 07920 |
| ROCK inhibitor | StemMACS | Cat# Y-27632 |
| Dual SMAD inhibitor cocktail | Stem Cell Technologies | Cat# 08581 |
| Neural progenitor medium | Stem Cell Technologies | Cat# 05833 |
| Astrocyte Medium | ScienCell | Cat# 1801 |
| Critical commercial assays |  |  |
| RNAeasy | Qiagen | Cat# 74106 |
| Silver Stain Plus Kit | Bio-Rad | Cat# 161-0449 |
| Pierce BCA protein assay | Pierce | Cat# 23228 |
| 4-20% SDS-PAGE gel | Bio-Rad | Cat# 4561094 |
| Sera-Mag Speedbead carboxylate-modified [E3] magnetic particles | Cytiva | Cat# 65152105050250 |
| Sera-Mag carboxylate-modified [E7] magnetic particles | Cytiva | Cat# 24152105050250 |
| Trypsin/Lys-C | Thermo Fisher | Cat# A40007 |
| PepMap™ 100 C18 column | Thermo Scientific | Cat# 164534 |
| FAM-labeled Aβ | Anaspec | Cat# AS23526-01 |
| iScript Reverse Transcription Supermix reagent | Bio-Rad | Cat# 170-8840 |
| iTaq Universal SYBR Green Supermix | Bio-Rad | Cat# 172-5124 |
| **Deposited data** |  |  |
| MassIVE | https://massive.ucsd.edu | Accession number MSV000099996.<br>username MSV000099996_reviewer<br>Password: BCM_iPEEL_AST_6M |
| **Experimental models: Cell lines** |  |  |
| Human induced pluripotent stem cells | BCM Stem Cell Core | male, GM23815, 196 iPSCs clone-13 |
| **Experimental models: Organisms/strains** |  |  |
| Mouse: 5xFAD | JAX<sup>75</sup> | Strain# 034840 |
| Mouse: GFAP<sup>Cre</sup> | JAX<sup>76</sup> | Strain# 24098 |
| Mouse: Aldh1l1<sup>CreER</sup> | JAX | Strain# 31008 |
| Mouse: Cd44-KO | JAX | Strain# 005085 |
| Mouse: Cd44<sup>fl/+</sup> | Generated in this study |  |
| Mouse: HRP<sup>loxP-STOP-loxP</sup> | JAX<sup>7</sup> | Strain# 037698 |
| **Oligonucleotides** |  |  |
| See Table S3 for a complete list of primer sequences |  |  |
| **Software and algorithms** |  |  |
| Leica confocal suite software | Leica<br>(https://www.leicamicrosystems.com) | RRID:SCR_013673 |
| Imaris (version 9.1) | Bitplane<br>(www.bitplane.com) | RRID:SCR_007370 |
| ImageJ (version 1.47) | NIH<br>(https://imagej.nih.gov/ij/) | RRID:SCR_003070 |
| GraphPad Prism (version 9.1.0) | GraphPad Software<br>(www.graphpad.com) | RRID:SCR_002798 |
| R (version 3.5.1) | R Core Team<br>(https://www.r-project.org/) | RRID:SCR_001905 |

### RESOURCE AVAILABILITY

Further information and requests for sources and reagents should be directed to the lead contact, Hui Zheng

#### Materials and data availability

All unique reagents generated in this study are available from the lead contact. The processed proteomic data is provided in Table S1, and Table S2. All mass spectrometry raw data and associated metadata have been deposited in the MassIVE repository and are publicly accessible. The dataset can be accessed at the MassIVE website (https://massive.ucsd.edu) under the accession number **MSV000099996**. Reviewer access is available using the username **MSV000099996_reviewer** (Password: BCM_iPEEL_AST_6M)

### EXPERIMENTAL MODEL AND SUBJECT DETAILS

#### Mice

All procedures were performed in accordance with NIH guidelines and approved by the Baylor College of Medicine Institution Animal Care and Use Committee (IACUC). The HRP^loxP-STOP-loxP^ mice were crossed with the GFAP^Cre^ and 5xFAD mice to generate littermate of HRP, Non-HRP, 5xFAD, and HRP;5xFAD. Because GFAP^Cre^ exhibits leaky expression in neuronal stem cells^77^ and in male germline, only females Cre mice were used for breeding. Both male and female offspring were included in all proximity labeling, biochemical and proteomic experiments. The *Cd44* floxed mouse line was crossed with Aldh1l1^CreER^ and 5xFAD mice separately to generate *Cd44*^fl/fl^; Aldh1l1^CreER^ and *Cd44*^fl/fl^; Aldh1l1^CreER^; 5xFAD mice. To selectively eliminate *Cd44* in adult astrocytes we administered tamoxifen (75 µg/g body weight) intraperitonially daily for 5 days at 5–7 weeks of age^46^. Heterozygous *Cd44*-KO mice were crossbred to generate *Cd44*-KO homozygous and WT littermates for primary glial cell culture experiments.

All mice were maintained on the C57BL/6 background. Both sexes were used in approximately equal numbers, and sex differences were noted where relevant. Unless otherwise specified, all analyses were conducted in 6-month-old mice.

#### Generation of *Cd44* floxed mice

A conditional *Cd44* allele was generated using CRISPR /Cas9-based genome editing approach. Two sgRNAs (upstream sgDNA: AGCAATAGACATGAACGTCTGGG. Downstream sgDNA: TTGTCAGCCAATGCTCCCACAGG) were designed to target intronic sequences immediately upstream and downstream of *Cd44* exon 3, enabling insertion of loxP sites flanking exon 3. A long-single stranded DNA donor containing the 5’ and 3’ homologues arm, the two loxP sites, and exon 3 was synthesized by in vitro transcription and served as a template. This template, together with the above two sgRNAs and a Cas9 mRNA, was microinjected into C57BL/6 fertilized zygotes. Founder mice were screened by PCR and DNA sequencing across both loxP junctions. The floxed allele was confirmed by breeding. The correctly targeted *Cd44*^fl/+^ line was established through germline transmission. Conditional deletion of exon 3 was validated by genotyping of loxP allele and WB of CD44 when Cre is introduced. Mice were backcrossed for more than five generations onto wild-type C57BL/6 mice to remove potential off-target or background mutations.

#### Human iPSC-derived astrocyte cultures

Early-passage human induced pluripotent steam cells (iPSCs) were obtained from the Baylor College of Medicine Stem Cell Core and maintained in Matrigel-coated dishes in mTeSR Plus medium as previously described^41^. Neuronal induction was perfumed using a dual SMAD inhibition monolayer protocol^78^. At ∼80% confluency, iPSCs were dissociated with Accutase, and seeded at 2 × 10⁶ per well in Matrigel-coated 6-well plates using neuronal induction media supplemented with with 10 µM ROCK inhibitor and dual SMAD inhibitor cocktail. After 24 hours, medium lacking ROCK inhibitor was provided and replaced daily for 6–9 days. On day 9, cells were re-plated (1.5 × 10⁵ cells/cm²) and maintained with daily medium changes. By day 16, neuronal progenitor cells (NPCs) were harvested and cultured in neuronal progenitor medium (NPM). Confluent NPCs (days 20-22) were either frozen or used for astrocyte differentiation. Low passage NPCs enriched for CD271⁻/CD133⁺/CD184⁺ were differentiated into astrocytes^79,80^. Cells were seeded at 1.5 × 10⁴ cells/cm² and switched to a 1:1 mix of NPM and Astrocyte Medium 1 (2 % fetal bovine serum, astrocyte growth supplement, and penicillin/streptomycin). From day 3 cultures were fed with astrocyte medium every other day. By day 30, matured astrocytes exhibit characteristic morphology and expressed express GFAP and S100β.

#### Primary astrocyte cultures

Primary astrocytes were prepared as previously described^81^. Briefly, brains from neonatal pups (P0–P3) were dissected in ice-cold HBSS, and meninges were carefully removed. Tissue was digested with 2.5% trypsin at 37 °C for 15 minutes, followed by inhibition with 1 mg/mL trypsin inhibitor. The suspension was centrifuged at 400 × g for 5 minutes, triturated, and resuspended in DMEM containing 10% FBS. Cells were plated into poly-D-lysine (PDL) coated T-75 flasks to establish mixed glial cultures. Cultures were maintained for 2 weeks with media changes every other day. To enrich astrocytes, flasks were shaken at 250 rpm for 2 hours at 37 °C. After a 24-hour recovery in fresh medium, a second 2-hour shake was performed, followed by firm tapping of the flasks to dislodge non-astrocytic cells. Astrocytes were then harvested using 0.25% trypsin/EDTA and replated onto PDL-coated coverslips for experiments.

#### Human iPSC-derived neuronal cultures

As previously described^41^, cells were maintained in Matrigel-coated 6 cm petri dishes (Millipore Sigma) using mTeSR Plus medium and passaged with Accutase (Stem Cell Technologies). For pre-differentiation, iPSCs were plated at 1.5 × 106 cells per well in Matrigel-coated 6-well plates (Millipore Sigma) and cultured in Knockout DMEM/F-12 medium (ThermoFisher) supplemented with 2 μg/mL doxycycline (Sigma-Aldrich), N2 supplement (ThermoFisher), 1X Non-essential amino acids (ThermoFisher), 10 ng/mL BDNF (PeproTech), 10 ng/mL neurotrophin-3 (PeproTech), 1 μg/mL mouse laminin (Millipore Sigma), and 2 μM ROCK inhibitor (Tocris). The medium was replaced 24 hours later, omitting ROCK inhibitor, and pre-differentiation was maintained for 3 days (day −3 to day 0). On day 0, pre-differentiated precursor cells were dissociated with Accutase and re-plated onto poly-D-lysine (PDL) and laminin-coated coverslips at a density of 5 × 104 cells per coverslip in 24-well plates. Neuronal cultures were established in a 1:1 mixture of Neurobasal-A (ThermoFisher) and DMEM/F-12 media, supplemented with B27 and N2 (ThermoFisher), 0.5X GlutaMAX (ThermoFisher), 1X non-essential amino acids, 10 ng/mL BDNF, 10 ng/mL NT-3, 1 μg/mL mouse laminin, and 2 μg/mL doxycycline. On day 7, doxycycline was removed, and half of the medium was replaced every other day up to a total of 3 weeks (day 21) prior to use for experiments.

### METHODS DETAILS

#### Proximity labeling in acute brain slices

We adapted the procedure as previously described^7^. Six-month-old HRP, Non-HRP, and 5xFAD;HRP mice were anesthetized with ketamine/xylazine and perfused with freshly prepared ice-cold artificial cerebrospinal fluid (ACSF) containing (nM): Choline chloride (110), NaHCO3 (25), NaH2PO4·H2O (.25), KCL (2.5), Myo-inositol (3), MgCl2 (3), CaCl2 (1), DNQX (0.02), AP5 (0.05), and 0.0319mg/mL (TTX), supplemented with 4.5g/L of D (+)-Glucose per litter. The ACSF solution was continuously bubbled with a gas mixture of oxygenated 95% O2 and 5%CO2 for at least 30 minutes to maintain pH (7.4) and osmolarity (∼300-310). Brains were rapidly dissected and transferred to a Leica vibratome, where 200-µm thick coronal sections were cut in ice-cold, originated ACSF. Slices encompassing the hippocampal region were collected and transferred to a 10-cm dish with continuous oxygenated ACSF, then placed in a 34 °C water bath for 30 minutes to recover. Recovered slices were transferred to carbonated ACSF-BxxP (100mM) and incubated for 1 hr at 34°C to allow diffusion of biotin-phenol (BxxP) tissue ensuring the slices don’t overlap. The labeling reaction was initiated by transferring the slices into ACSF-BxxP containing 0.003% H2O2 and incubating for 5 min at room temperature without bubbling; gentle swirling ensured even reagent exposure. The reaction was quenched by washing slices 5 times with ice-cold quenching buffer (10 mM sodium ascorbate, 10 mM sodium azide, and 5 mM Trolox in ACSF) using the same volume as in the labelling step. The first wash lasted 1 min followed by four washes of 5 min each. A subset of slices (1-2 per genotype) was fixed overnight in 4% paraformaldehyde (PFA) in phosphate-buffered saline (PBS) at 4°C for immunostaining, while the remaining tissue was snap-frozen in liquid nitrogen and store at -80°C for biochemical and proteomic analysis. One slice from each sample submitted for TMT-MS/MS-quantitative proteomic analysis.

#### Preparation of RIPA buffers

Three variants of modified radioimmunoprecipitation (RIPA) assay buffer were used for tissue lysis and protein enrichment: Normal RIPA, High-SDS, and SDS-free RIPA. All buffers were prepared in 100 mL of Milli-Q water, filter through a 0.44 µm membrane and store at 4°C until use. Immediately before use, all buffers were supplemented with 1X protease inhibitor cocktail and 1mM PMSF. Normal RIPA buffer contained 0 mM Tris-HCl (pH 8.0), 150 mM NaCl, 0.5% sodium deoxycholate, 1% Triton X-100, and 0.2% SDS. High-SDS RIPA buffer was prepared using the same components but with an increased SDS concentration to 1%; while free SDS-RIPA omitted SDS.

#### Lysis and enrichment of biotinylated proteins

Frozen mouse brain slices were lysed one at a time in 100 µL of high-SDS RIPA buffer supplemented with protease inhibitor cocktail and PMSF. Samples were homogenized on ice using disposable motor-driven pestles for 1 min, followed by a second round of grinding. The homogenate was briefly centrifuge, and the pestle was rinse with an additional 200 µL of high-SDS RIPA buffer. Lysates were then spun down and kept on ice before processing. Samples were sonicated twice for 10s each, with a minute of cooling interval on ice in between sonication, then heated to 95°C for 5 min and cooled on ice for 1 min. Subsequently, 1.2 mL of SDS-free RIPA buffer containing protease inhibitor cocktail and PMSF was added, samples were sealed and incubated for 2 hrs. at 4°C with gentle rocking. When enrichment was not performed immediately, samples were flash-frozen in liquid nitrogen and stored at -80°C.

For streptavidin-based enrichment of biotinylated proteins, lysates were centrifuged at 100,000 X g for 30 min at 4°C and the supernatant was collected for binding. Streptavidin magnetic beads (30 mL per sample were prewashed twice with 1 mL of normal RIPA buffer without inhibitors and gently resuspended between washes. Cleared lysates were added to the pre-washed beads and incubated for 1 hr. at 4°C with rotation. Beds were subsequently washed using 1 mL of each of the following buffers: twice with normal RIPA buffer (with inhibitors and PMSF), once with 1 M KCL, once with 0.1 M Na₂CO₃, once with 2 M urea in 10 mM Tris HCL (pH 8.0), and twice with PBS.

An aliquot (100 µL) of the streptavidin bead-enriched lysate was reserved for quality control (QC) by silver staining and streptavidin immunoblotting, while the remaining bead-bound proteins were store at -20°C for subsequent TMT-MS/MS analysis. Beads were mixed with 200 µL of elution buffer containing 4 µL of 1 M DTT (final 20 mM), 4 µL of 100 mM biotin in DMSO (final 2 mM), and 92 µL of distilled water. Samples were boiled at 95°C for 10 min, briefly vortexed and centrifuged, and the eluates were either stored at −20°C or immediately analyzed by SDS–PAGE followed by streptavidin blotting to verify biotinylating efficiency and enrichment consistency across samples.

#### QC validation

Protein electrophoresis was performed in 4-20% SDS-PAGE gel (Bio-Rad) according to the manufacturer’s instructions, followed by silver staining (Pierce Silver Stain Kit) and streptavidin blotting as previously described^7^. For chemiluminescence detection, membranes were incubated with HRP-Conjugated Streptavidin (2.4 µl in 10 ml blocking buffer) and visualized using Clarity Western ECL blotting substrate (Bio-Rad) on a ChemiDoc imaging system (Bio-Rad).

#### Immunoblotting

Total protein from brain tissue samples was extracted using RIPA buffer as described previously^82^. Protein concentrations were determined using a BCA kit (Thermo Fisher). Equal amounts of lysate were speared in a 4-20% SDS-PAGE gel (Bio-Rad) and transferred onto a nitrocellulose membrane. Membrane was blocked in TBS buffer containing 5% bovine serum albumin (BSA) and incubated overnight at 4 °C with appropriate primary antibodies: Anti-HA (1:1000), Anti-b-Actin (1: 10000), Anti-APP (1:1000). After washing, membranes were incubated with IRDye® 800CW or 680RD secondary antibodies (1:10000) for 1 hr. at room temperature. Blots were imaged on ChemiDoc Imaging Systems and intensities were quantified using ImageJ software, normalizing to β-Actin loading control.

#### On-bead digestion

The biotinylated samples were washed three times after enrichment using buffer (50 mM HEPES, 6 M urea, 0.5% NaDoc, pH 8.5)^1^. Beads were resuspended in the same volume of washing buffer, and 10% of the sample was reserved for protein quantification and enrichment quality evaluation by silver staining. The remaining samples were digested sequentially with LysC (protein:enzyme = 100:1; Lysyl Endopeptidase®,#121-05063) for 3 hours and Trypsin (protein:enzyme = 50:1; # V511C) overnight. Reduction and alkylation of the digested peptides were carried out with DTT and IAA, respectively. The samples were then acidified to ∼pH 3 by adding 5% formic acid prior to desalting. Desalting was performed using ultra micro spin columns packed with C18 material (#74-7206, Harvard Apparatus). Finally, all samples were dried in a SpeedVac before TMT reagent labeling.

#### TMTpro 18-plex labeling and LC/LC-MS/MS measurements

Peptides were resuspended in 50 mM HEPES (pH ∼8.5) at a protein concentration of ∼1 µg/µL and fully labeled with TMTpro 18-plex reagent (TMT:protein = 1.5:1). Labeled samples were equally pooled, quenched, and desalted using a 100 mg Sep-Pak C18 cartridge prior to basic offline fractionation. Offline basic reverse-phase (RP) LC was performed with an ACQUITY UPLC BEH C18 column (1.7 *μ*m particle size, 2.1 × 150 mm, Waters) using buffer A (10 mM ammonium formate, pH 8.0) and buffer B (10 mM ammonium formate in 90% acetonitrile, pH 8.0). Pooled samples were fractionated with a gradient of 15–50% buffer B, and the resulting fractions were concatenated into 24 fractions.

The concentrated samples were dried by SpeedVac, resuspended in 5% formic acid, and analyzed on an Orbitrap Exploris 480 MS (Thermo Fisher Scientific) using a 95 min nano-LC gradient of 15–55% buffer B at 250 nL/min (buffer A: 0.2% FA, 5% DMSO; buffer B: buffer A with 65% acetonitrile). Mass spectrometry was performed in positive mode. MS1 scans were acquired at 60,000 resolutions with a 460–1600 m/z range and a maximum ion injection time of 50 ms. MS2 scans were acquired in data-dependent mode with up to 20 scans per cycle, at 60,000 resolution, starting from 120 m/z, with a maximum ion injection time of 110 ms, a 1.0 m/z isolation window with a 0.2 m/z offset, HCD fragmentation at 34% normalized collision energy, and a 20s dynamic exclusion.

#### Proteomics data analysis for in vivo samples

Protein identification, filtering and quantification were analyzed using the JUMP software suite^5^. Search parameters included precursor and product ion mass tolerance of 15 ppm, a maximum of three variable modifications, full tryptic specificity, and up to two missed cleavages. Static modifications were set as TMT tag (+304.20715) and cysteine carbamidomethylating (+57.02146), and methionine oxidation (+15.99492) was specified as a dynamic modification. Peptides with a minimum length of seven amino acids were considered, and multistep FDR filtering was applied prior to quantification.

Quantitative data were log2-transformed and median-normalized. Differential expression analysis was performed in R using the LIMMA package (v3.62.2). Enriched biotinylated proteins were first defined by the cutoffs p < 0.05 (Cre vs. non-Cre) and log2 fold change (Cre/non-Cre) > 0. From this enriched protein set, differentially expressed (DE) proteins were identified as those additionally meeting the criteria *p* < 0.05 and |log2 fold change (Cre-FAD vs. Cre-WT) > 0.16. Volcano plots were generated using the EnhancedVolcano package (v1.24.0).

#### Mouse tissue immunofluorescence staining

Mice were anesthetized with ketamine/xylazine and perfused with ice-cold saline (0.9% NaCl). Brains were extracted and fixed overnight in 4% PFA at 4 °C and dehydrated in 30% sucrose until sectioning. Sagittal brain slices of 30 μm thickness were cut using Leica freezing microtome, and stored in cryoprotectant at -20 °C. For staining, free floating sections were washed in PBS and blocked for 1 hour at room temperature in 4% normal donkey serum (NDS) with 0.4%Triton-X100 in PBS, with gentle rocking. Sections were incubated with appropriate primary antibodies diluted in blocking buffer overnight at 4 °C, such as anti-CD44, anti-Iba-1, anti- S100b, anti-GFAP, anti-HA, anti-Hommer1 (1:500 each); anti-SPP1 (1:100) and anti-Bassoon (1:400), and anti-LAMP1 (1:400). The following day, sections were washed 3 x 5 min with PBS, then incubated for 1 hr. at room temperature with appropriate fluorescent secondary antibodies (1:1000) and DAPI (1:1000), both diluted in blocking buffer. After 3 washes with PBS, sections were mounted on glass slides, air-dried in the dark, and cover slipped with Fluoromount-G.

For methoxy-X04 staining, antibody-stained sections were dried on slides overnight, protected from light. The following day, slides were briefly rehydrated on PBS (30 s), then subsequently incubated in 40% ethanol/PBS (30 s). Sections were stained with 1 μM methoxy-X04 in 40% ethanol/PBS for 30 s, followed by graded rinses in 40%, 70%, and 90% ethanol/PBS (30 s each). Slides were allowed to dry prior to mounting with fluoromount G.

#### In vitro Immunofluorescence staining

iPSC-derived astrocytes on Matrigel-coated coverslips and primary astrocytes on poly-D-lysine (PDL)-coated coverslips were fixed post-treatment with 4% PFA for 20 min at room temperature followed by three washes in PBS. Coverslips were then incubated in blocking buffer (4% NDS with 0.4% Triton-X100 in PBS) for 1 hr at room temperature. After blocking, cells were incubated overnight at 4 °C with appropriate primary antibodies diluted in blocking buffer. The following day, the coverslips were washed with PBS, followed by incubation in fluorescent secondary antibody diluted in blocking buffer for 1-2 hr at room temperature. The coverslips were washed with PBS three times prior to mounting on glass slides with fluoromount G.

#### In vivo lipid accumulation assessment

A FACS-based concurrent brain cell type acquisition (CoBrA) method we developed was used for astrocyte isolation and lipid assessment^83^. Briefly, brains were perfused with PBS. Following removal of olfactory bulb and cerebellum, the tissue was minced and digested with papain and DNAse. After incubation, papain digestion was neutralized by adding ice-cold HBSS+ (HBSS with 2 mM EDTA and 0.5% BSA) after which cells were pelleted and subjected to 5-6 rounds of trituration for mechanical dissociation. The suspension was filtered, and the pellet was resuspended in 20% Percoll PLUS in 1×PBS and centrifuged with low brake to remove myelin debris. Dissociated single cells were incubated in HBSS+ containing Mouse BD Fc-Block (1:100) and labeled with anti-ACSA2-APC (1:100), anti-CD44 (1:50) and BODIPY (1:1000). Astrocytes were identified as ACSA-2+ cells, this astrocyte population was further assessed for CD44 and BODIPY expression. The cells were analyzed and sorted using a BD FACS Aria II cell sorter.

#### In vitro lipid accumulation treatment and staining

Primary mouse astrocytes were seeded onto PDL coated coverslips at a density of 100,000 cells per well, and human iPSC-derived astrocytes were seeded onto Matrigel-coated coverslips at 50,000 cells per well. Cells were allowed to adhere and recover for 24 hours before treatment. Depending on the experimental design, cultures were then treated for 24 hours with recombinant mouse or human OPN (1500 ng/mL), 1.6-2MDa hyaluronic acid (HA; 450 µg/mL), the TIC cytokine cocktail (IL-1α, 30 ng/mL; TNFα, 5 ng/mL; C1q, 400 ng/mL), or IL-6 (100 ng/mL).

For CD44 blockade experiments in human iPSC-derived astrocytes, cells were pre-treated with an anti-CD44 monoclonal antibody for 24 hours, followed by co-treatment with the same antibody and the indicated stimuli for an additional 24 hours^16^.

To assess the impact of γ-secretase inhibition on OPN–CD44 interactions, primary mouse astrocytes and human iPSC-derived astrocytes were treated with 1 µM DAPT in the presence or absence of recombinant mouse or human OPN (1500 ng/mL) for 24 hours.

Upon completion of treatments, cells were fixed and subjected to immunofluorescence staining as described above. For BODIPY staining, following the final wash of the immunostaining procedure, coverslips were incubated with BODIPY (1 µM or 1:1,000 dilution) for 15 minutes at room temperature. Coverslips were then washed and mounted for imaging.

#### Gamma secretase inhibitor treatment in i3N and human iPSC derived astrocytes

i3N neurons and human iPSC-derived astrocytes were treated with the γ-secretase inhibitors DAPT (1 µM) or Compound E (100 nM) for 24 hours to assess their impact on APP processing. Following treatment, cells were washed with PBS and lysed in RIPA buffer supplemented with protease and phosphatase inhibitors. Lysates were prepared for western blotting for APP and β-actin.

#### In vitro lipid peroxidation assay

Primary mouse astrocytes and human iPSC-derived astrocytes were seeded onto PDL- or Matrigel-coated coverslips as described above. Cells were treated for 24 hours with recombinant OPN (1500 ng/mL) or with the positive-control oxidant Rotenone/Antimycin A (0.3 µM for 15 minutes). Following treatment, cells were incubated with lipid peroxidation sensor BODIPY 581/591 C11 (2 µM) for 40 minutes at 37 °C, then fixed with 4% paraformaldehyde for 20 minutes at room temperature. After fixation and washing with PBS, cells were stained with DAPI for imaging. Lipid peroxidation was quantified by measuring the intensity of oxidized versus reduced BODIPY C11 fluorescence using Fiji/ImageJ.

#### Astrocyte Aβ uptake assay

FAM-labeled Aβ was prepared as previously described^84^. Briefly, lyophilized peptide was dissolved in PBS. WT and CD44KO primary astrocytes were seeded on PDL-coated coverslips in 24-well plates at a density of 100,000 cells per well and cells were allowed to grow overnight. FAM-labeled Aβ (500 nM) was added to the culture medium, with or without recombinant mouse OPN (1500 ng/mL) and incubated for 24 hours. Following incubation, cells were washed thoroughly with PBS (3 times), fixed in 4%PFA for 20 minutes followed by washing and immunostaining with anti-GFAP antibody to identify astrocytes.

#### Proteomic sample preparation and analysis for in vitro samples

Primary astrocytes (WT and *Cd44*-KO) from three independent animals were treated overnight with mrOPN (1500 ng/mL) or vehicle. Harvested cells were lysed in 50 µL HEPES/SDS buffer and 10 units of Benzonase. The samples were briefly centrifuged and incubated at 37°C for 30 minutes while shaking at 1000 rpm. Following lysis, protein disulfide bonds were reduced by the addition of 2 µL of 120 mM TCEP (final concentration 5 mM). The samples were incubated at 55°C for 15 minutes with shaking, followed by alkylation performed by adding 2 µL of 500 mM MMTS (final concentration 20 mM) and incubating the samples at room temperature for 10 minutes. The total proteins from the cell lysate were further purified and enriched by SP3 (Single-Pot Solid-Phase-enhanced Sample Preparation)^85^ method on KingFisher instrument. One to one ratio mixture of Sera-Mag Speedbead carboxylate-modified [E3] magnetic particles (Cytiva, 65152105050250) and Sera-Mag carboxylate-modified [E7] magnetic particles (Cytiva, 24152105050250) was used for the protein purification and 1:50 ratio of Trypsin/Lys-C (Thermo Scientific, A40007) was used for overnight digestion of the proteins. The digested peptides were purified by C18 stage tips (CDS, Empore™ 6091). Tryptic peptides were analyzed on a Thermo Scientific Orbitrap Astral mass spectrometer operated in data-independent acquisition (DIA) mode with a total method duration of 7 minutes. Dried peptides were resuspended in 5% acetonitrile (ACN) containing 0.1% formic acid (FA), and 200 ng of each sample was injected twice as technical replicates. Peptide separation was performed on a Vanquish Neo UHPLC system using a 15 cm PepMap™ 100 C18 column (Thermo Scientific, Cat. No. 164534). Peptides were eluted with a 2–28% ACN gradient in 0.5% FA at a flow rate of 2.5 µL/min over 6 minutes, followed by a 1-minute column wash at 80% ACN.

For full MS (MS1) acquisition, data were collected from m/z 380–980 over the entire 0–7 min LC run. The Orbitrap analyzer was operated at a resolution of 240,000 (at m/z 200). The AGC conditions included a maximum injection time of 3ms, and one microscan was acquired. For MS2, DIA spectra were acquired across the 7-minute run using auto-defined isolation windows spanning m/z 380–980 with a 2 m/z window width and no overlap. Fragmentation was performed using higher-energy collisional dissociation (HCD) at 25% normalized collision energy, and fragments were detected using the ASTRAL analyzer. DIA scans were collected across m/z 150–2000 with a maximum injection time of 5ms and one microscan. All data were acquired in positive polarity and centroid mode.

MSFragger-DIA was used to directly search the DIA data. The search results were processed using MSBooster for deep learning-based score calculation, Percolator33 for rescoring and posterior error probability calculation, ProteinProphet34 for protein inference, Philosopher55 for FDR filtering, and EasyPQP for spectral library building. The peptide ions in the spectral library were filtered with 1% global peptide and protein FDR. The resulting library was passed to DIA-NN23 to extract and quantify precursors, peptides, and proteins from the DIA data.

Spectra were searched using FragPipe (v23.0) MSFragger 4.3^86^ with DIA_SpecLib_Quant workflow against Uniprot database (UPR_Mouse_10090 accessed on 25.08.23). One missed cleavage of tryptic peptide was allowed. MSBooster^87^ and Percolator were used for data rescoring and PSM validation. DIA-NN (1.8.2)^88^ was used for label-free quantification.

#### Astrocyte-microglia co-culture

For primary microglial culture, mouse microglial cells from WT mixed glial cell culture were selected with CD11b microbeads according to the manufacturer’s instructions (130-093-634, Miltenyi Biotec). Enriched microglia were plated in 24-well PDL-coated plates in complete DMEM media with 10% FBS and 1% Pen/Strep along with WT and CD44KO astrocytes at a ratio of 1:2. Co-cultured cells were allowed to grow for 48 hours before cell lysis and RNA extraction for analysis.

#### RNA extraction, reverse transcription, and qPCR

Total RNA was isolated from cultured astrocytes and microglia derived from three independent neonatal mice using RLT buffer (Qiagen) supplemented with 1% β-mercaptoethanol, followed by purification with the RNeasy Mini Kit (Qiagen) following manufacturer’s protocol. cDNA was synthesized from 1 μg of RNA using iScript Reverse Transcription Supermix (Bio-Rad). Quantitative real-time PCR (qRT-PCR) was performed using iTaq Universal SYBR Green Supermix on a CFX384 Touch Real-Time PCR Detection System (Bio-Rad). Gene expressions were normalized to GAPDH and analyzed using CFX Manager software.

#### Behavioral assessment

Behavioral assays were performed as previously described^41^. For novel object recognition (NOR), mice were placed in a clear Plexiglass arena (22 × 44 cm). On the training day, animals freely explored two identical LEGO objects for 10 minutes. Twenty-four hours later, one familiar object was replaced with a novel object of similar size and volume but differing in shape and color. Mice were allowed to explore for 10 minutes, and object exploration time was recorded using ANY-maze software (Stoelting). Exploration was defined as the nose being within 2 cm of an object. Novel object preference was calculated as the percentage of total exploration time spent on the novel object.

Fear conditioning was performed using a standard three-phase protocol^89^. During training, mice were placed in the conditioning chamber and allowed to explore for 2 minutes before receiving two tone–shock pairings (80 dB, 5 kHz tone for 30 s followed by a 0.8 mA foot shock for 2 s). 24 hours later, contextual memory was assessed by returning mice to the same chamber for 5 minutes without tone or shock. One hour later, cued memory was tested in a novel chamber differing in shape, lighting, flooring, and scent. After 3 minutes of exploration, the tone was presented for 3 minutes. Freezing behavior was quantified using FreezeFrame4 software (Actimetrics) and expressed as a percentage of total time.

### QUANTIFICATION AND STATISTICAL ANALYSIS

#### General immunofluorescence quantification

Images were acquired on Leica confocal microscope using identical microscope settings for an experiment across all samples to ensure quantitative comparability. Percent area fluorescence and mean intensity were quantified using consistent thresholding parameters applied across all images within an experiment. All images were quantified using Fiji/ImageJ unless otherwise noted.

#### In vitro Aβ Internalization

Z-stack images were acquired using a Leica confocal microscope with a 0.7 μm step size. The IMARIS Co-localization module (Oxford Instruments) was used to quantify internalized Aβ within GFAP-positive astrocytes.

#### In vivo synaptic colocalization

Synaptic colocalization was quantified as previously described^41,90^. Images were acquired using a Leica confocal microscope with a 63× oil immersion objective and 6× digital zoom. Z-stacks were collected at 0.2 μm intervals over a 10 μm depth. Pre- and postsynaptic puncta were detected using the Spots function in IMARIS (Oxford Instruments), with manual adjustment for optimal detection in each channel. The total number of puncta was recorded, and colocalization was determined using the IMARIS Co-localize Spots MATLAB plugin (MathWorks). Puncta were considered colocalized if their centers were within 200 nm.

#### Gliosis and plaque characterization

Coronal brain sections (30 μm) were imaged using a Leica confocal microscope with a 63× oil objective and 1.5× digital zoom. Z-stacks spanning the full section thickness were acquired at 1 μm intervals. Fluorescent signal area was quantified using Fiji/ImageJ (NIH), and 3D surface rendering of microglia and plaque channels was performed using the Surface function in IMARIS (Oxford Instruments). Six images per animal were analyzed from three males and three females.

#### Statistics

All statistical analysis was performed using GraphPad Prism software v8.0.2. All data are presented as mean ± SEM. Unless otherwise noted, all group comparisons were made using 1-way ANOVA with Tukey’s correction, and all pairwise comparisons by 2-sided Student’s t tests, depending on experimental design. For all tests, *p* values less than 0.05 were considered significant, and those over 0.05 were considered nonsignificant: \**p* < 0.05, \*\**p* < 0.01, \*\*\**p* < 0.001, \*\*\*\**p* < 0.0001.

#### Study approval

All animal procedures were performed in accordance with the NIH Guide for the Care and Use of Laboratory Animals (National Academies Press, 2011) and with the approval of the Baylor College of Medicine Institutional Animal Care and Use Committee.

---

**Figure S1.**
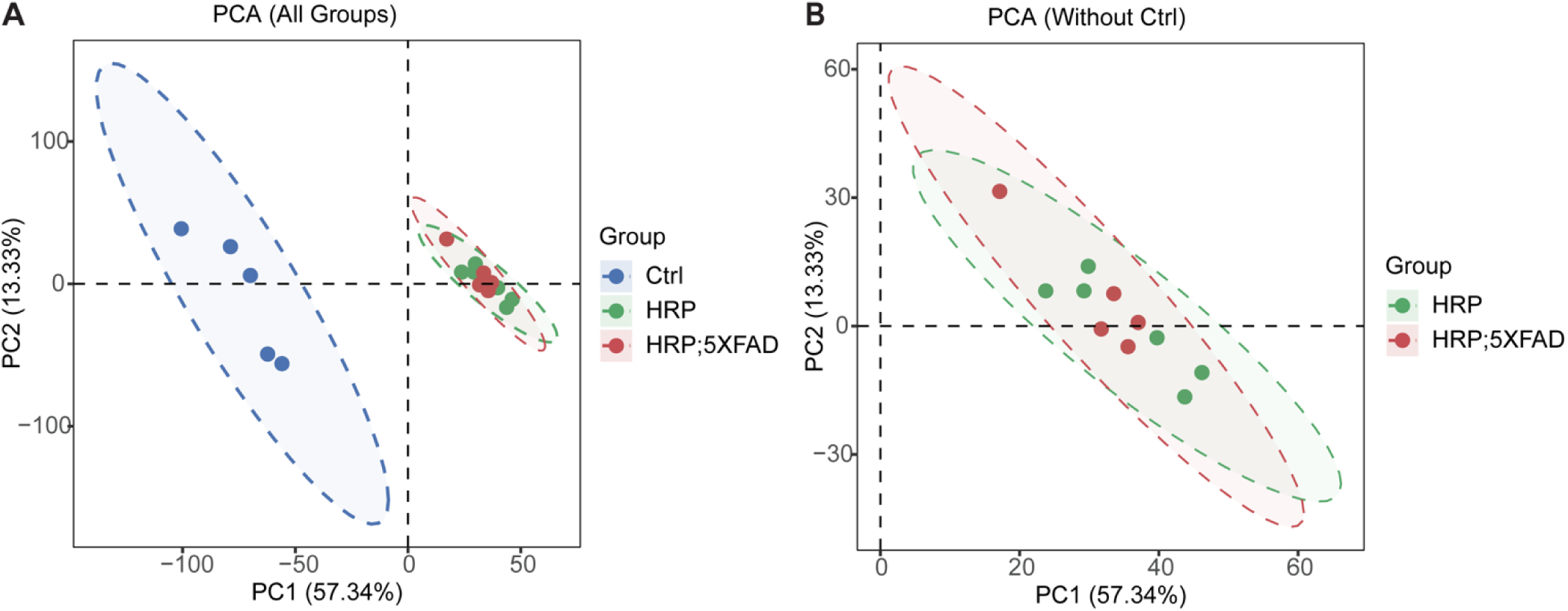
(associated with Figure 1). PCA analysis of iPEEL proteomics. **(A)** Principal component analysis (PCA) including all groups (Ctrl, HRP, HRP;5xFAD) showing clear separation of control samples from the HRP-labeled groups along PC1. **(B)** PCA of only HRP and HRP;5xFAD samples showing tighter clustering and spatial separation between genotypes. Each dot represents one biological replicate; shaded ellipses indicate 95% confidence intervals.

**Figure S2.**
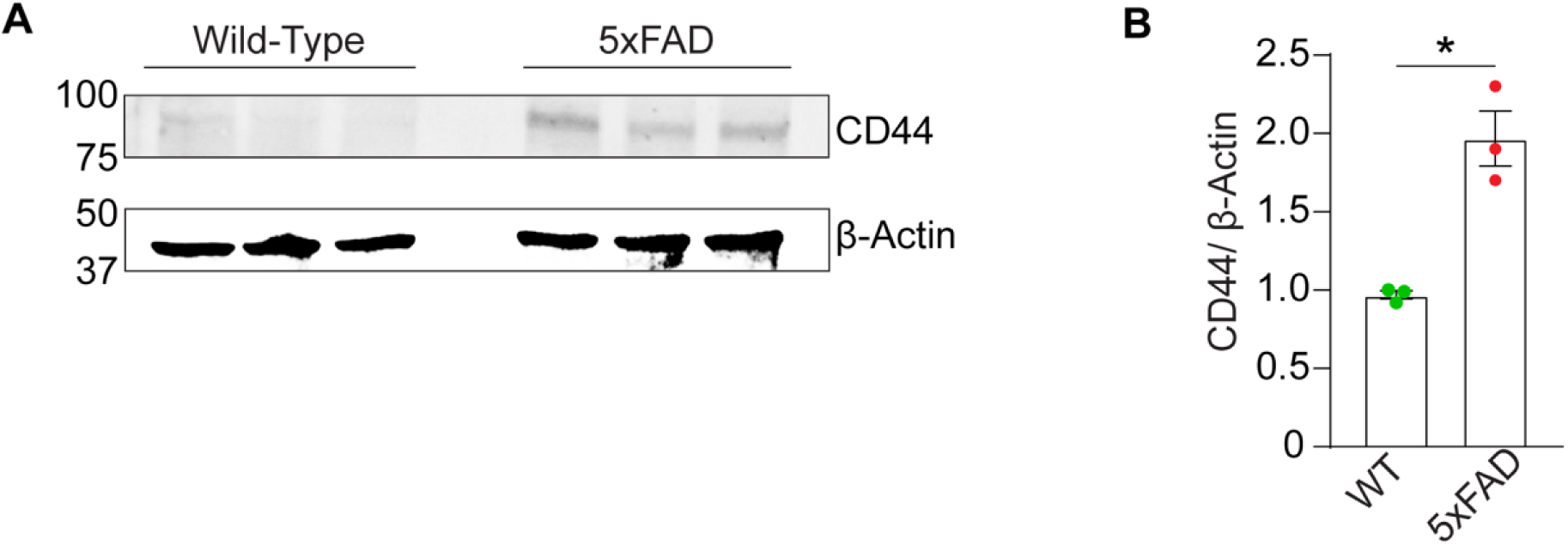
(associated with Figure 2). Western blot analysis for CD44 expression in mouse hippocampus. **(A)** Western blot and **(B)** quantification of CD44 protein levels in hippocampal lysates from WT and 5xFAD mice (normalized to β-actin). Data is presented as mean ± SEM; n=3 mice per group. Statistical analysis by Student’s *t* test. \**p* < 0.05.

**Figure S3.**
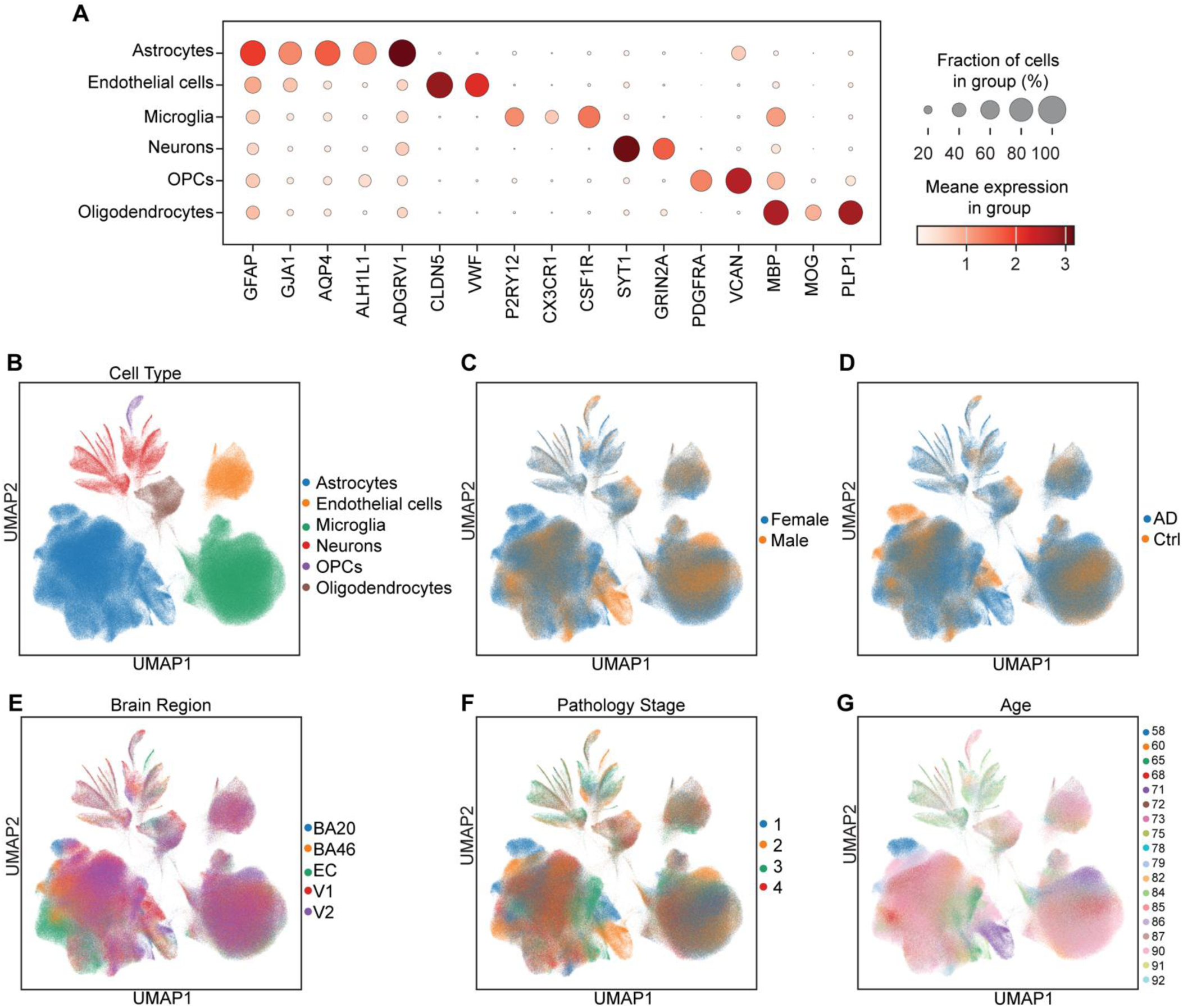
(associated with Figure 2). snRNAseq analysis of astrocyte nuclei. **(A)** Dot plot showing the expression of canonical marker genes across major cell types from 1,079,333 nuclei from 32 donors. **(B)** UMAP of the same data set annotated by major cell types. **(C-G)** UMAPS showing the cell distribution by sex **(C)**, diagnosis (AD vs Control) **(D)**, brain region **(E)**, pathology stage **(F)**, and donor’s age rage (58-92 years) **(G).**

**Figure S4.**
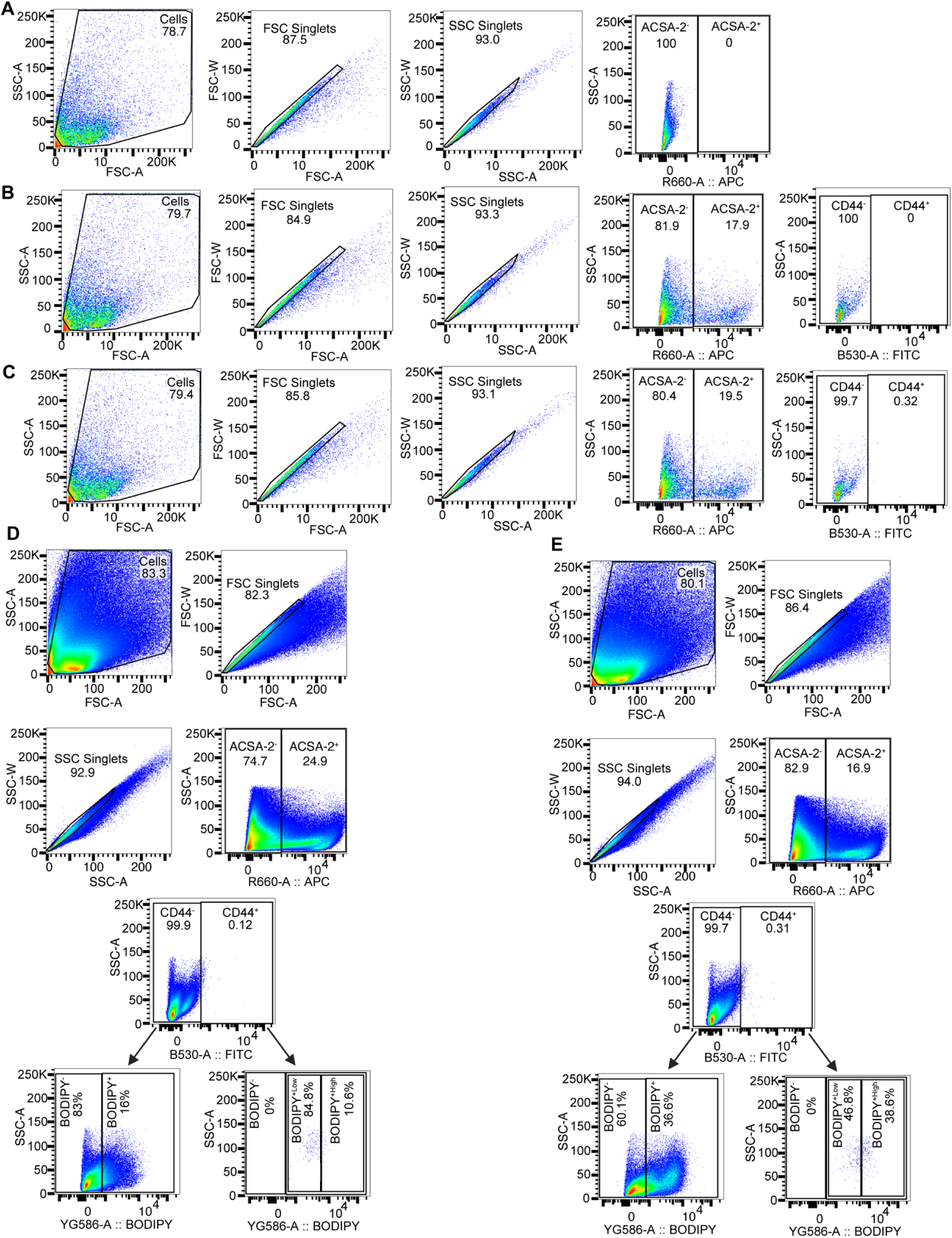
(associated with Figure 3). Gating strategy and representative plots for astrocyte lipid content analysis. **(A-C)** Flow cytometry plots showing the gating strategy for astrocyte selection from control unstained sample **(A),** single color control for ACSA2 antibody **(B),** and single-color control for CD44 antibody **(C)**. **(D and E)** Representative flow cytometry plots showing sequential gating strategy for astrocyte selection, followed by CD44 expression assessment and BODIPY intensity assessment from 12-month-old WT **(D)** and 5xFAD **(E)** mice.

**Figure S5.**
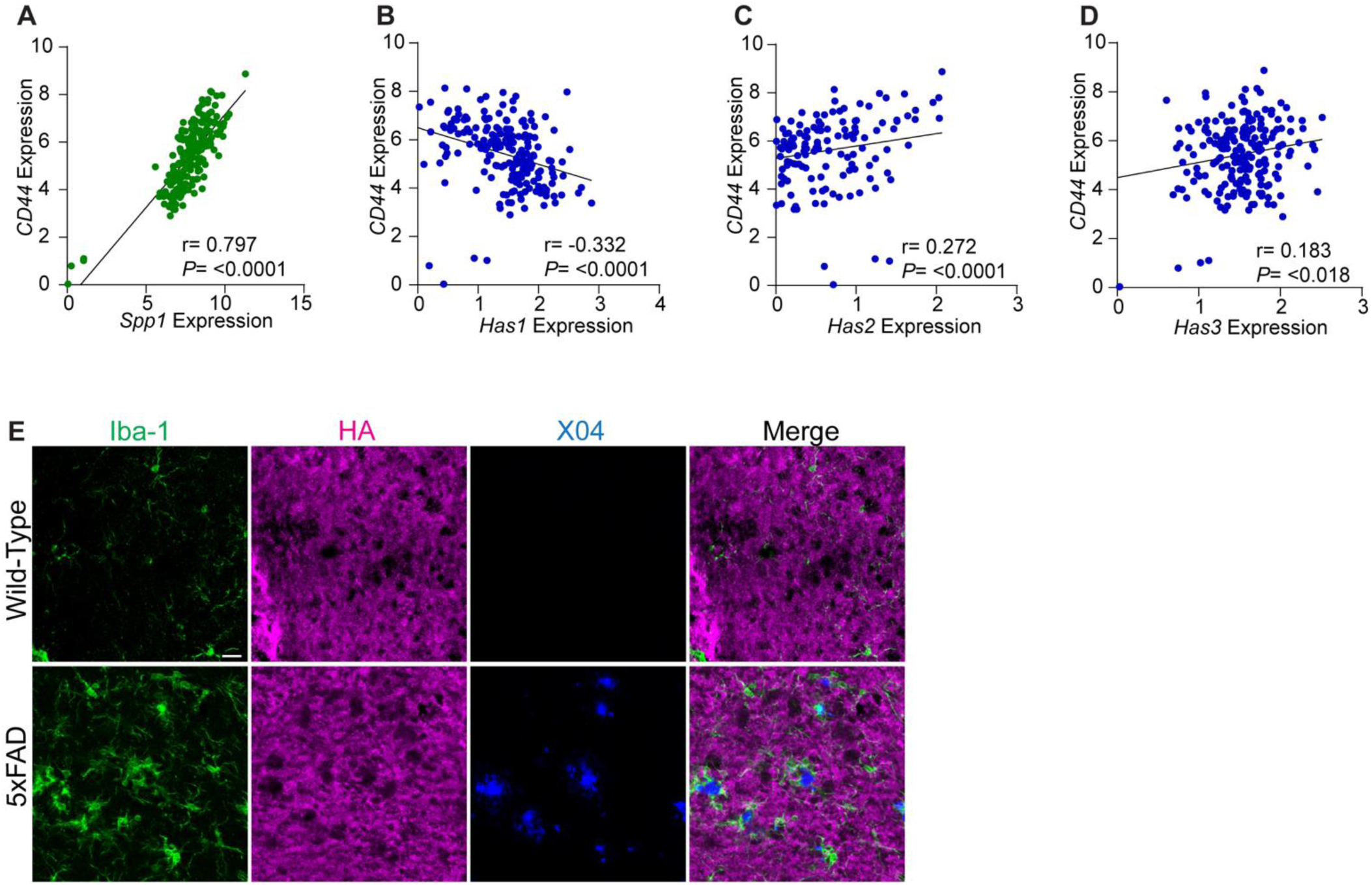
(associated with Figure 3). Positive correlation between CD44 and OPN, but not HA. **(A-D)** Correlation analysis between *CD44* expression and *SPP1* (**A**), *HAS1* **(B)**, *HAS2* **(C)**, and *HAS3* **(D)**. Pearson correlation coefficients and *P* values are indicated. **(E)** Representative images of hippocampal sections from 6-month-old WT and 5xFAD mice stained with Iba-1, HA, and methoxy-X04, showing comparable HA staining with and without Aβ plaques.

**Figure S6.**
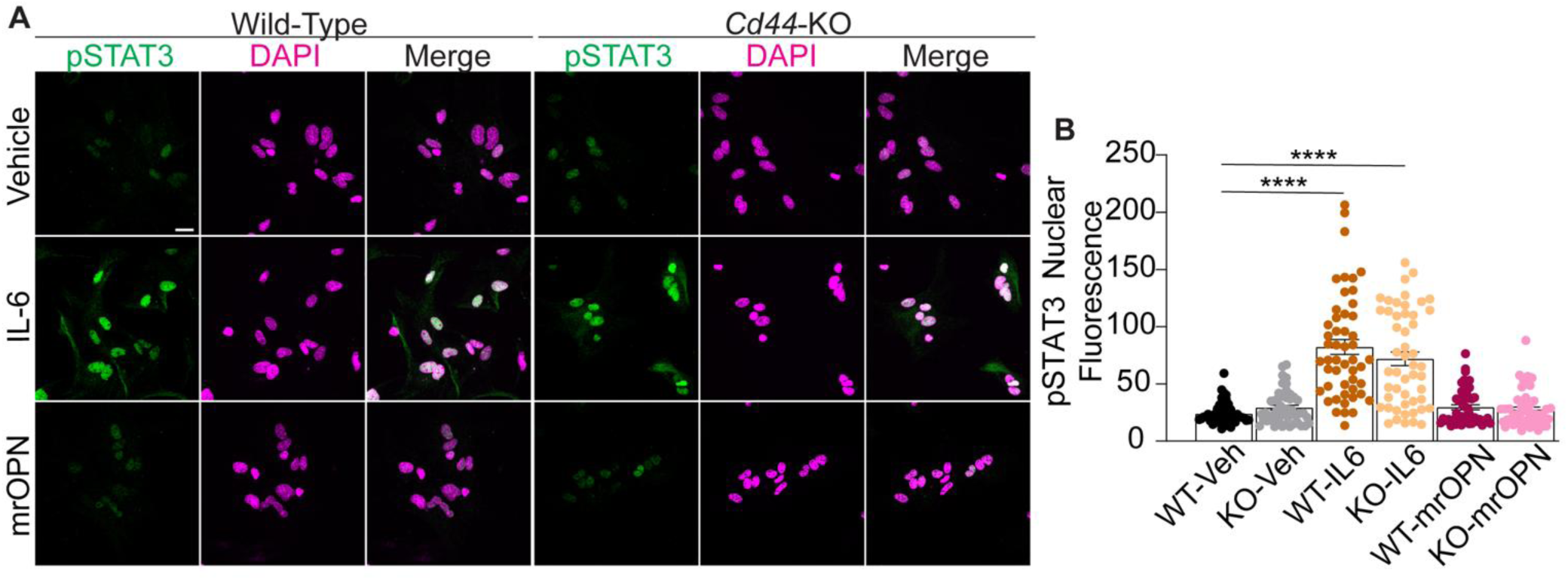
(associated with Figure 3). OPN-CD44 signaling does not induce STAT3 activation. **(A)** Representative images of primary mouse astrocytes from WT and *Cd44*-KO mice treated with vehicle, IL-6 or mrOPN, stained for phospho-STAT3 (pSTAT3) and DAPI. Scale bar, 20 µm. **(B)** Quantification of nuclear pSTAT3 fluorescence showing robust activation by IL-6 but not by mrOPN. Each dot represents one nucleus with a total of 50 nuclei analyzed per condition. Data is presented as mean ± SEM. Statistical analyses were performed using One-way ANOVA with Tukey’s multiple comparison test. \*\*\*\**p* < 0.0001.

**Figure S7.**
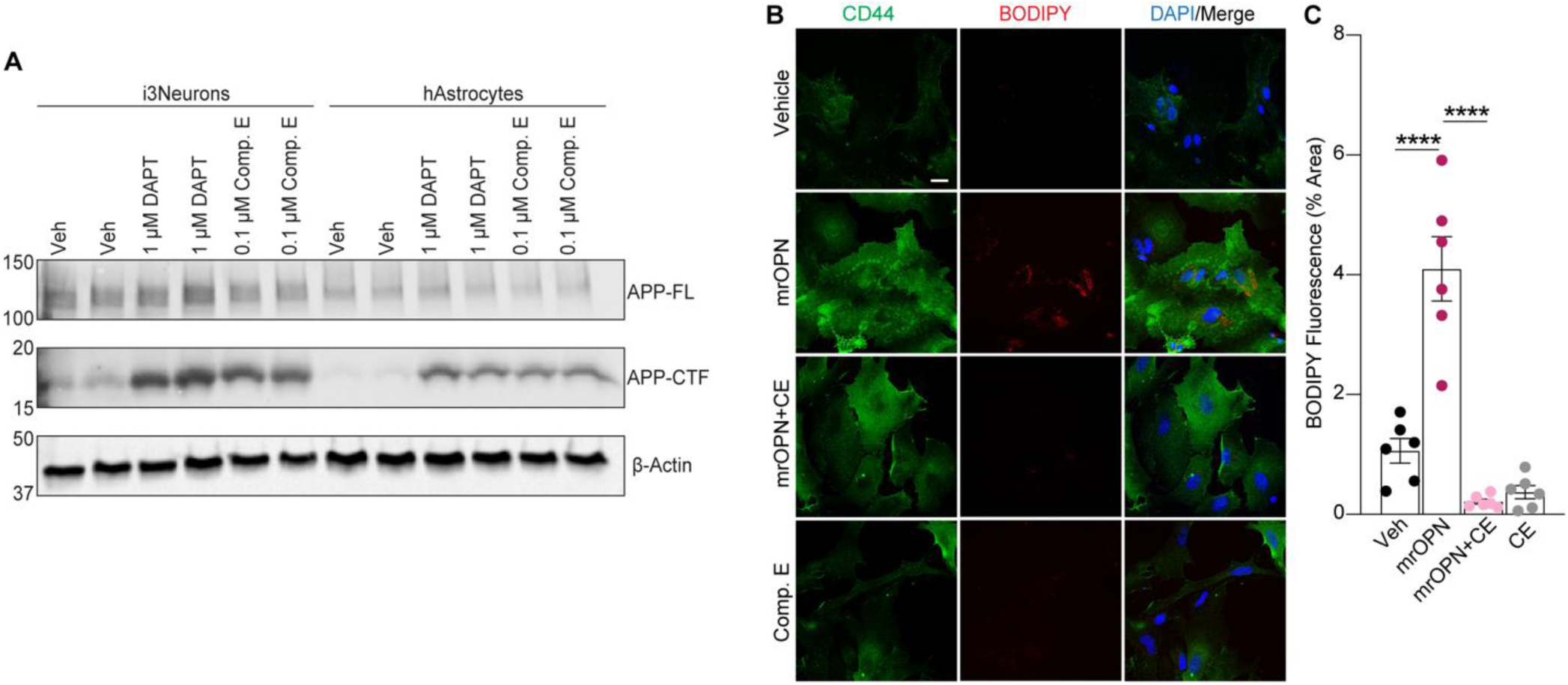
(associated with Figure 4). γ-secretase inhibition blocks OPN-induced lipid accumulation in astrocytes. **(A)** Immunoblot analysis of human iPSC-derived neurons (i3Neurons) and human iPSC derived astrocytes (hAstrocytes) treated with vehicle, DAPT (1 µM) or Compound E (CE) (0.1 µM). Both γ-secretase inhibitors caused accumulation of APP-CTF. β-Actin is the loading control. **(B)** Representative confocal images of primary wild-type mouse astrocytes treated with vehicle, mrOPN, mrOPN + CE, or CE alone. Cells were stained for CD44 (green), BODIPY (red), and DAPI (blue). **(C)** Quantification of BODIPY-positive area (% area) in (B). n = 3 mice per condition; 6 images per mouse. Each dot represents the average value of three images from one animal. Data are presented as mean ± SEM. Statistical analysis was performed using one-way ANOVA with Tukey’s multiple comparison test. \*\*\*\**p* < 0.0001. Scale bars: 20 µm.

**Figure S8.**
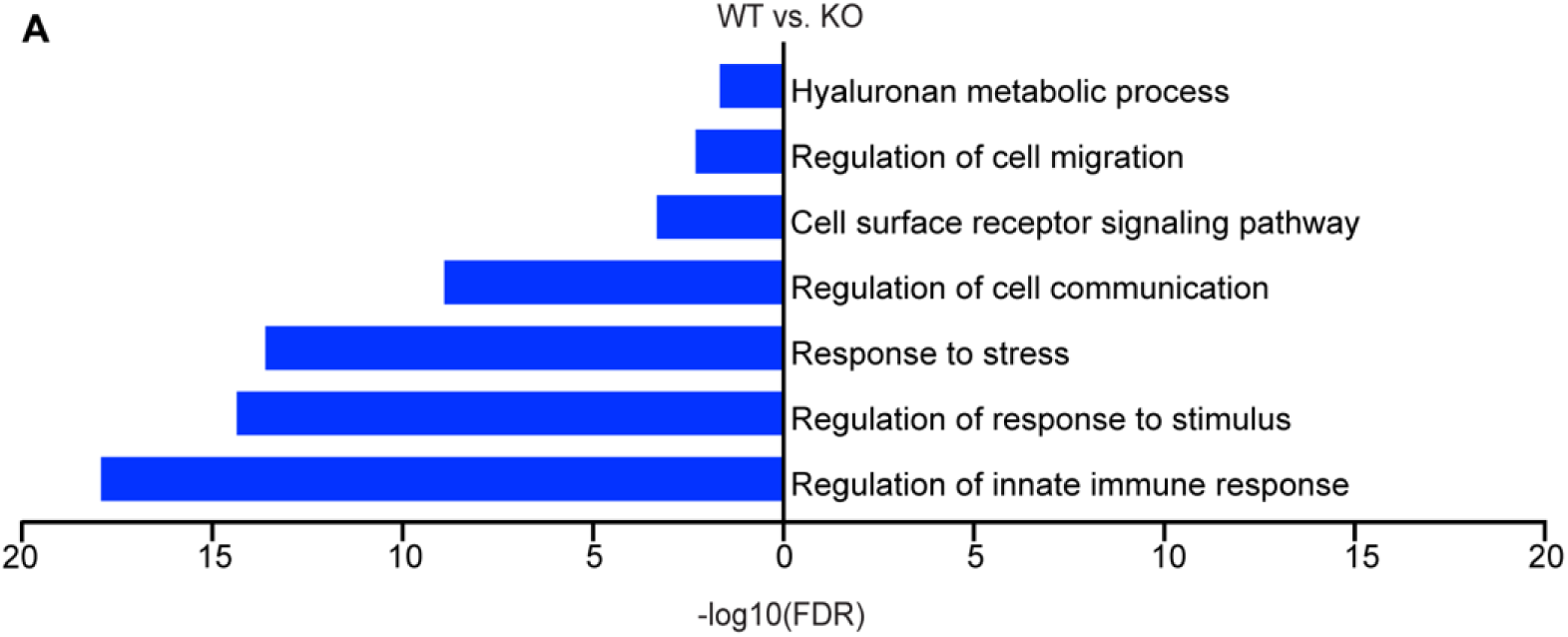
(associated with. Figure 4**). Suppressed pathways in *Cd44*-KO cells.** GO term enrichment analysis comparing WT and *Cd44* -KO astrocytes, highlighting significantly downregulated pathways in blue bars.

**Figure S9.**
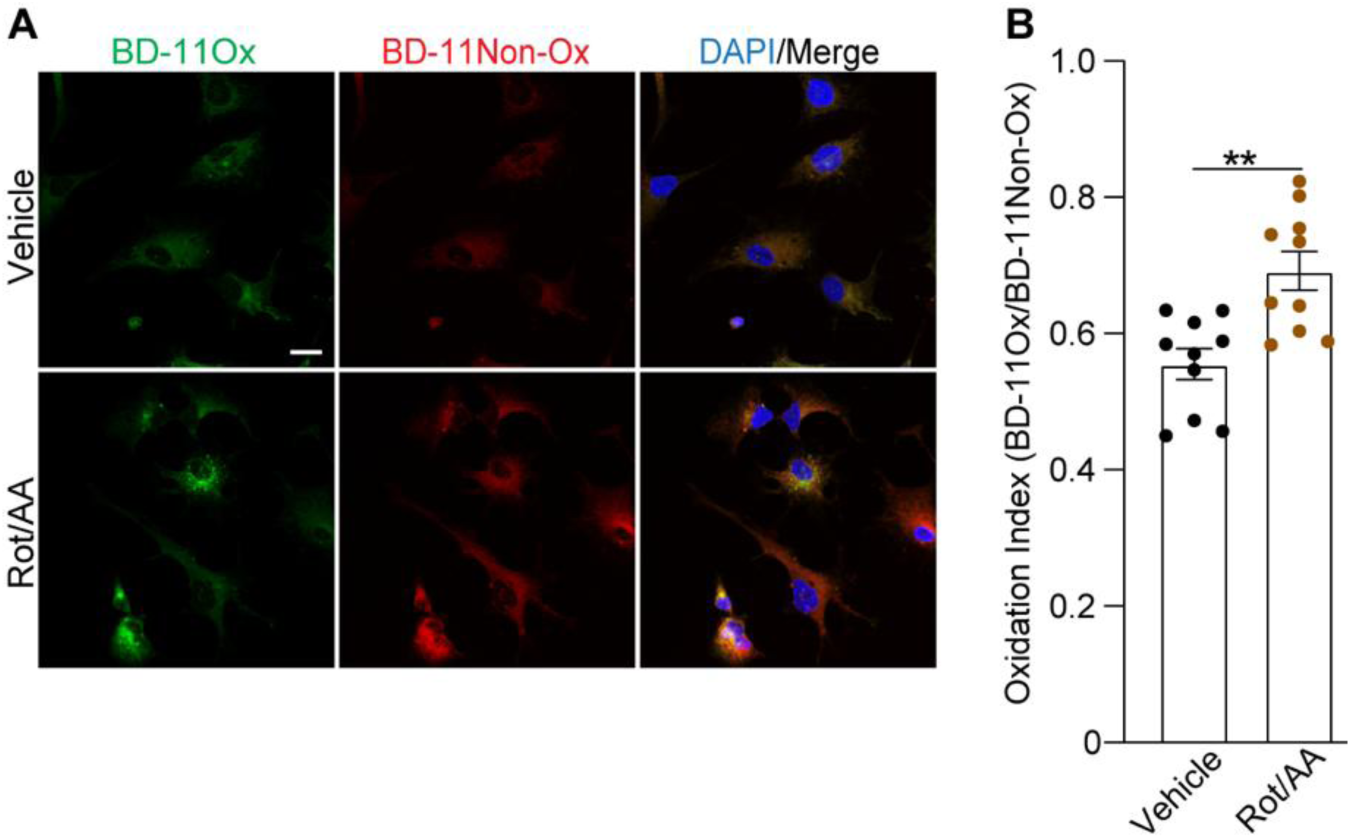
(associated with Figure 4). Rotenone antimycin A (Rot/AA) treatment induces lipid peroxidation comparable to hrOPN treatment. **(A)** Representative images from human iPSC-derived astrocytes treated with vehicle or Rot/AA for 24 hrs and stained for BODIPY C11 (green:oxidized, red:non oxidized) and DAPI (blue). Scale bar: 20 μm. **(B)** Quantification of lipid peroxidation index (BD-11Ox/BD-11Non-Ox). Each dot represents one field of view. n= ∼100 cells per condition. Data is presented as mean ± SEM. Statistical analysis was performed using unpaired 2-tailed Student’s *t* test \*\**p* < 0.01. ns, not significant.

**Figure S10.**
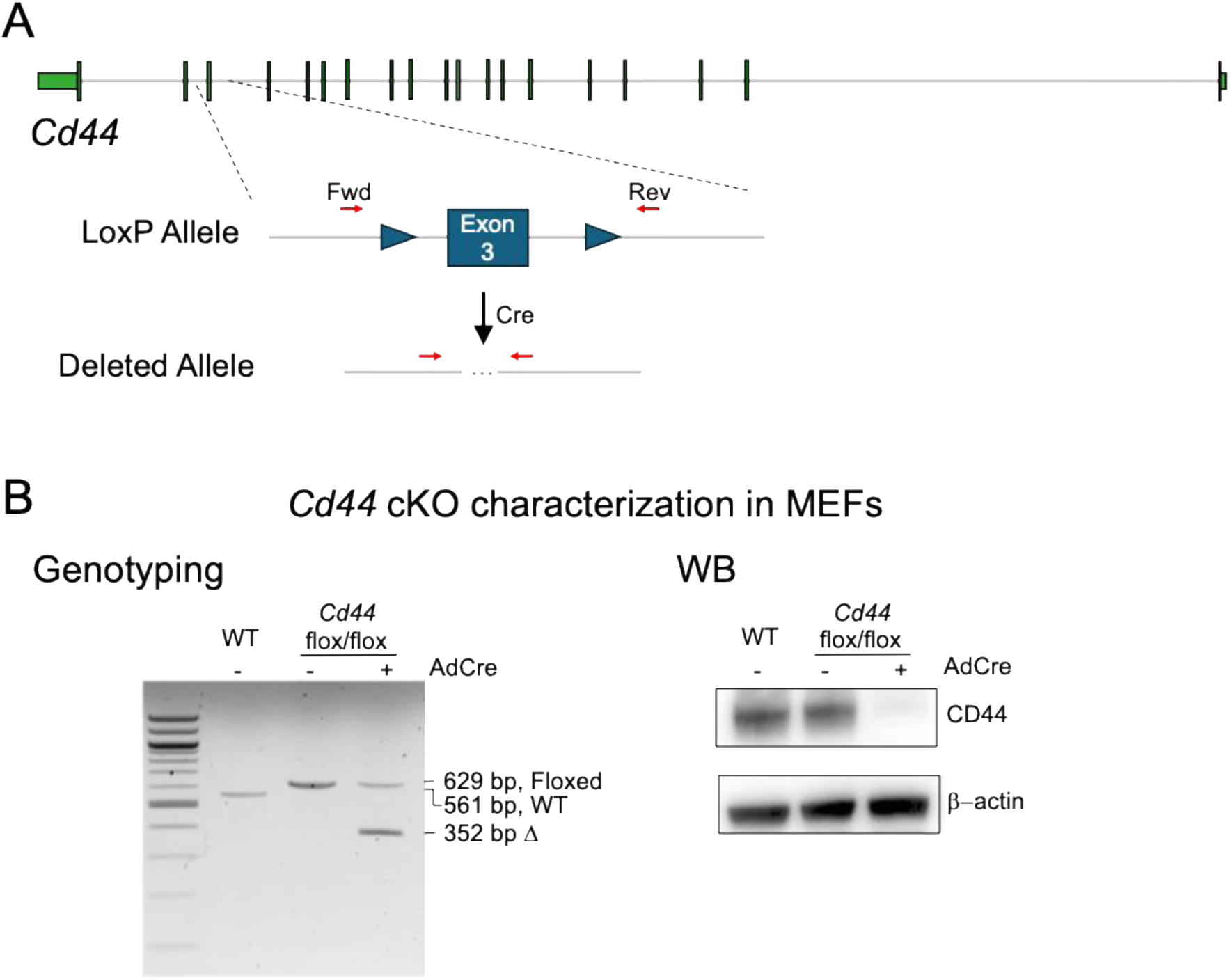
(associated with Figure 5). Generation and characterization of *Cd44* floxed mice. **(A)** A diagram of *Cd44* conditional allele. CRISPR-guided insertion of LoxP cassettes were inserted upstream and downstream of *Cd44* exon 3. Cre recombination excises exon 3, resulting in loss of CD44 protein expression. Red arrows show locations of primers that flank exon 3 and were used for genotyping in Panel B. **(B)** Mouse embryonic fibroblasts (MEFs) isolated from WT and *Cd44* flox/flox mice were treated with (+) and without (-) an AdenoCre-recombinase. *Bottom Left,* PCR analysis of the *Cd44* allele using primers shown in Panel A (red arrows). The PCR products of 629, 561, 352 base pairs correspond to amplified DNA products from *Cd44* flox/flox, WT *Cd44*, and Cre-induced exon 3 deleted *Cd44*, respectively. *Bottom right,* Western blot (WB) analysis showing normal CD44 level in *Cd44* flox/flox mice without Cre but depletion of CD44 in Cre-induced MEFs. β-action serves as a loading control.

**Figure S11.**
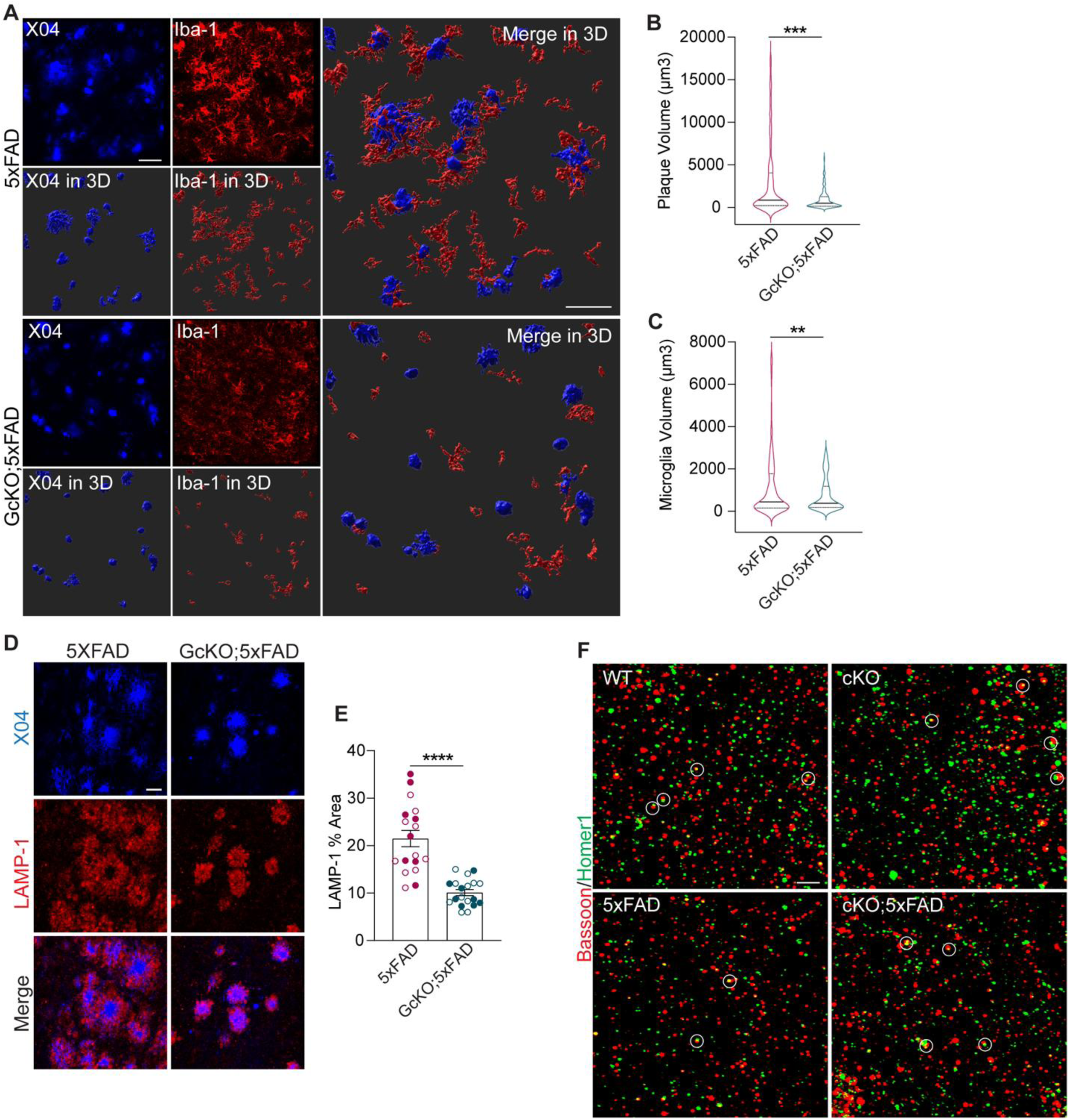
(associated with Figure 6). Astrocytic *Cd44* deletion by the GFAP^Cre^ driver ameliorates amyloid burden and glial reactivity. **(A)** Representative images of Aβ plaques and microglia labeled with X04 and Iba-1 respectively and their 3D surface reconstruction from the of 6 months-old 5xFAD and GcKO;5xFAD mice (n=2/group; 1 male, 1 female). **(B, C)** Violin plots quantifying plaque volume (X04⁺) **(B)** and associated microglial volume (Iba1⁺) per plaque **(C)** across 202 plaques (5xFAD) and 170 plaques (GcKO;5xFAD). **(D)** Representative images from tissues stained with X04 and LAMP-1 to label dystrophic neurites in 6-month-old 5xFAD and GcKO;5xFAD animals. **(E)** Quantification of LAMP-1 percent area corresponding to images in **(D)**. **(F)** Representative images of DG region stained for presynaptic (Bassoon) and postsynaptic (Homer1) markers in WT, cKO, 5xFAD and cKO;5xFAD mice, with quantifications displayed in main Figs. 6, K-M. Graphs display mean ± SEM. Filled circles denotes male, open circles female. n=2 mice per group. For each mouse, six images were analyzed, derived from two intendent brain sections (three images per section). Each dot represents one image. Statistics: unpaired 2-tailed Student’s *t* test. *\*\*p* < 0.01, \*\*\**p* < 0.001, \*\*\*\**p* < 0.0001. Scale bars: 20 µm (A, D), 3 µm (F).

**Figure S12.**
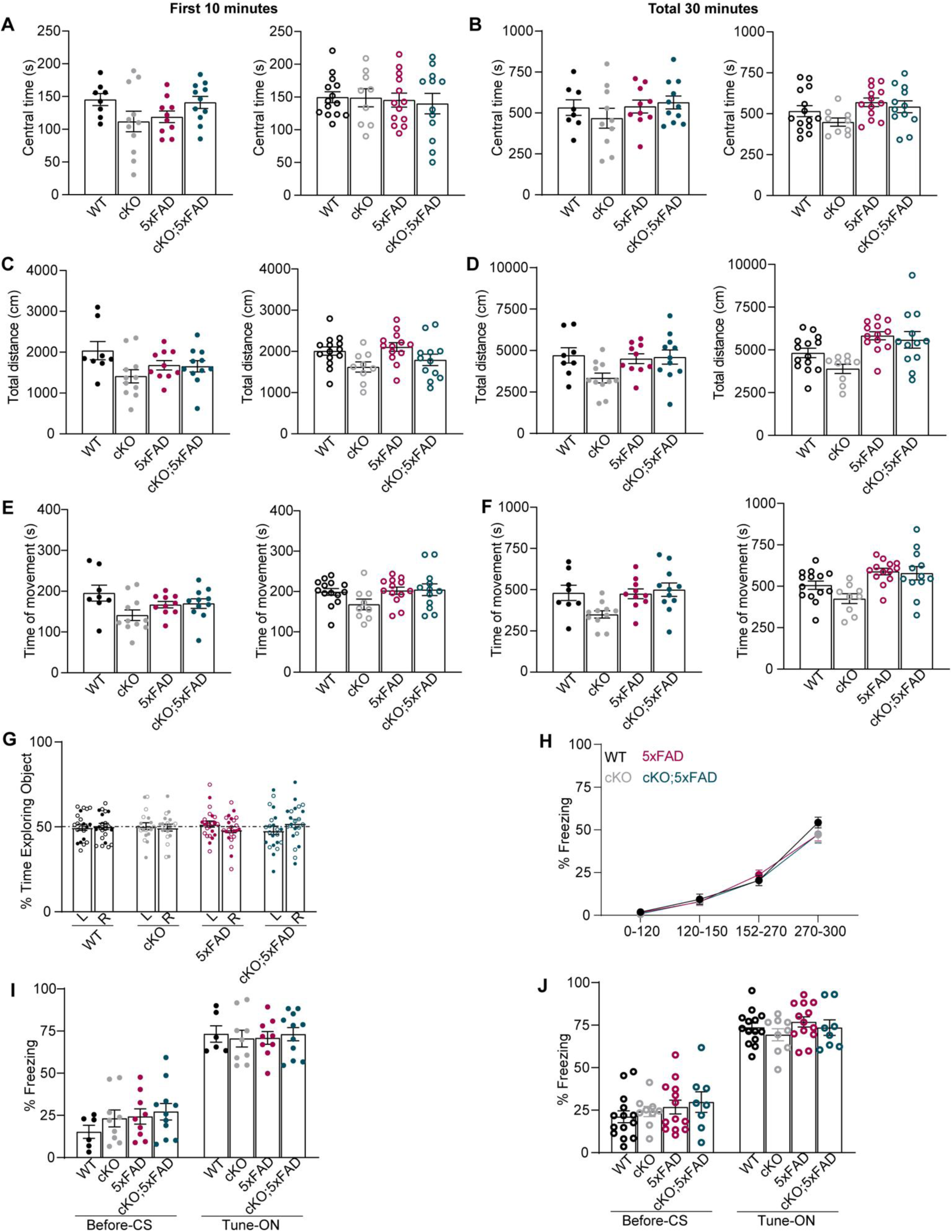
(associated with Figure 6). Behavioral analyses in WT, cKO, 5xFAD, and cKO;5xFAD mice. **(A-F)** Open field test showing the time spent in the central zone during the first 10 minutes **(A)** and across the total 30 minutes **(B).** total distance traveled during the first 10 minutes **(C)** and across the total 30 minutes **(D),** and total movement time during the first 10 minutes **(E)** and across the total 30 minutes **(F)**. Group sizes: WT (n = 8♂, 14♀), Cd44 cKO (n = 11♂, 9♀), 5xFAD (n = 10♂, 13♀), and cKO;5xFAD (n = 11♂, 12♀). **(G)** NOR training phase showing similar percent time spent on identical objects placed in left (L) versus right (R) arms for each group. Group sizes: WT (n = 8♂, 14♀), cKO (n = 11♂, 8♀), 5xFAD (n = 10♂, 13♀), and cKO;5xFAD (n = 10♂, 12♀). **(H)** Time course of freezing behavior across groups. **(I and J)** Percentage of freezing before conditioned stimulus (CS) and during tune presentations (Tune-On). Group sizes: WT (n = 6♂, 14♀), cKO (n = 9♂, 9♀), 5xFAD (n = 9♂, 13♀), and cKO;5xFAD (n = 11♂, 8♀). Graphs display mean ± SEM. Filled circles denotes male, open circles female. Statistical analyses were performed using one-way ANOVA with Tukey’s multiple comparison test. All non-significant.

**Table S1.**
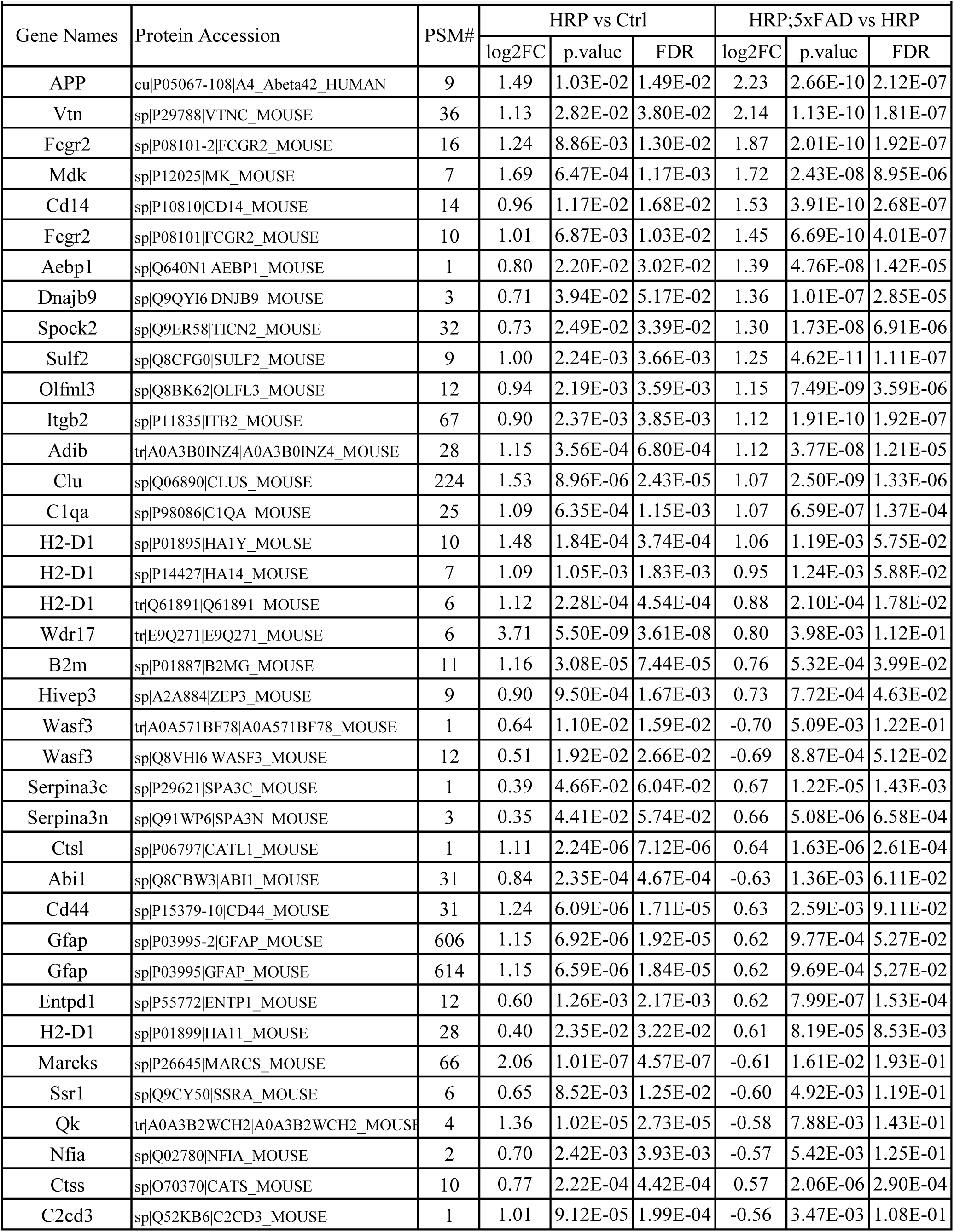
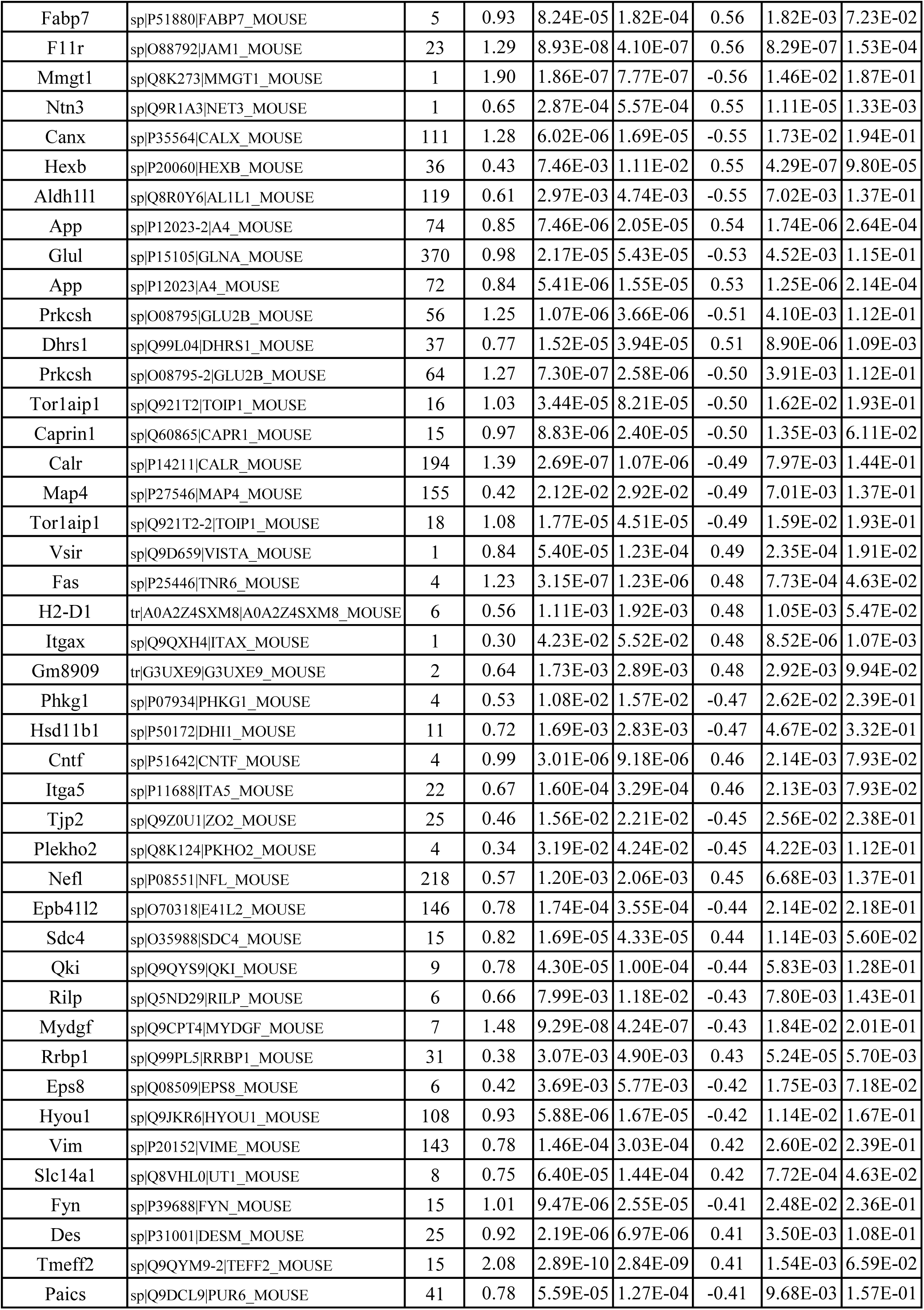
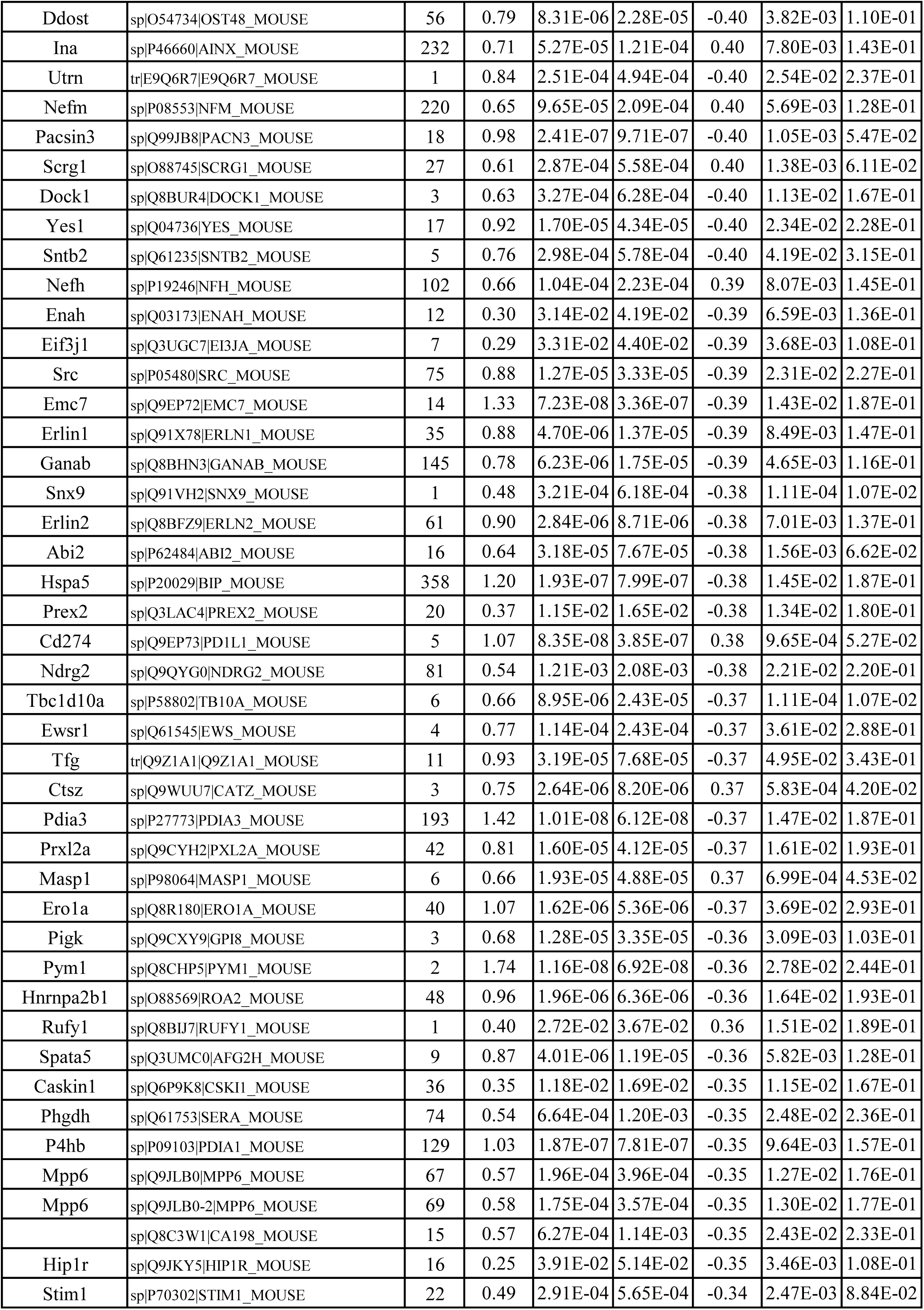
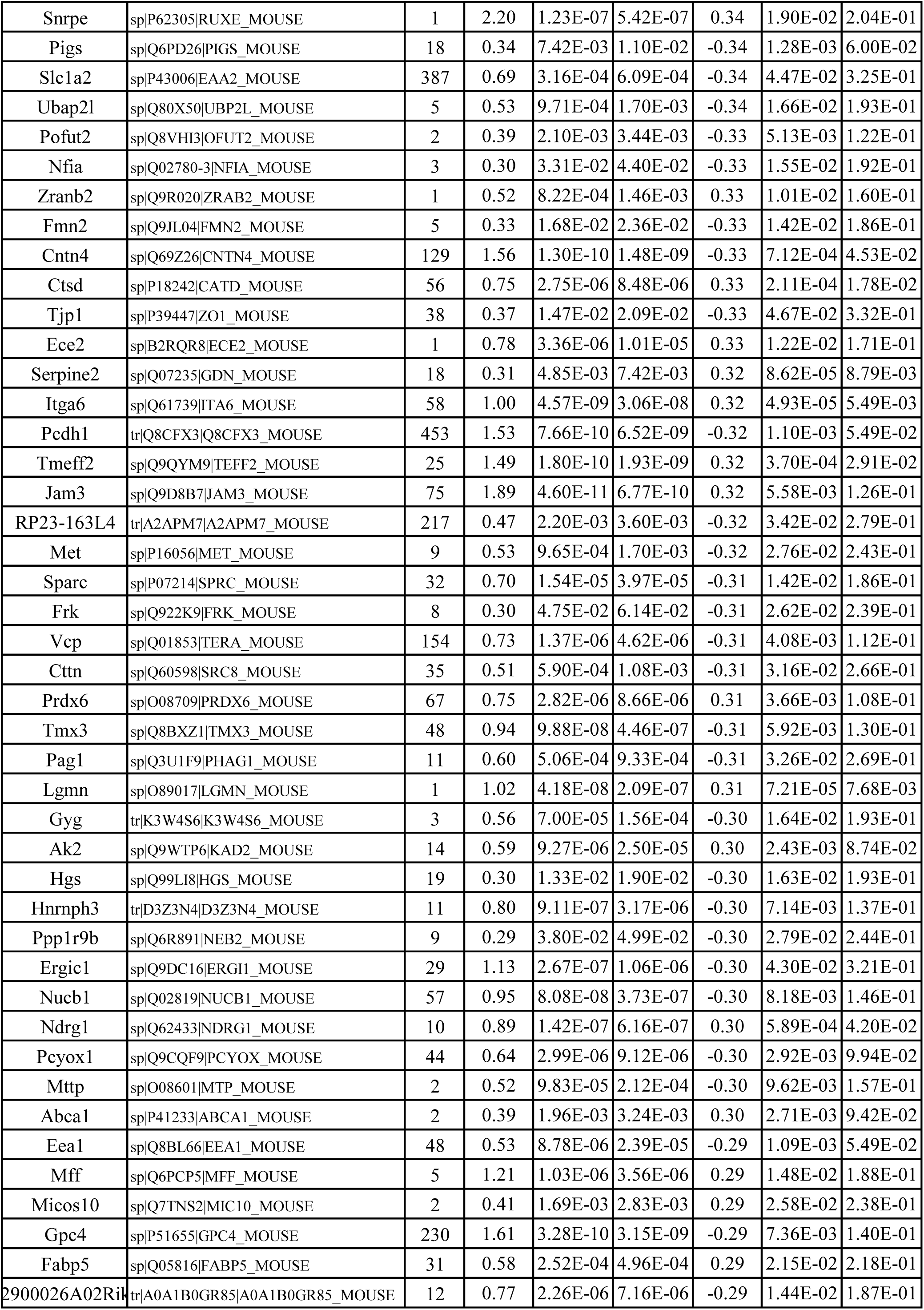
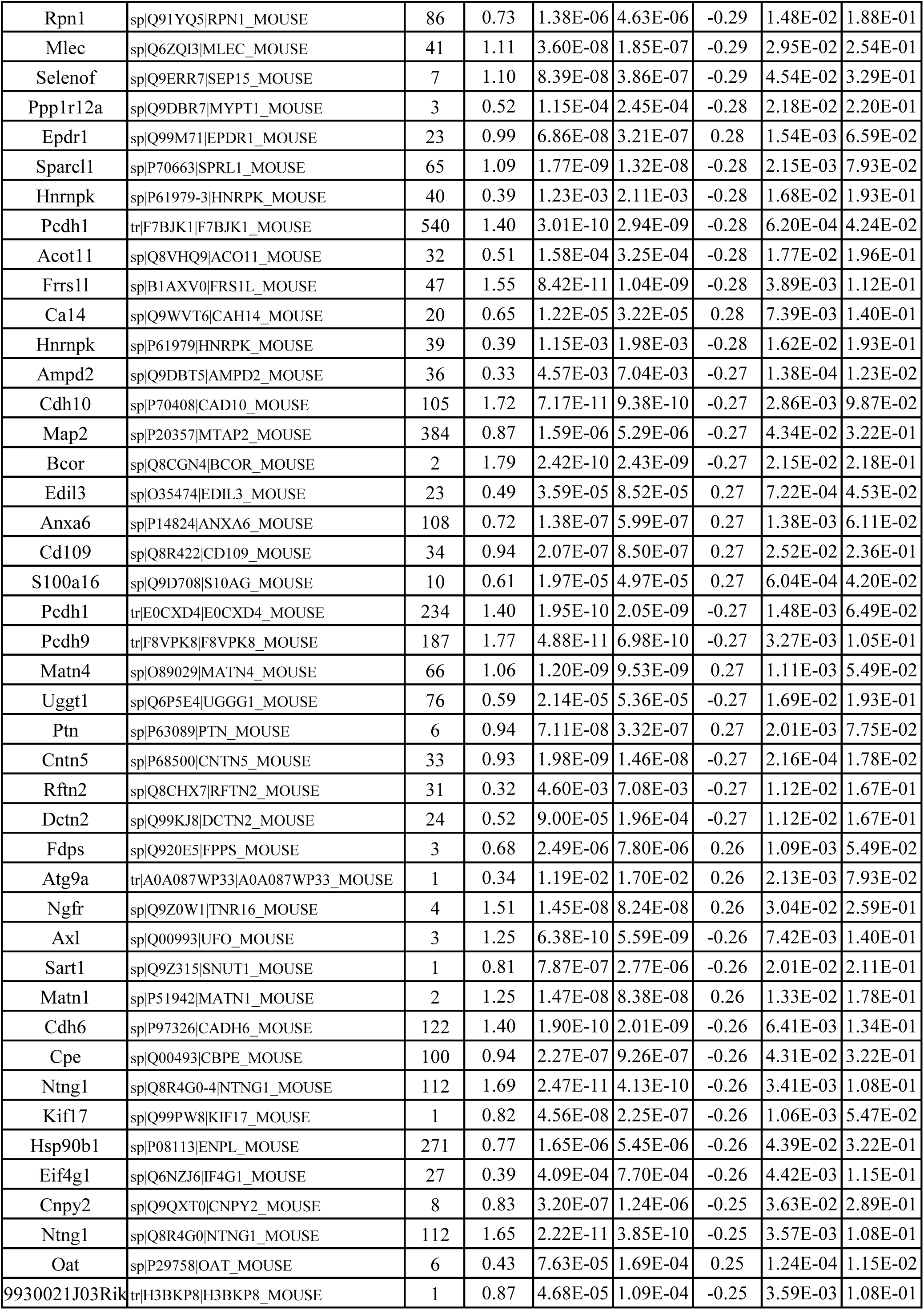
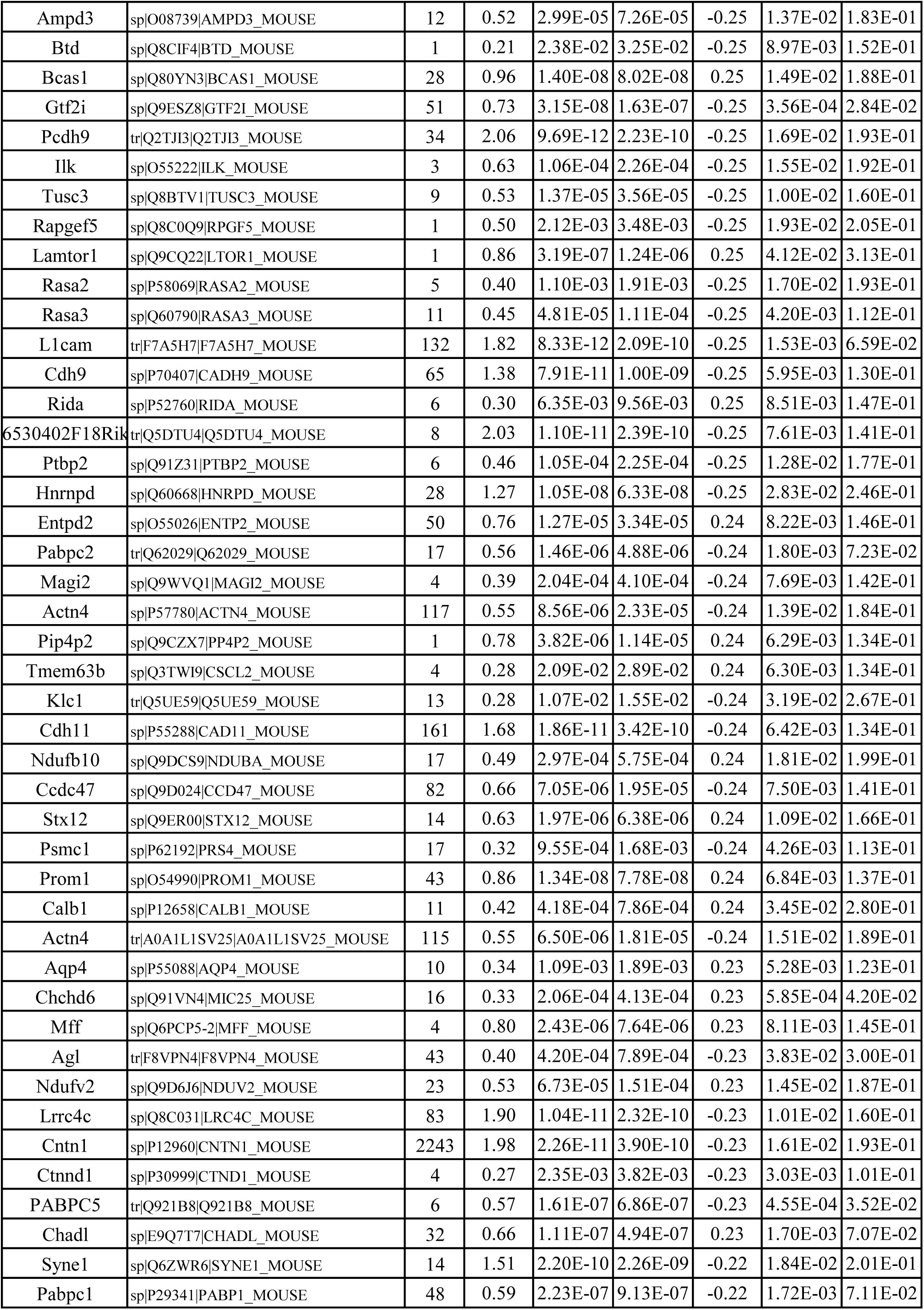
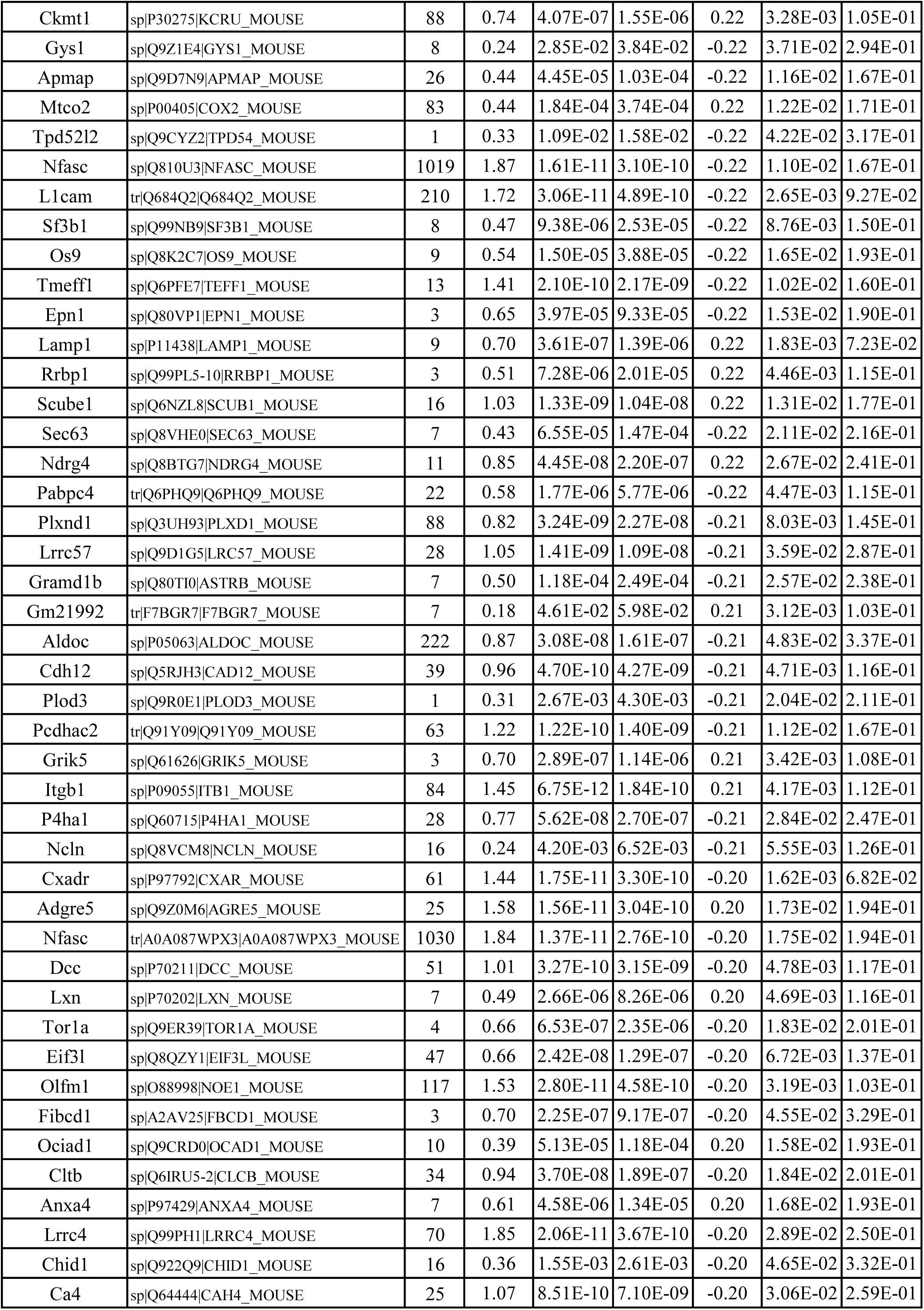
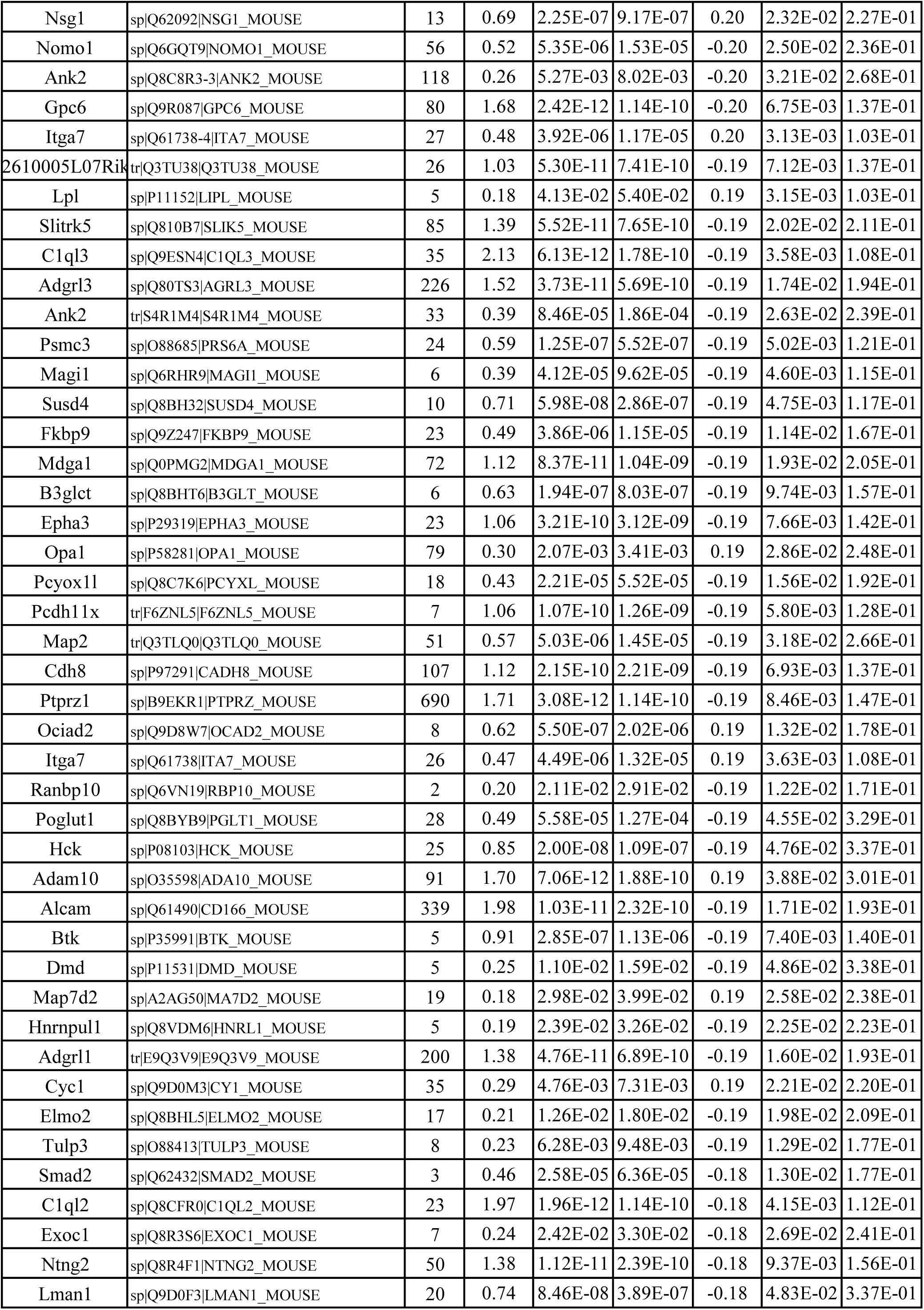
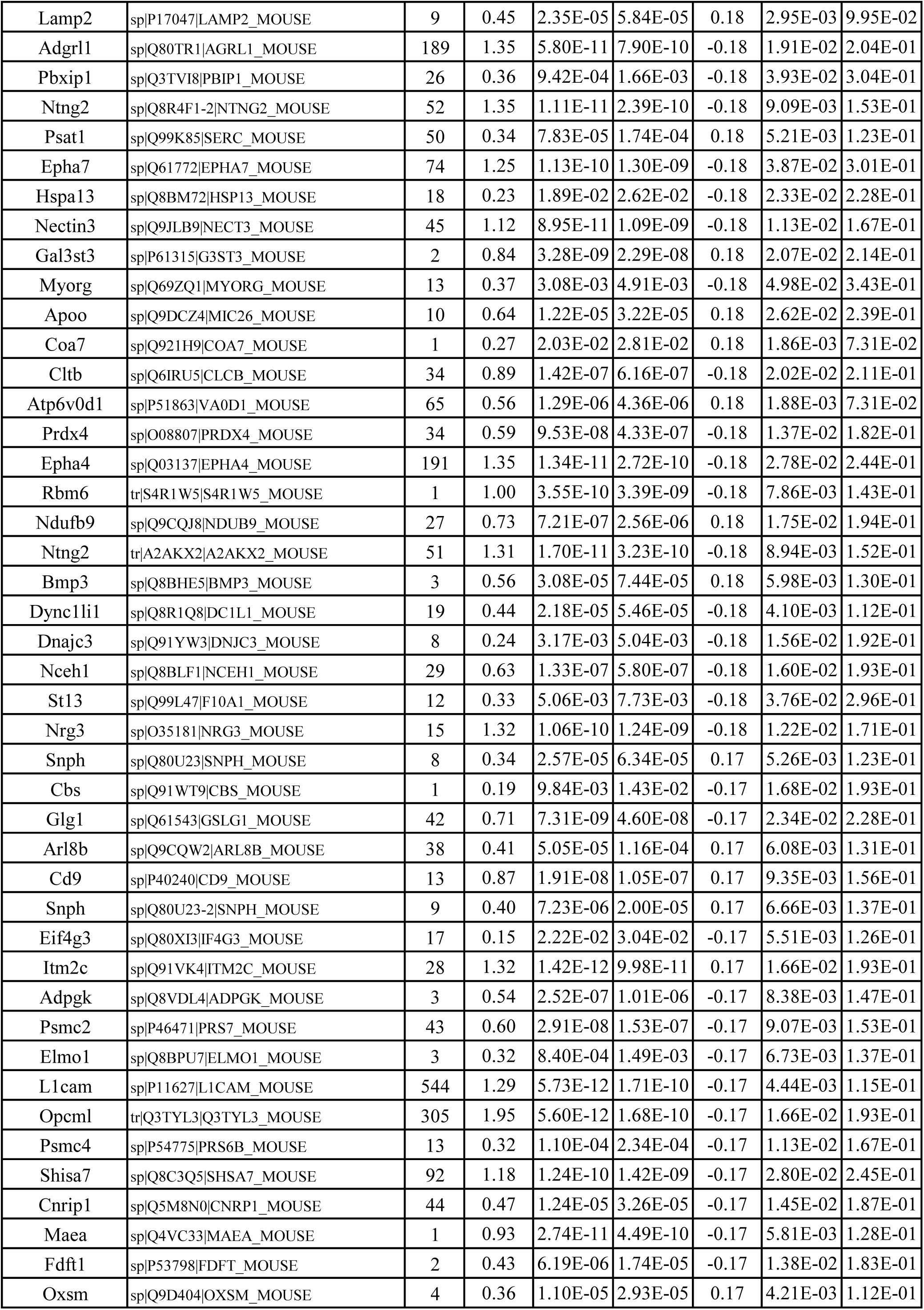
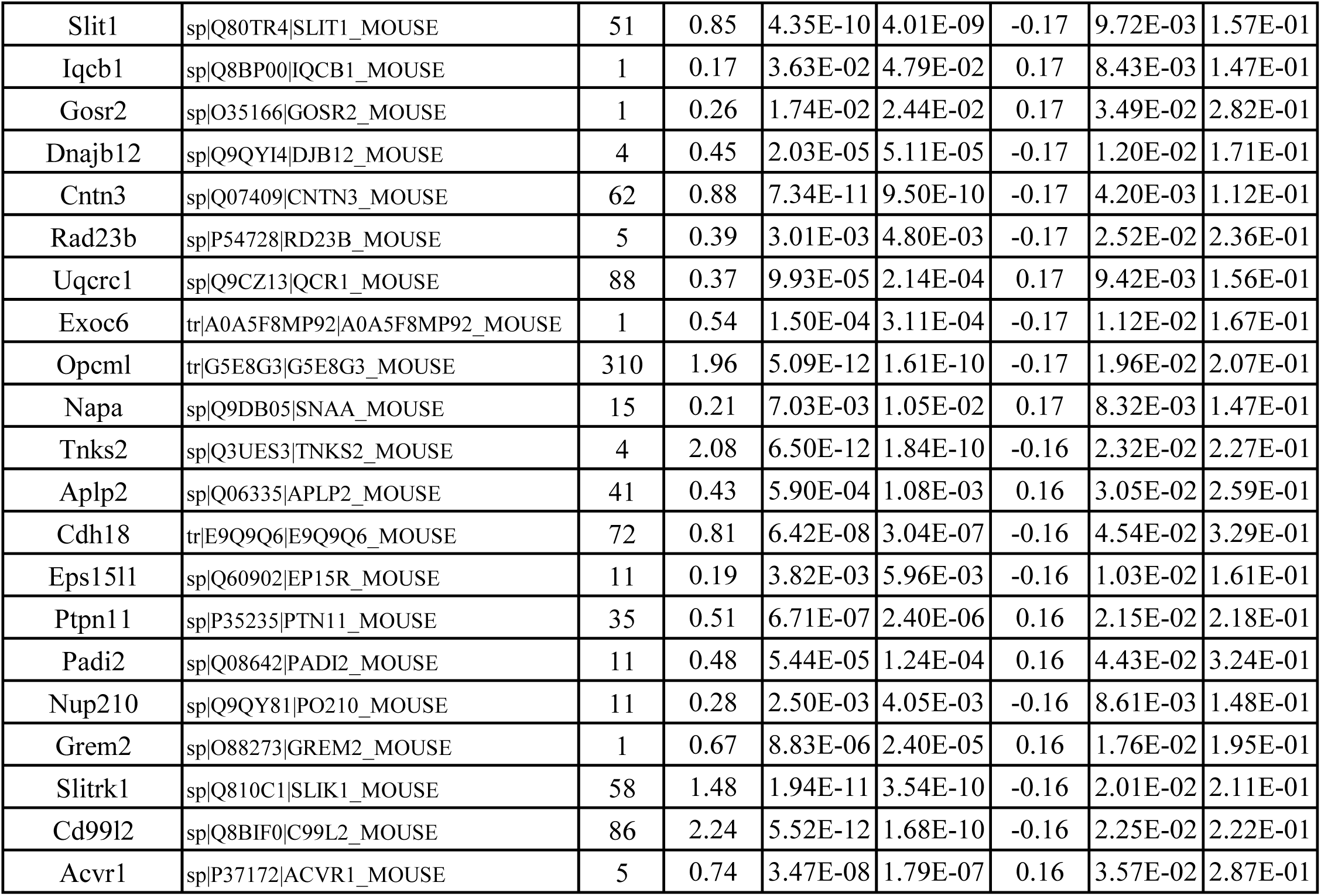
A list of 411 differentially expressed proteins (DEPs) in astrocytes comparing 5xFAD;HRP and HRP mice. DEPs were defined by enrichment in biotinylated samples (log2(HRP/Ctrl)>0, p<0.05) and significant disease-associated changes (log2(HRP;5xFAD/HRP)>1 SD [0.16], p<0.05).

**Table S2:**
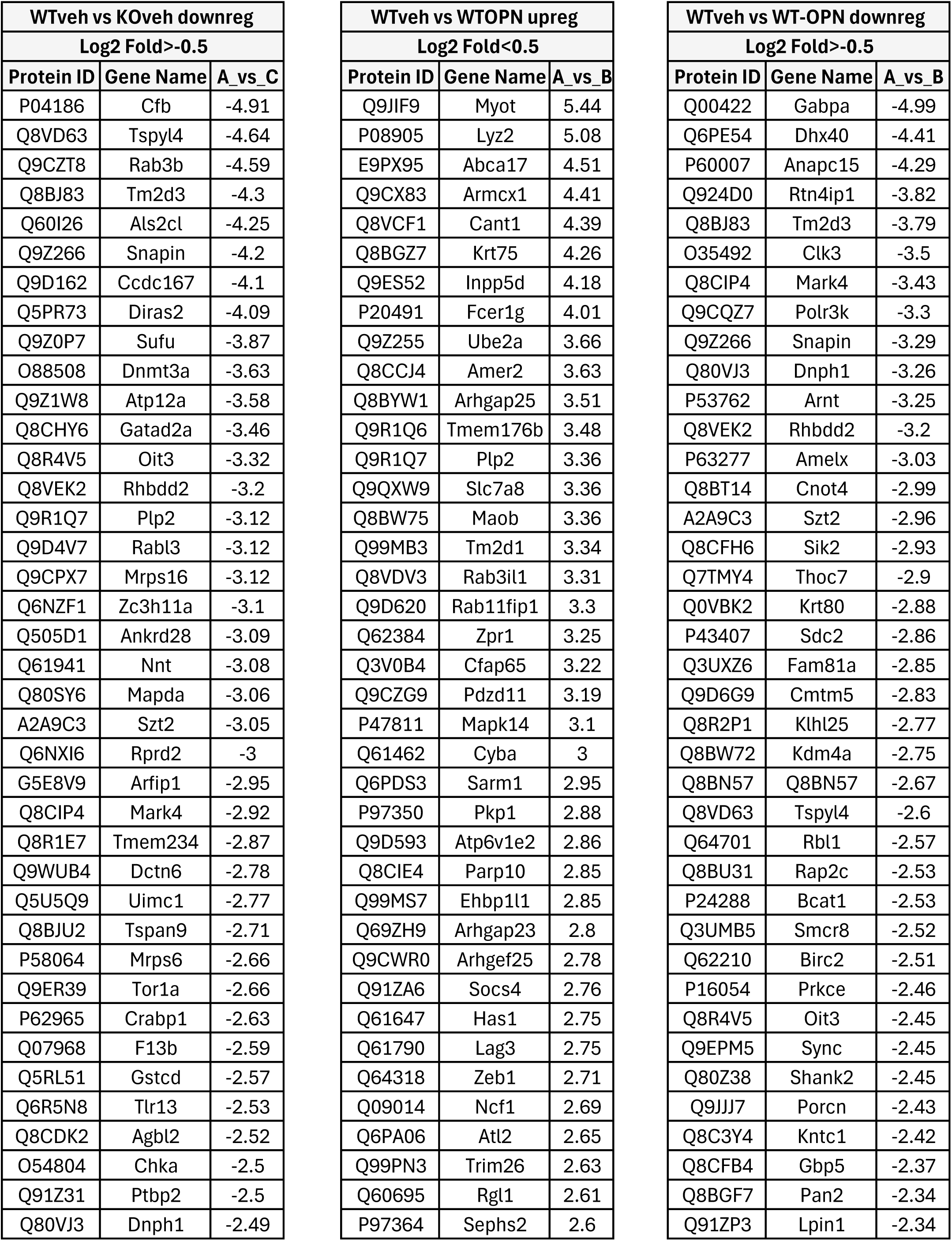
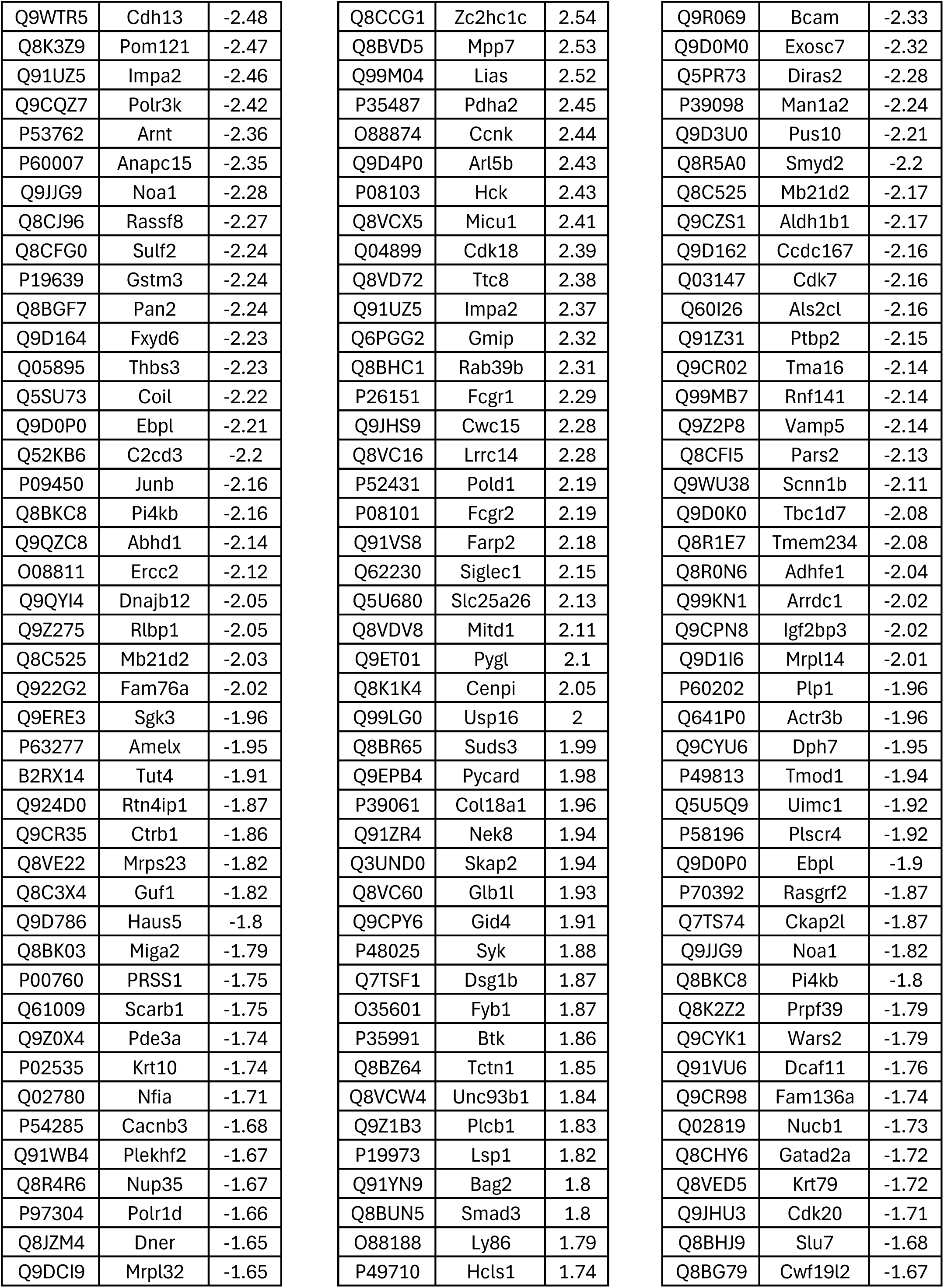
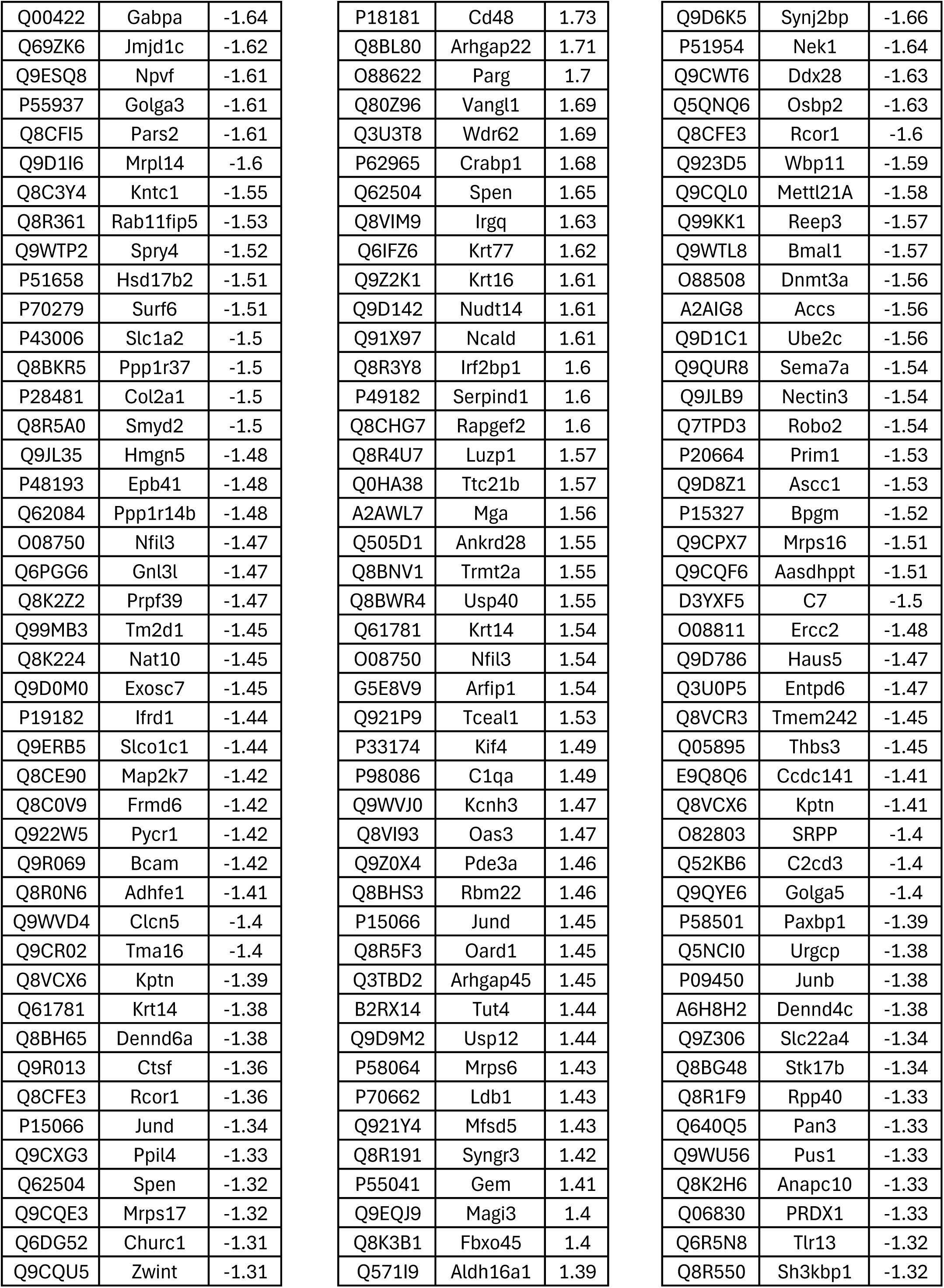
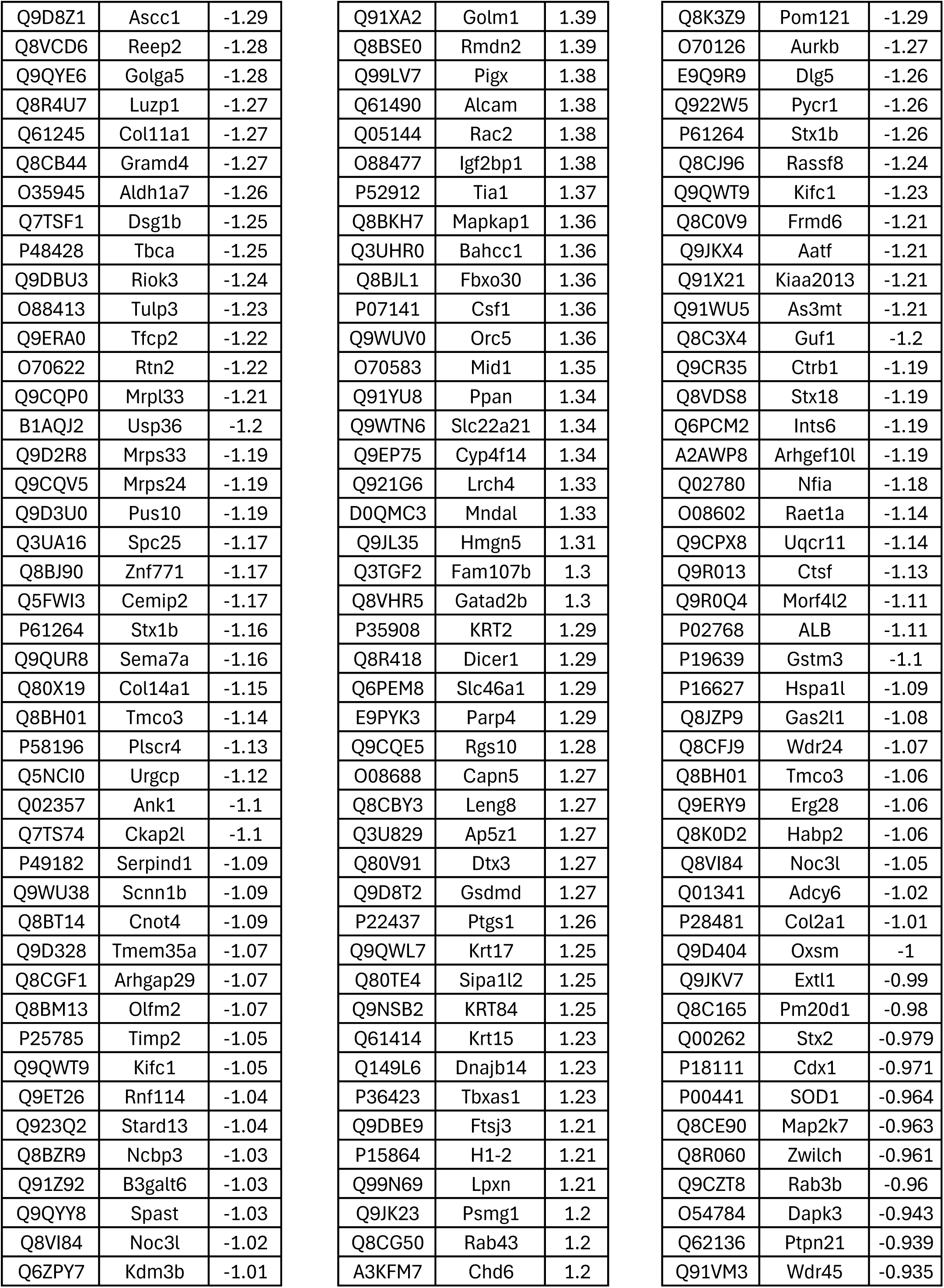
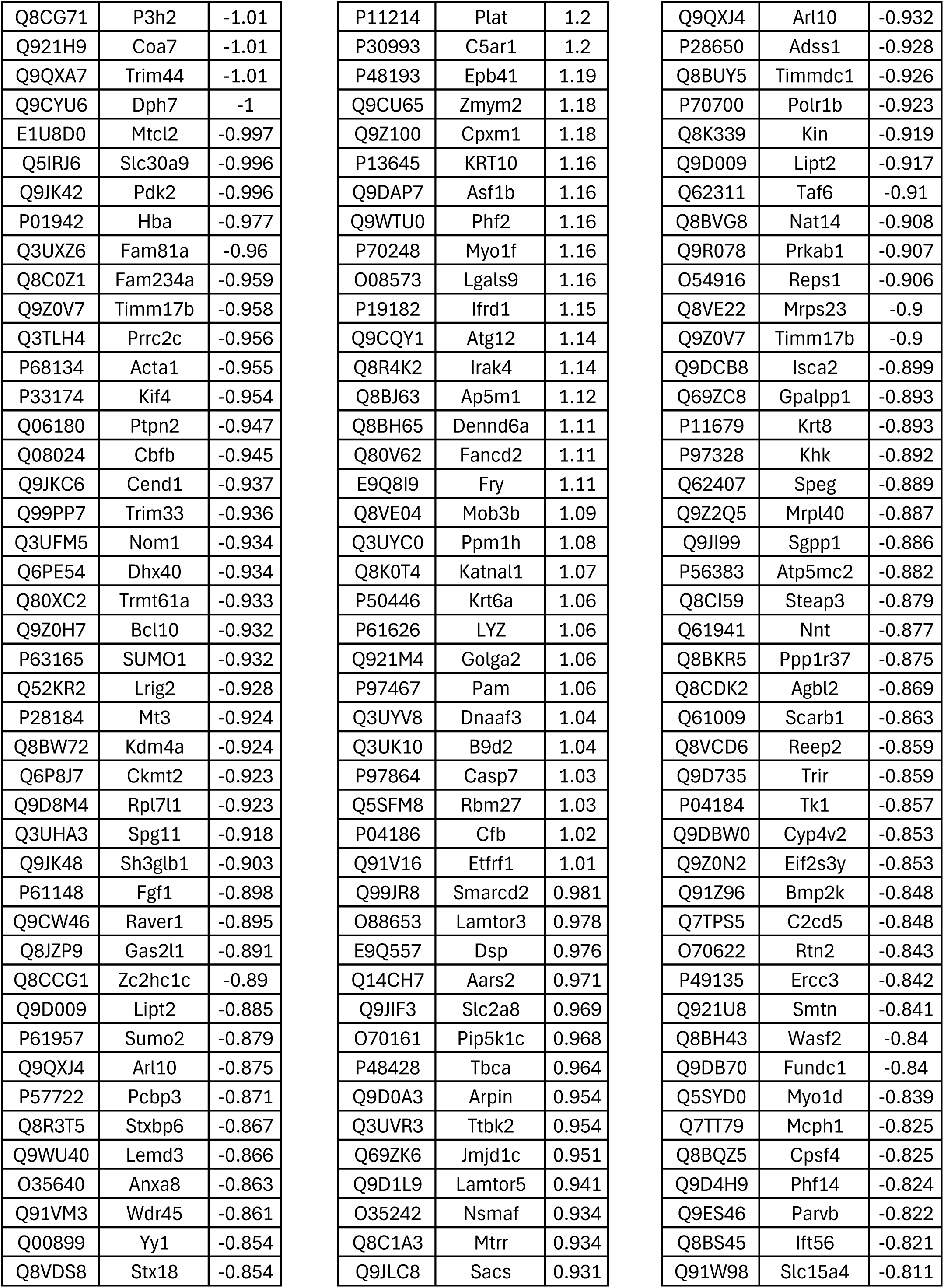
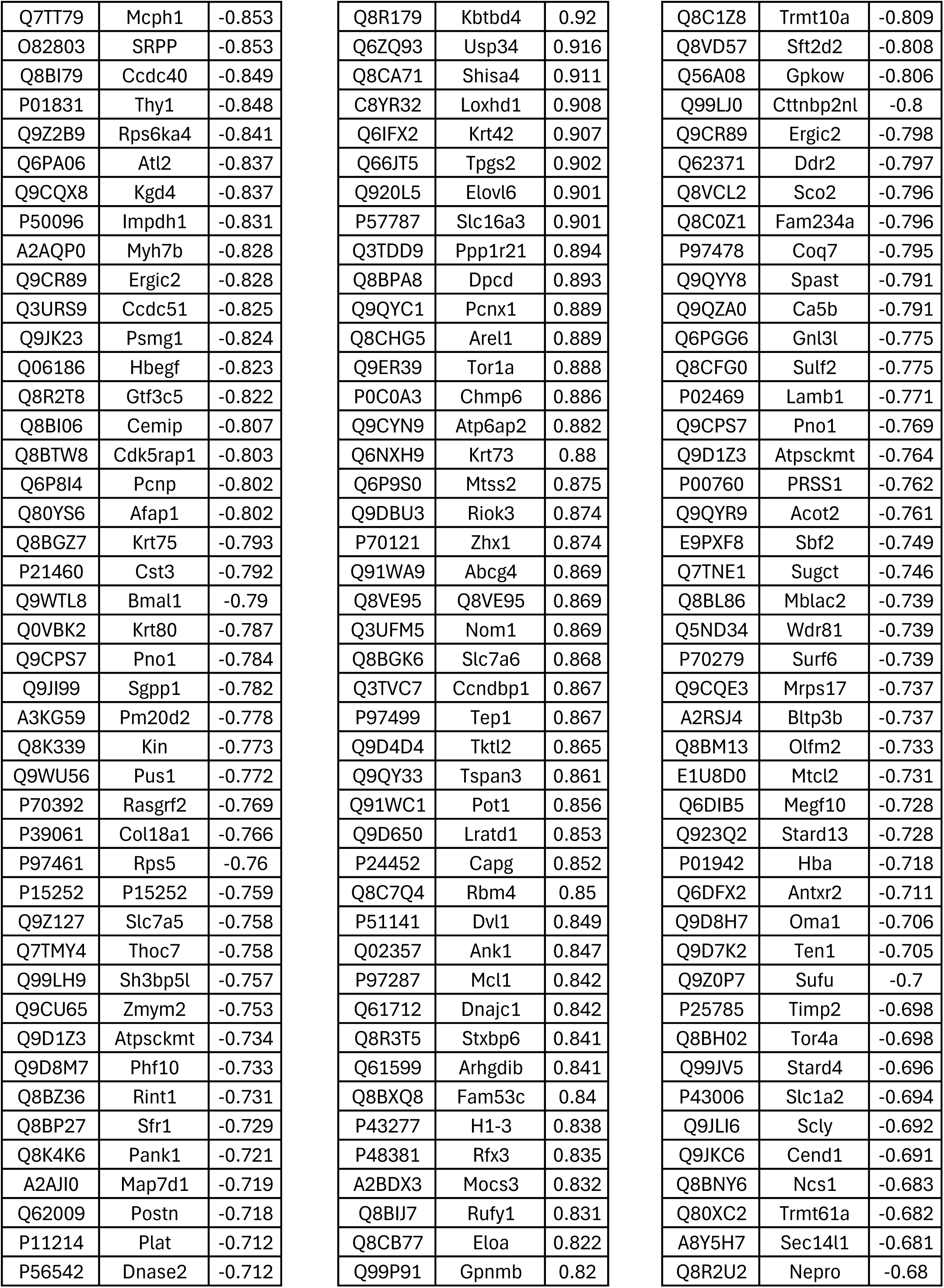
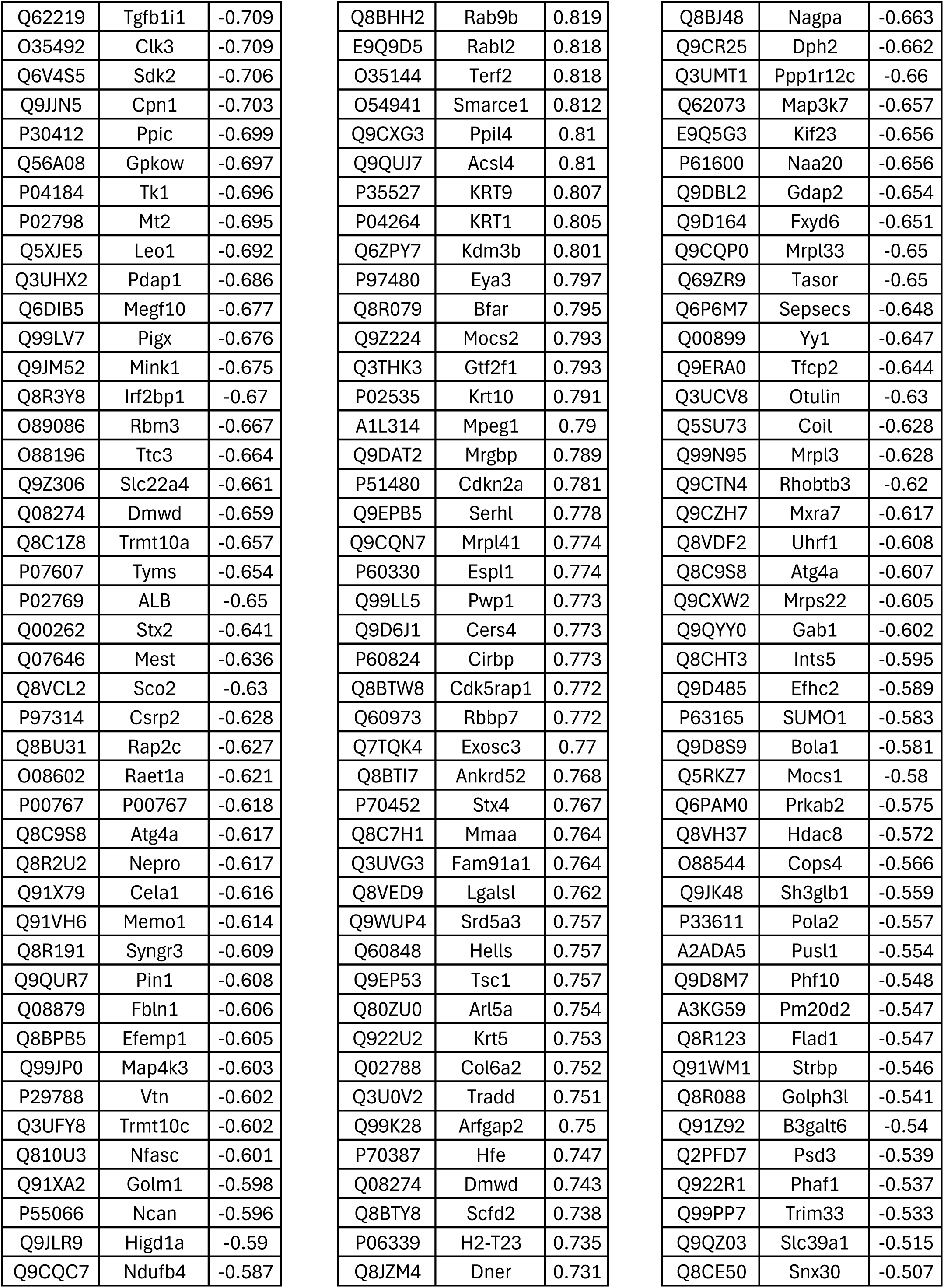
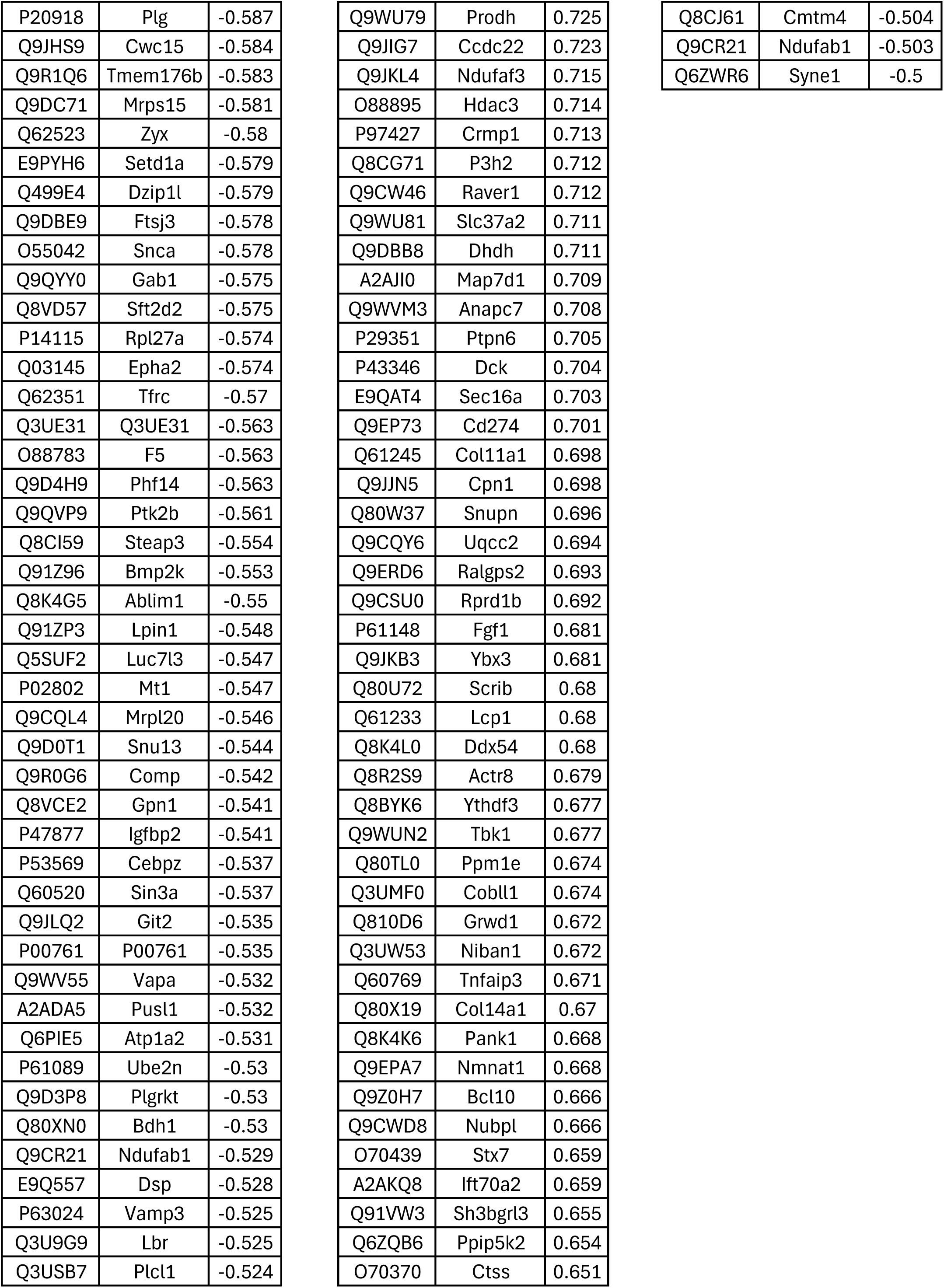
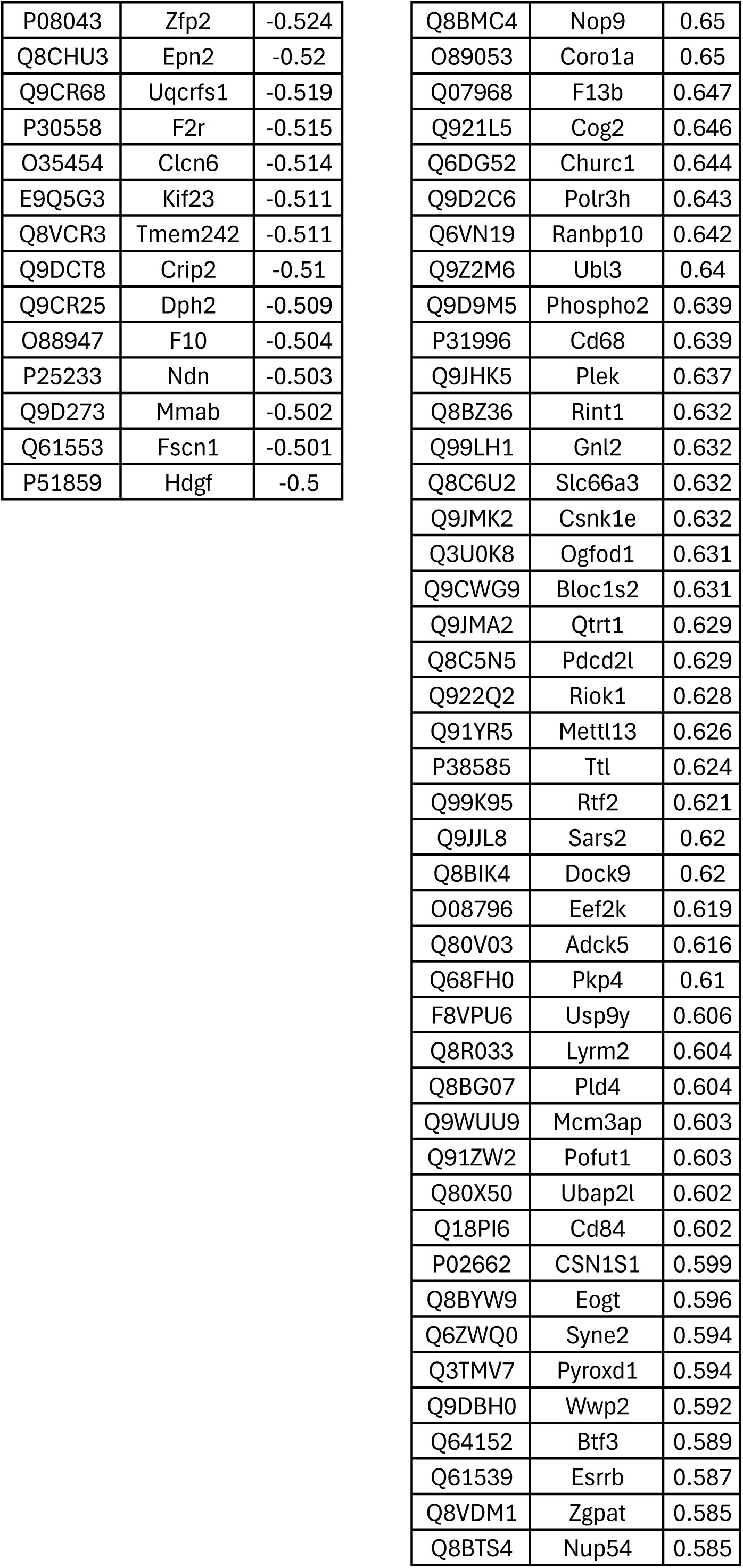
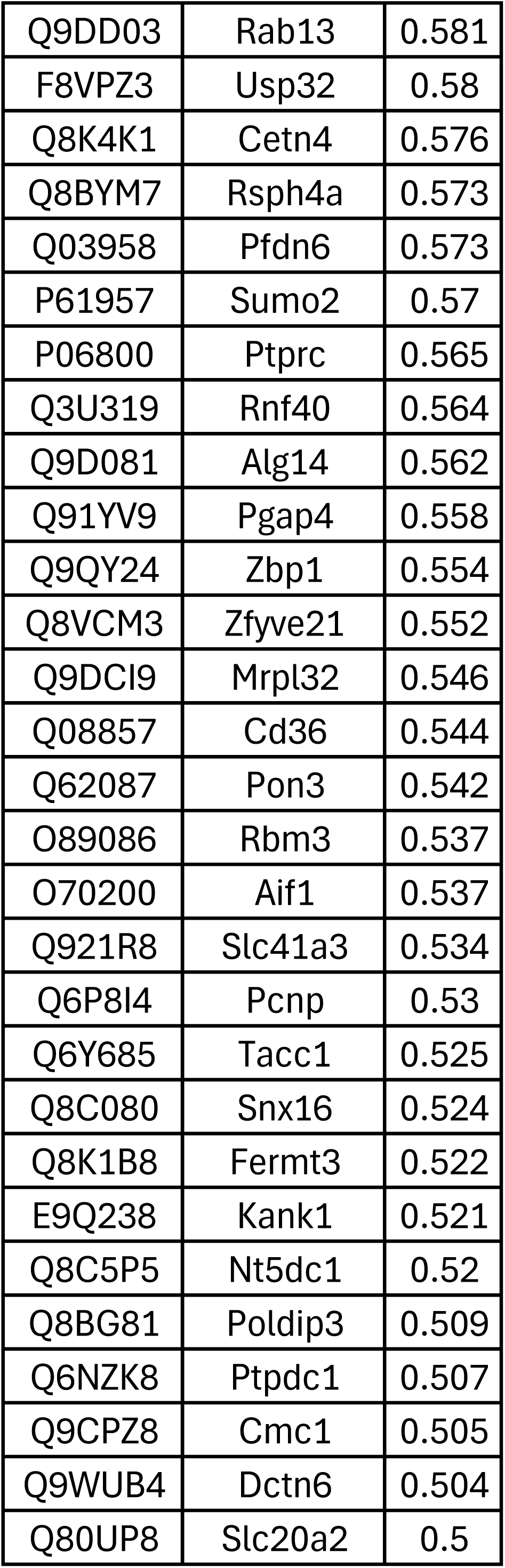
DEPs detected in WT and *Cd44*-KO astrocytes treated with vehicle or mrOPN.

**Table S3:**
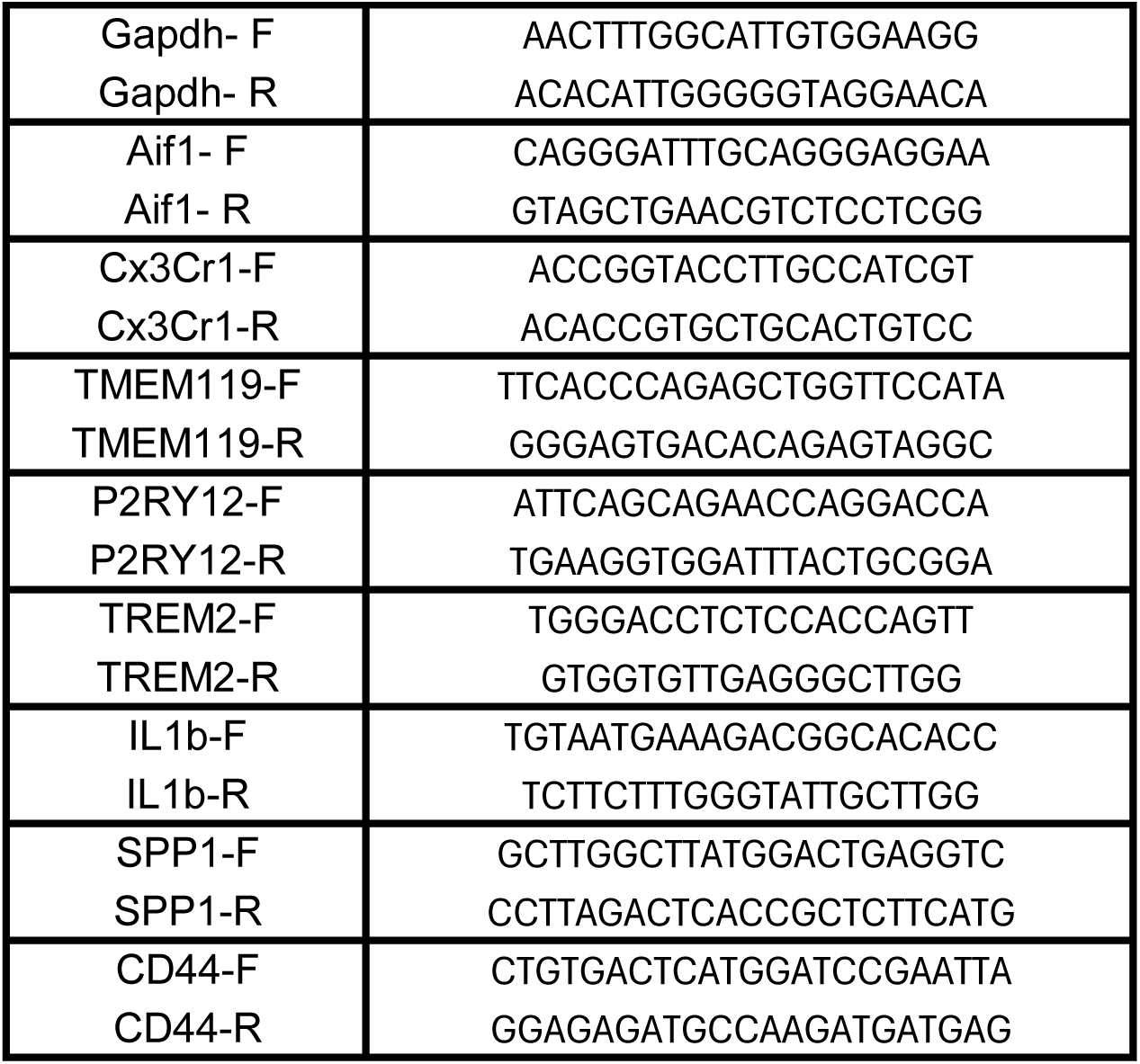
Primer Sequences

## Notes

https://massive.ucsd.edu

### Competing Interest Statement

The authors have declared no competing interest.

